# A Meta-analysis of Longevity Estimates of Mosquito Vectors of Disease

**DOI:** 10.1101/2022.05.30.494059

**Authors:** Ben Lambert, Ace North, H. Charles J. Godfray

## Abstract

Mosquitoes are responsible for more human deaths than any other animal, yet we still know relatively little about their ecology. Mosquito lifespan is a key determinant of the force of transmission for the diseases they vector, but the field experiments and dissection methods used to determine this quantity produce estimates with high uncertainty. In this paper, we use Bayesian hierarchical models to analyse a previously published database of 232 mark-release-recapture (MRR) experiments and two databases of different types of mosquito dissection experiments. One, compiled by us, consisted of 131 detailed estimates of the number of egg-laying (gonotrophic) cycles, the other, a recently published dataset of 1490 studies of parity (whether a mosquito has laid eggs or not) in anopheline malaria vectors. We analysed all studies with the same methodology and used Bayesian hierarchical statistics to obtain estimate at the species and genus level. For the major African malaria vector *Anopheles gambiae s*.*l*., we estimate lifespans ranging from 4.4 days (from MRR analysis) to 10.3 days (from parity analysis). For the predominantly East-African vector *An. funestus s*.*l*., our lifespan estimates range from 4.2 days (MRR) to 7.1 days (dichotomous parity analysis). We estimate lifespans ranging from 4.7 days (physiological age analysis) to 6.2 days (MRR) for *Aedes aegypti*, and a lifespan of 11.6 days for *Ae. Albopictus* (only present in MRR data) – the predominant vectors of key arboviruses. Additionally, we estimate that female mosquitoes outlive males by 1.2 days on average (mean estimate; 25%-75% CI: 0.3-1.6 days). By fitting a range of survival models to the data, we found relatively little evidence for senescence in the field. Our analyses, supplemented by power analyses, indicate the considerable uncertainty that remains about mosquito lifespan in the wild. We conclude further progress will require larger and longer experiments or the development of novel new methodologies.

**Author summary:** Mosquitoes transmit some of the most important diseases afflicting humans, with malaria alone killing between 390,000 and 460,000 people in 2019, chiefly children in low-income countries. The force of transmission of these diseases depends critically on the duration of mosquito lifespans, and some of the most successful disease control interventions, including insecticide-treated bednets, work because they reduce mosquito longevity. In this study, we conduct meta-analyses of two important classes of field experiments used to estimate wild mosquito lifespan: mark-release-recapture studies, where mosquitoes are marked with dye then released with the number of marked mosquitoes caught monitored over time; and experiments involving dissection of wild-caught females, whose reproductive anatomy is used as a biological clock to determine physiological age. In all analyses, we estimate that most mosquito species live less than 10 days on average, which suggests that relatively few mosquitoes live sufficiently long to transmit disease. The estimates obtained across the two field experiment types were largely discordant and indicated conflicting sources of heterogeneity in lifespan, likely due to the weak power of small-scale experiments. Finally, by fitting a range of survival models to the data, we conclude that, for most species, mosquitoes do not experience strong age-related increases in mortality. We critique the quality of the existing evidence base about mosquito lifespans in the field and suggest how it may be improved.

**Author contributions:** BL, AN and HCJG designed this study. BL was responsible for data curation, developing the statistical methodology and conducting the investigation. All authors were involved in drafting the original manuscript and revising it.

## Introduction

Some of the most important infectious diseases afflicting humans are transmitted by mosquitoes, including pathogens such as the causative agent of malaria that have been associated with humans throughout our evolutionary history (Carter and Mendis 2002), as well recently emergent infections, such as the Zika virus. Most mosquito species have a “gonotrophic cycle” involving successive episodes of vertebrate blood feeding, egg maturation and oviposition (Silver 2007). In order for a mosquito to transmit a pathogen it must feed on an infectious person and live long enough to complete at least one gonotrophic cycle and feed on an uninfected and susceptible individual. Adult lifespan is thus a critical determinant of the ability of a mosquito population to allow the persistence of an indirectly transmitted infection (Macdonald 1957). Lifespan can of course be straightforwardly assessed in the laboratory, but it is generally accepted that measurements under relatively benign laboratory conditions have limited relevance in the field, and much effort has been directed at estimating this parameter in the vector’s natural environment (Clements and Paterson 1981, Guerra, Reiner et al. 2014). Most work has focused on assessing average daily mortality rates, and the simplest assumption is that these do not vary with mosquito age – in this case, longevity is simply the reciprocal of mortality. Testing this assumption and discovering whether mosquitoes senesce or show other types of age-dependent mortality has also been studied in the field (Clements and Paterson 1981, Harrington, Vermeylen et al. 2014, Hugo, Jeffery et al. 2014).

There are two main strategies to estimate mosquito longevity. The first is through mark-release-recapture (MRR) experiments, a technique that is widely used to estimate lifespan in many types of animal. As applied to mosquitoes, insects are caught in the field or reared in the laboratory and then marked, typically with fluorescent dust. The mosquitoes are then released into the field and attempts made to recapture them, for example using human baits or light traps, usually over an extended period of time. Mortality rates can be statistically estimated from the numbers of recaptures given certain assumptions (Silver 2007). The main challenges with MRR is ensuring the marking technique does not affect recapture probability and distinguishing mortality from mosquitoes dispersing out of range of being recaptured. Also, releasing insects that can transmit disease (especially if this significantly increases ambient population levels) raises important ethical issues.

The second approach is specific to female mosquitoes and makes use of their gonotrophic cycle and involves two distinct dissection-based techniques. The simplest and most widely used approach is based on the observation that the appearance of the fine tracheoles encasing ovaries changes irreversibly when ovaries first develop (Detinova and Bertram 1962). The proportion of parous individuals – those individuals that have produced offspring – can be determined by dissecting field-caught specimens and, by making assumptions of the duration of gonotrophic cycles, an estimate of lifespan can be derived. In honour of the entomologist who first made this observation, this approach is known as Detinova’s method. The straightforward dissection technique needed to apply this method means it has been widely adopted, but the assumptions that need to be made limit the information that can be derived about mortality. The second technique requires more sophisticated dissection and involves counting the number of reproductive cycles a mosquito has undertaken. The mosquito ovary is made up of ovarioles, each of which typically produces one egg every gonotrophic cycle. After the egg passes into the oviduct, the distended ovariole does not completely recover its previous form, but a discrete dilation remains which can be detected by dissecting the female reproductive organs (Polovodova 1949). A skilled dissector can determine the number of such dilations, so providing richer data on longevity. This approach is known as Polovodova’s method after the scientist who first observed these changes. The challenges of this method include the amount of time and expertise it takes to collect data. There is also concern that ovarioles may not necessarily produce visible dilations during each oogenesis (Fox and Brust 1994) meaning that dissecting a given ovariole and counting its dilations may understate physiological age. This suggests that dissecting many ovarioles is required to obtain a good approximation of underlying age of a mosquito. Both dissection approaches are specific to females and require conversions between physiological and chronological time (though the distribution of the number of gonotrophic cycles wild-caught mosquitoes have gone through is of direct epidemiological relevance).

An issue with all methods is that they require logistically difficult and expensive field campaigns. There is thus value in conducting a meta-analysis of existing data to explore consistency across studies, to identify correlates of lifespan and to learn lessons for further studies. Here, we apply a common statistical methodology to analyse data from 232 MRR experiments, 1490 observations of parity obtained through Detinova’s method, and 131 studies that used Polovodova’s method to determine physiological lifespan. For both MRR and Detinova’s method, we make use of valuable published databases: for MRR, we use that published by (Guerra, Reiner et al. 2014); for Detinova’s parity determination, we use a study of anopheline malaria vectors assembled by Massey et al. (2016). In addition, we extracted data from studies that used Polovodova’s method ourselves via a literature search. We concentrated on the three major genera of mosquito vectors, *Anopheles, Aedes* (in its traditional sense) and *Culex*, which constitute the majority of the data.

## Glossary

- MRR experiments – mark-release-recapture experiments, where mosquitoes are marked with a dye or equivalent, released and then subject to potential recapture.
- Gonotrophic cycle – the sequence of searching for a host, blood-feeding, egg maturation and oviposition for a female mosquito.
- Parity rate – the proportion of female mosquitoes that have laid eggs.
- Nulliparous – a female that has not laid eggs.
- Parous – a female that has laid eggs.
- Uniparous / biparous / triparous – a female that has undergone 1 / 2 / 3 gonotrophic cycles.
- Physiological or reproductive age / time – the number of gonotrophic cycles a female has undergone throughout their life / over a period of time.
- Chronological age / time – age or time measured in calendar time (e.g. days).
- Detinova’s (dissection) method – dissecting mosquitoes to determine whether a female is nulliparous or parous (i.e. providing a dichotomous measure of reproductive status).
- Polovodova’s (dissection) method – dissecting female mosquitoes to determine the physiological age.

## Results

We report estimates of the lifespan of different mosquito species (the mean unless otherwise stated) obtained using the techniques described in the Introduction. We used a Bayesian approach to parameter estimation which provides posterior distributions describing uncertainty in the lifespan estimates. In the Supplementary Online Material (SOM), we provide detailed quantiles and summary measures but here report only the posterior median of mean lifespan with 25%-75% central posterior intervals given as uncertainty measures.

### Most estimates of mosquito lifespan from MRR studies are less than ten days, though there is considerable variation

We first estimated lifespan independently for each available MRR time-series (Fig. 1; Methods). The estimates varied substantially both within and among species, though a majority were less than ten days (187 of 236 time-series point estimates). In comparison, mosquito longevity in laboratory conditions is typically found to exceed 30 days (Styer, Carey et al. 2007). Our estimates ranged from 0.7 days from a study of the predominantly Australasian *Anopheles annulipes* to 38.3 days from a study of *Aedes aegypti*. It is likely that the very short longevity estimates reflect dispersal out of the recapture zone or a violation of the assumptions of our analyses (see Discussion). There are multiple data sets for the most important vector species such as *An. gambiae s*.*l*. (malaria), *Ae. aegypti* and *Ae. albopictus* (yellow fever, dengue and Zika viruses) and *Culex tarsalis* (West Nile Fever, Western Encephalitis), all of which show considerable variation. For example, there were 54 estimates of lifespan for *Ae. aegypti* which range from 2.2 days to 38.3 days with a mean of 8.3 days and coefficient of variation of 0.7.

**Figure 1:**
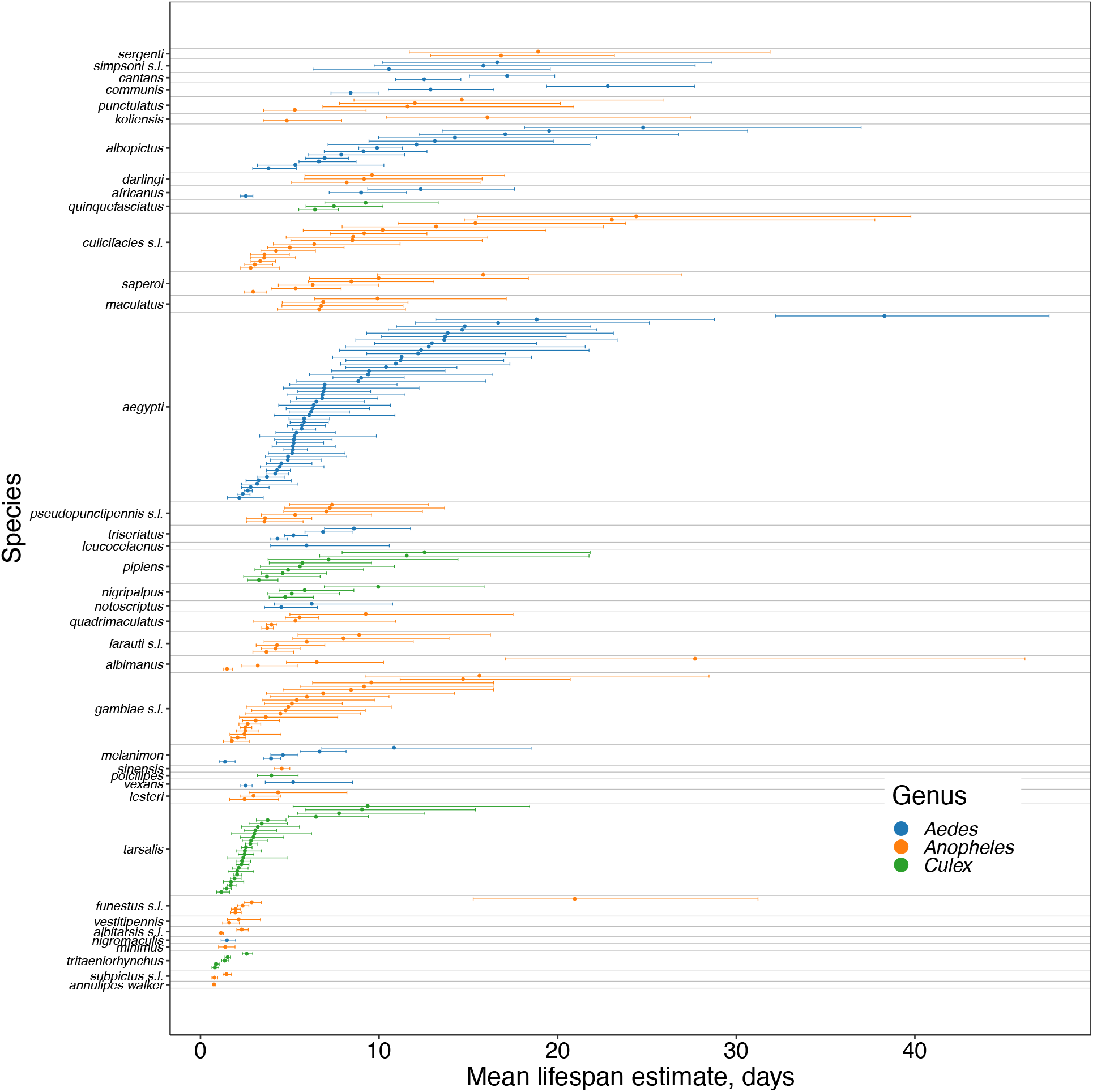
MRR: lifespan estimates for each time series. The middle point shows the median estimates, and the left and right box whiskers show the 25% and 75% posterior quantiles respectively. All estimates were obtained using the non-hierarchical exponential survival model as described in text. The individual estimates within each species are ordered according to the median lifespan estimates; the species estimates are ordered corresponding to the species-level medians.

### MRR data show more evidence of lifespan variation at the genus than species level

We next combine the data from individual studies using a suite of Bayesian hierarchical models. The first model pools studies by species, the next by genus, and a final model pools all studies together. To ensure comparability, in this analysis we exclude studies where females were blood or sugar fed before release (Fig. 2 and Table S1) (below we investigate the role of feeding prior to release). We also set a threshold of at least two time series (i.e. two separate MRR studies were required) for inclusion in the species level analysis, which is why some species present in Fig. 1, for example *An. annulipes*, are not included in Fig. 2. The most long-lived species was *Ae. simpsoni s*.*l*. (an African vector of yellow fever), and the most short-lived was *An. subpictus s*.*l*. (an Asian malaria vector), although this estimate was unfeasibly low and almost certainly reflects dispersal out of the recapture zone or a violation of the assumptions of our analyses. There were also differences in longevity at the genus level, with *Culex* estimated to have the shortest lifespan and *Aedes* the longest.

**Figure 2:**
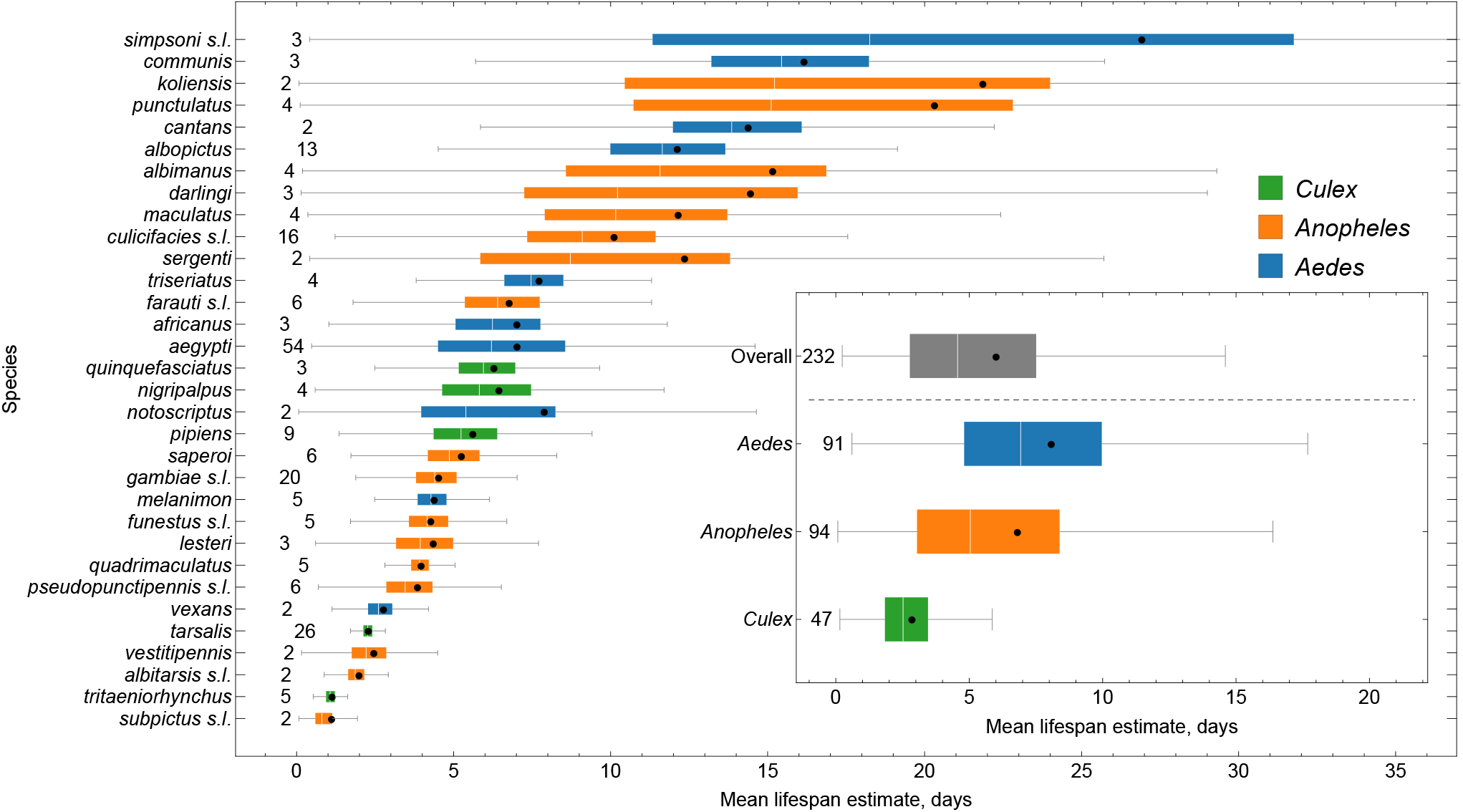
MRR: grouped lifespan estimates. The lifespans shown are for mosquitoes that were not fed with sugar or blood before release. The middle line in each box shows the median estimates and the solid dot indicates the mean. The left and right box edges show the 25%, and 75% posterior quantiles respectively. The whiskers show the range of the data, excluding points lying more than 1.5 times the interquartile range away from each edge of the box. The numbers before the start of the left whisker indicate the number of individual time-series within each species. All estimates were obtained using the hierarchical exponential survival model.

The considerable variation in these results probably stems from a combination of biological differences at each of the levels in the hierarchy and idiosyncratic study-level variation. To better understand the relative roles of these factors, we compared the fit of the different hierarchical models. Since models with more parameters will naturally fit the data more closely, we used a cross-validation approach that explicitly accounts for this when comparing models (see SOM). The genus level model fitted the data significantly better than the overall model, yet the species model did not significantly improve the fit over the genus model (Table S1). This suggests that there are consistent differences in lifespan across the three genera, yet within each genus there are no consistent differences among species.

### MRR data suggest females tend to live longer than males

The MRR studies included male-only and female-only releases, and mixed releases of both sexes, allowing us to estimate male and female lifespan at the genus level (Fig. 3). Each genus showed a trend for females to live longer than males, with the greatest difference for *Aedes* (2.5 days; *p*<0.01, where *p* is the fraction of pairwise posterior samples in which males outlive females), followed by *Anopheles* (2.0 days; *p*=0.17) and *Culex* (0.3 days; *p*=0.34). Overall, female mosquitoes were estimated to live 0.9 days longer than males (*p*=0.10). It is possible that these differences in lifespan are due to sex-specific variation in dispersal, but, to our knowledge, there is limited evidence to suggest this.

**Figure 3:**
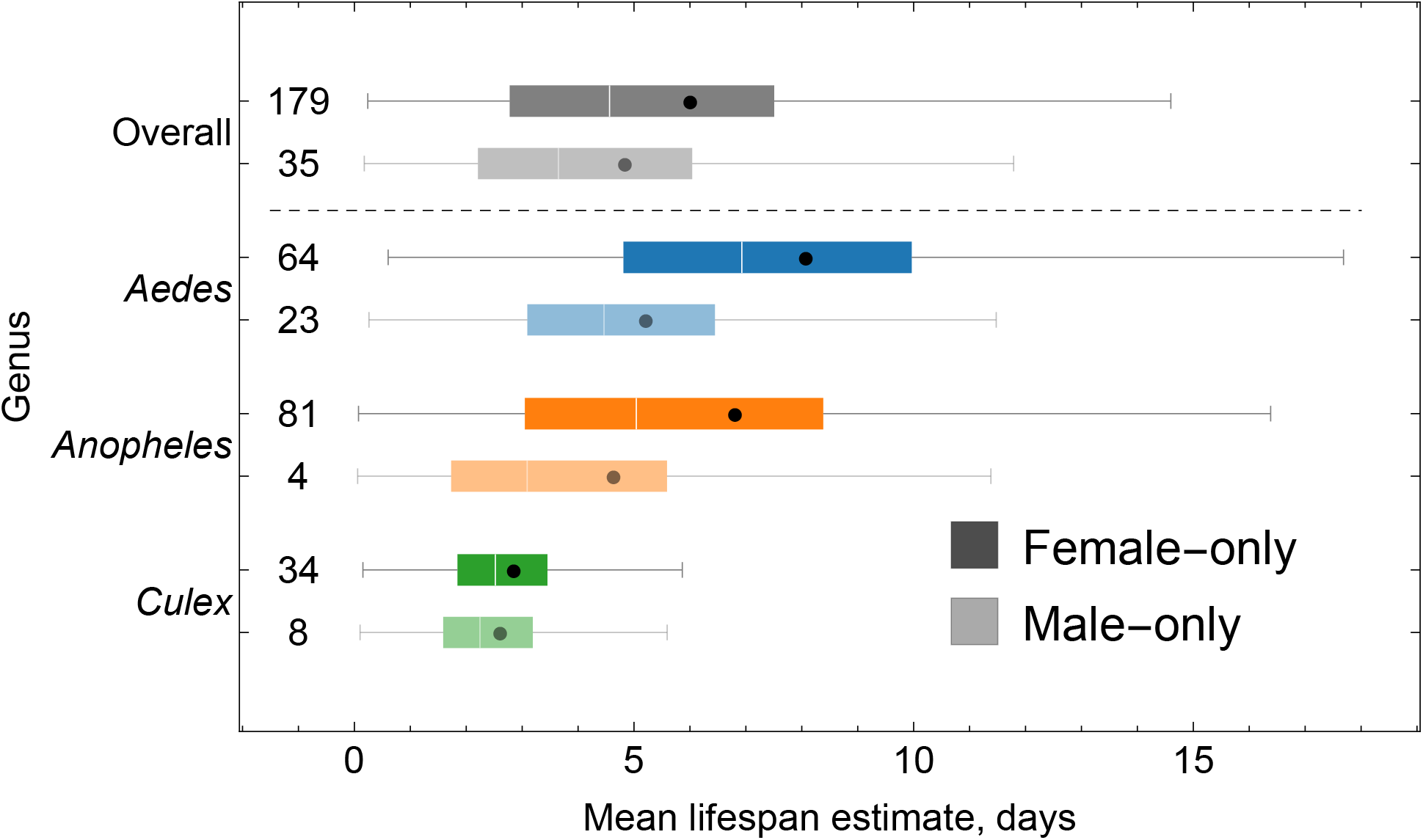
MRR: lifespan estimates by sex. The lifespans shown are for mosquitoes that were not fed with sugar or blood (for females) before release. The middle line in each box shows the median estimates and the solid dot indicates the mean. The left and right box edges show the 25% and 75% posterior quantiles respectively. The whiskers show the range of the data, excluding points lying more than 1.5 times the interquartile range away from each edge of the box. The numbers before the start of the left whisker indicate the number of individual time-series within each species. All estimates were obtained using the hierarchical exponential survival model.

### MRR data suggest that sugar feeding modestly increases the lifespan of marked mosquitoes, yet blood-feeding of females makes little difference

MRR experiments differ in whether mosquitoes are fed or not before their release, and if so whether with sugar, blood, or both. We studied the effects of feeding on female lifespan at the genus level and across all studies (Fig. S1). Since there were insufficient data at the genus level on males that were fed with sugar versus unfed, we combined all genera to estimate a pooled effect of sugar-feeding. For both sexes and across all genera, we found modest evidence that sugar feeding prior to release extended life-expectancy, by an average of 0.6 days for females (*p*=0.15, where *p* is the fraction of pairwise posterior samples in which unfed outlived fed) and 0.5 days for males (*p*=0.15). The effect of blood-feeding on female mosquitoes was not significant, with blood-fed individuals living about 0.1 days longer than unfed mosquitoes (*p*=0.44).

### Lifespan estimates from MRR data were not affected by spatial scale of the experiment as would be expected if dispersal out of the study area was causing severe bias

Following a release of marked mosquitoes, the rate of their recapture typically reduces over time because some mosquitoes die and because some disperse out of the recapture area. These factors are indistinguishable in spatially averaged recapture data, which is why our estimates are lower bounds on lifespan. If dispersal out of the recapture area commonly reduces the lifespan estimate below the true lifespan then we should expect a positive correlation between the spatial extent of the recapture zone and lifespan. This pattern was not apparent in our data, suggesting that dispersal may not be responsible for severe underestimation (Fig. S2). However, we suspect that dispersal bias may have affected a number of individual studies where we obtained unfeasibly short lifespan estimates, for example the case of *An. annulipes* noted above.

### Detinova dissection data show that the use of insecticides reduces anopheline longevity

The Massey et al. (2016) database contained 1490 parity observations across 26 anopheline species-complexes (henceforth ‘Detinova data’). Insecticide based control measures were in use at the time and place of the study in 519 cases, not in use in 364 cases, and this was unspecified in the remaining 607 cases. Since insecticides act by killing mosquitoes, their use should reduce mean lifespan, and we investigated whether this was detectable in the data. We analysed 867 observations (509 with insecticide and 358 without) representing all data from the 16 species complexes with 5 or more observations (see SOM). The effect of insecticides was large in all cases (Fig. S3), on average reducing lifespan by 56% at the species level (51%-58%; difference in posterior median estimates of mean lifespan).

In the following analyses of the Detinova data, we omitted those cases where insecticide was known to be in use, leaving 1126 observations. It is possible that insecticides were used in some of the 607 cases where this was not recorded. However, 76% of these studies took place before 2000, when bednets impregnated with pyrethroid insecticides began to be widely distributed across the African continent (Bhatt et al., 2015; elsewhere, their distribution remained low) suggesting that the majority of these results will be unaffected by insecticide.

### Detinova dissection data suggest most female anopheline mosquitoes complete fewer than three gonotrophic cycles, though this varies considerably between species

Detinova dissection determines parity, which indicates the fraction of female mosquitoes that have laid eggs (i.e. gone through one gonotrophic cycle). Assuming equal-length cycles throughout a mosquito’s life, we can estimate (see SOM) the average number of gonotrophic cycles (physiological lifespan) completed by the mosquitoes sampled in each study (Fig. S4). In 78% of cases, fewer than three gonotrophic cycles were completed before death. We next combine these data into Bayesian hierarchical models which pool the studies by species, then species-complex, then continent (here, including Africa, Asia, and Americas), and finally all the studies are combined together. The lifespan estimates, now in terms of gonotrophic cycles rather than chronological time, are shown in Fig. 4 and Table S2. Two of the species with the lowest estimated lifespans belong to the *An. albitarsis* species complex (Sp. A & Sp. B), which is a malaria vector found throughout South America. The taxon with the greatest lifespan was *An. albitarsis marajoara*, which is part of the same complex (formerly Sp. C), indicating extensive variation in this complex across the continent. Outside of the Americas, the species with the longest estimated lifespans were the major African vector *An. funestus*, and, in Asia, *An. acontinus*. Across the species complexes, the shortest lifespans were estimated for *An. aquasalis s*.*l*., which is a dominant malaria vector species in the Amazon (Sinka, Bangs et al. 2010), and *An. gambiae s*.*l*.. The anopheline species in Africa were estimated to live longest, followed by those in Asia and then the Americas.

**Figure 4:**
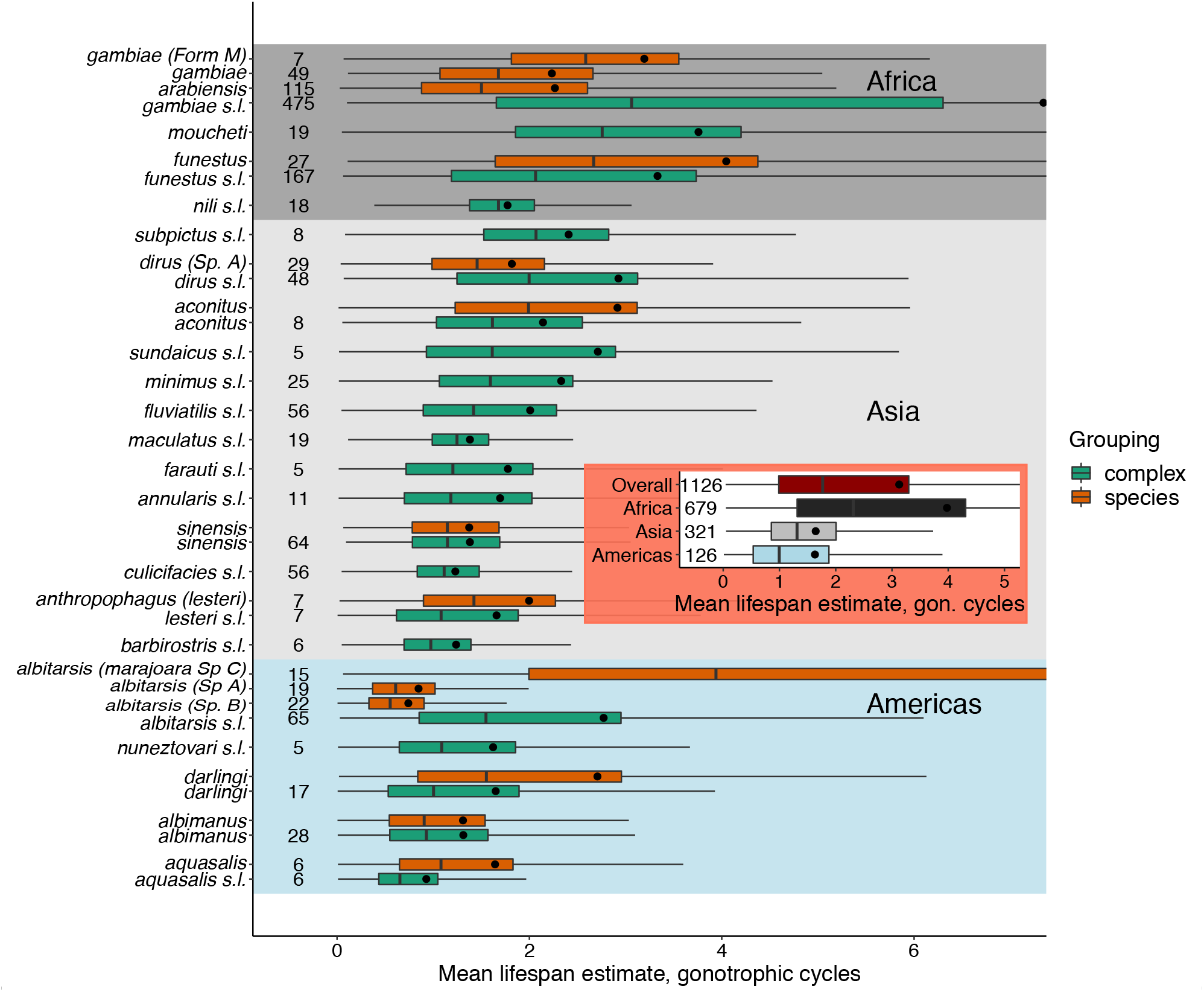
Detinova dissection: grouped lifespan estimates for anophelines. As described in SOM, both species and morphospecies estimates are provided where available. The three coloured backgrounds show data according to continent where experiment was carried out (Americas-blue, Asia-grey, Africa-black). The middle line in each box shows the median estimates and the solid dot indicates the mean. The left and right box edges show the 25%, and 75% posterior quantiles respectively. The whiskers show the range of the data, excluding points lying more than 1.5 times the interquartile range away from each edge of the box. The numbers before the start of the left whisker indicate the number of individual time-series within each species.

To help understand these results, we compared the fit of our hierarchical models at different taxonomic levels in a similar manner to our analysis of variance in the MRR data. Across all continents, the species-level model had the best fit followed by the species-complex level model then the continent-level model (Table S1). Introducing grouping at the species-complex level led to similar gains in the estimated fit over the continent-level model as introducing species did over the species-level model. This suggests that there are consistent differences in lifespan between the species-complexes in our analysis, and also amongst the species within these complexes. This differs from our MRR results, a matter we will return to in the Discussion.

### Polovodova dissection data show no variation in lifespan by species or genus

We estimated lifespan from 131 studies reporting Polovodova data (Fig. 5 and Table S3), again using Bayesian hierarchical models to pool information across studies. This data included species in four genera (*Anopheles, Mansonia, Culex*, and *Aedes*), and we thus used species, then genus, and finally all species as the hierarchy levels.

**Figure 5:**
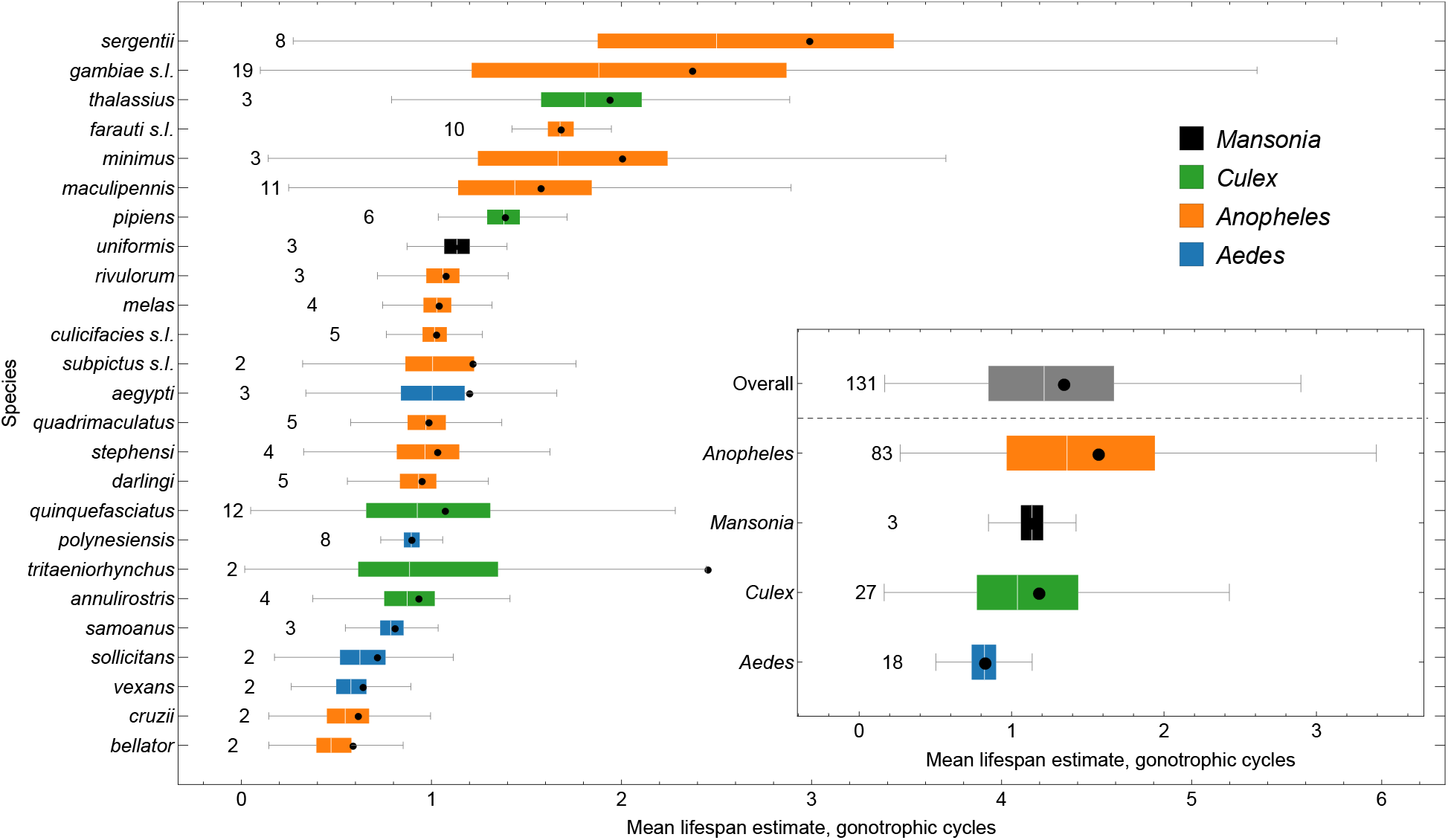
Polovodova dissection: grouped physiological lifespan estimates. The middle line in each box shows the median estimates and the solid dot indicates the mean. The left and right box edges show the 25% and 75\% posterior quantiles respectively. The whiskers show the range of the data, excluding points lying more than 1.5 times the interquartile range away from each edge of the box. The numbers before the start of the left whisker indicate the number of individual time-series within each species. All estimates were obtained using the hierarchical exponential survival model.

The mean number of cycles completed in a lifetime was estimated to be 1.2, and ranged from 0.45 for *An. bellator*, which transmits malaria in Brazil’s Atlantic Forest, to 2.45 for *An. sergenti*, which is adapted to desert conditions (it is known as the “oasis vector” of malaria; (Sinka, Bangs et al. 2010)) possibly accounting for its greater longevity.

At the genus level, *Anopheles* were estimated to go through the most gonotrophic cycles (1.4) followed by *Mansonia* (1.1), *Culex* (1.0), and Aedes (0.8).

As was done for the MRR and Detinova dissection data, we compared the fit of the models at each taxonomic grouping (species-level, genus-level and all data pooled) using a cross-validation approach. In contrast to the other two datasets, the models grouped at each of the three levels all fit the data equally well (Table S1).

### There is evidence that gonotrophic cycle duration varies across mosquito genera

To convert reproductive lifespan estimates into calendar time, it is necessary to estimate the duration of the gonotrophic cycle. We conducted a literature survey of gonotrophic cycle duration measurements, yielding 45 estimates based on laboratory observations and 36 estimates based on MRR methods. In the latter studies, an estimate of the gonotrophic cycle duration was obtained by releasing mosquitoes of known age and examining the parity of recaptured individuals. In 20 cases, separate estimates for the first gonotrophic cycle were reported which we included in our analysis. We combined these data using a regression approach which modelled the individual estimates, including any reported uncertainties, as quantiles of a normal distribution (SOM).

We found *Culex* mosquitoes to have the longest first gonotrophic cycles (with a mean of 5.2 days; Fig. 6), followed by *Aedes* (4.5 days) then *Anopheles* (3.7 days). These differences among genera were significant (ANOVA: F_2,116_ = 8.7, p<0.01), and the genus ordering was maintained for subsequent cycle durations. After pooling data across all genera, the first cycle duration was estimated to take 4.0 days, and subsequent cycles were estimated to take 3.6 days.

**Figure 6:**
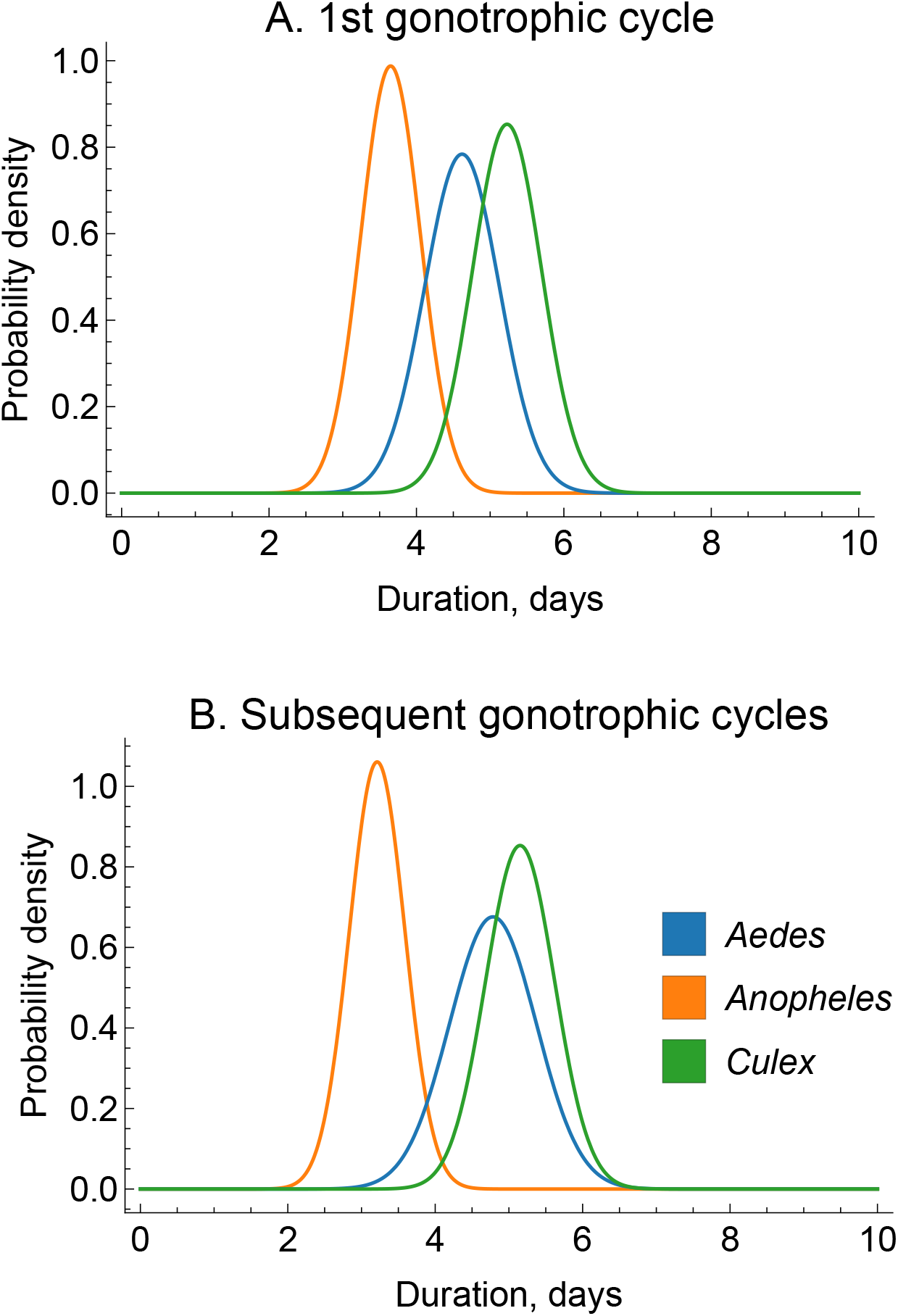
Estimates of gonotrophic cycle duration. Coloured lines represent different genera, with each line indicating the estimated Gaussian probability density representing uncertainty over the reproductive cycle length.

### There are no significant correlations in species-level lifespan estimates using the three different methodologies

We used our estimates of gonotrophic cycle durations to convert the dissection-based estimates of reproductive lifespan into chronological lifespan as described in the SOM (Tables S4 and S5 provide summaries of chronological lifespan estimates for the species and genera in the Detinova and Polovodova analyses). This allowed us to compare the lifespan estimates from the dissection studies and the MRR studies, where data from the same species were available (Fig. 7). Comparing the two dissection methods (Fig. 7A), there was a positive correlation across six species. For the ten species with both Detinova and MRR estimates, there was a slight negative correlation (Fig. 7B), while there was a positive correlation between the Polovodova estimates and those from MRR (Fig. 7C; n=12). In no cases were these correlations statistically significant, though the sample size for each comparison is small.

**Figure 7:**
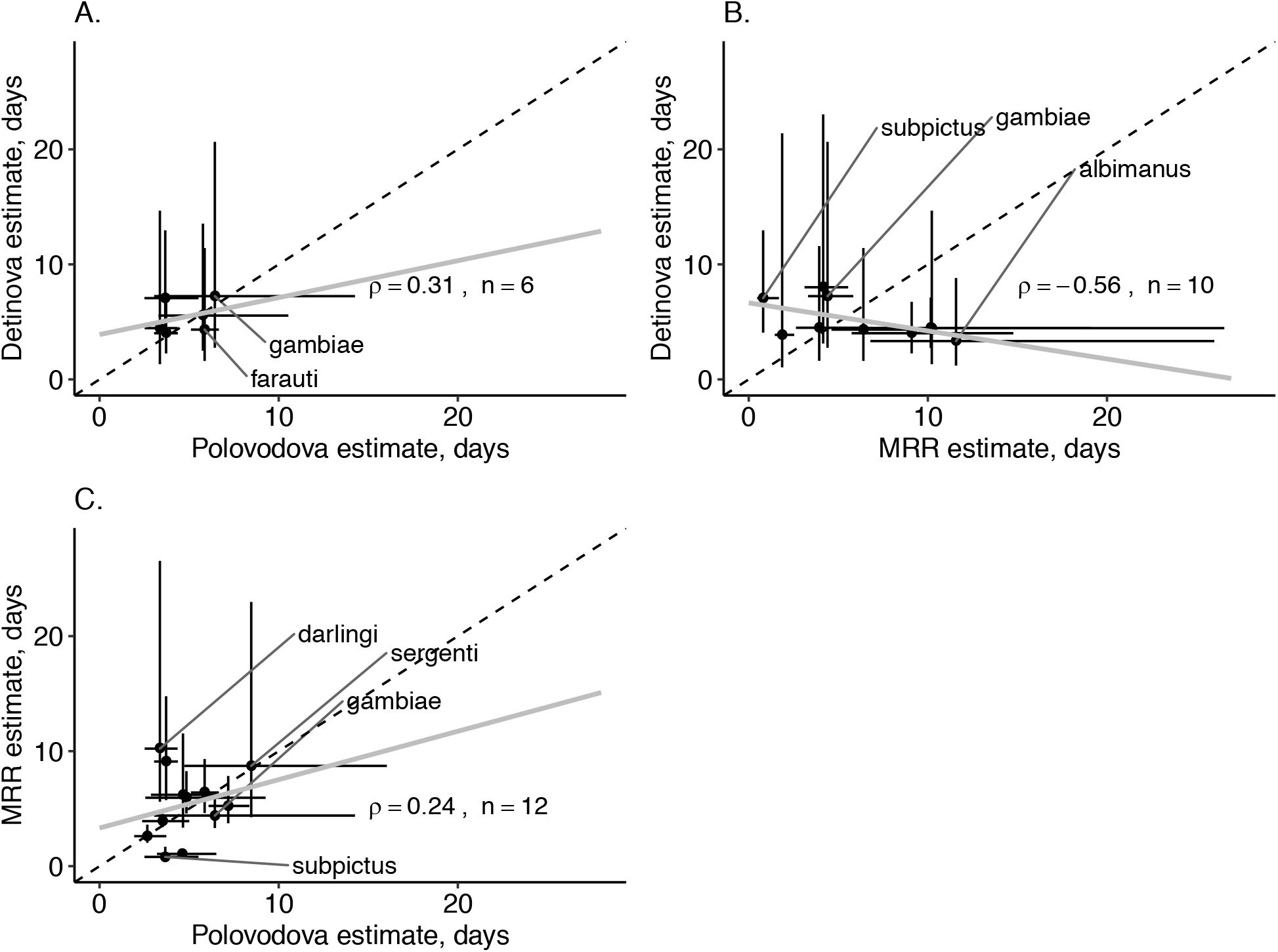
Comparing lifespan estimates. In A, B and C, we show pairwise estimates of lifespan for those species available in both datasets. For the MRR data, we show estimates for females that were neither fed with sugar or blood pre-release. Points indicate the posterior mean estimates and the whiskers show 25% and 75% quantiles. The values of *ρ* indicate Pearson correlations (in no cases were these significant) and *n* indicates sample size.

### The Detinova data does not support an effect of average temperature on the lifespan of African malaria vectors, yet temperature daily range may be important

*An. gambiae s*.*l*. and *An. funestus s*.*l*. are the two most important malaria vector complexes in Africa, and the Massey et al. (2016) dataset has many parity observations for these taxa allowing us to analyse the effects of temperature and temperature range on lifespan (after removing cases where insecticide was used, there were n=257 studies for *An. gambiae s*.*l*. and n=109 for *An. funestus s*.*l*.). We collected environmental data on daily temperature mean and range for the sites and dates where studies were conducted and included them as covariates in our analysis (see SOM). We did not detect a significant association between lifespan and average temperature for either complex. However, there was a domed relationship between daily temperature range and lifespan for *An. gambiae s*.*l*., suggesting these mosquitoes live longest when the range is approximately 1.5<u>º</u>C (Fig. 8, based on a quadratic regression model). Our estimates suggest that at a daily temperature range of 0.5<u>º</u>C, *An. gambiae s*.*l*. lifespan is approximately 9.1 days, which increases by nearly 50% if the daily temperature range increases by 1<u>º</u>C (change of 4.3 days: 95% CI: 2.2-6.4 days). We did not detect a relationship between temperature range and lifespan for *An. funestus s*.*l*..

**Figure 8:**
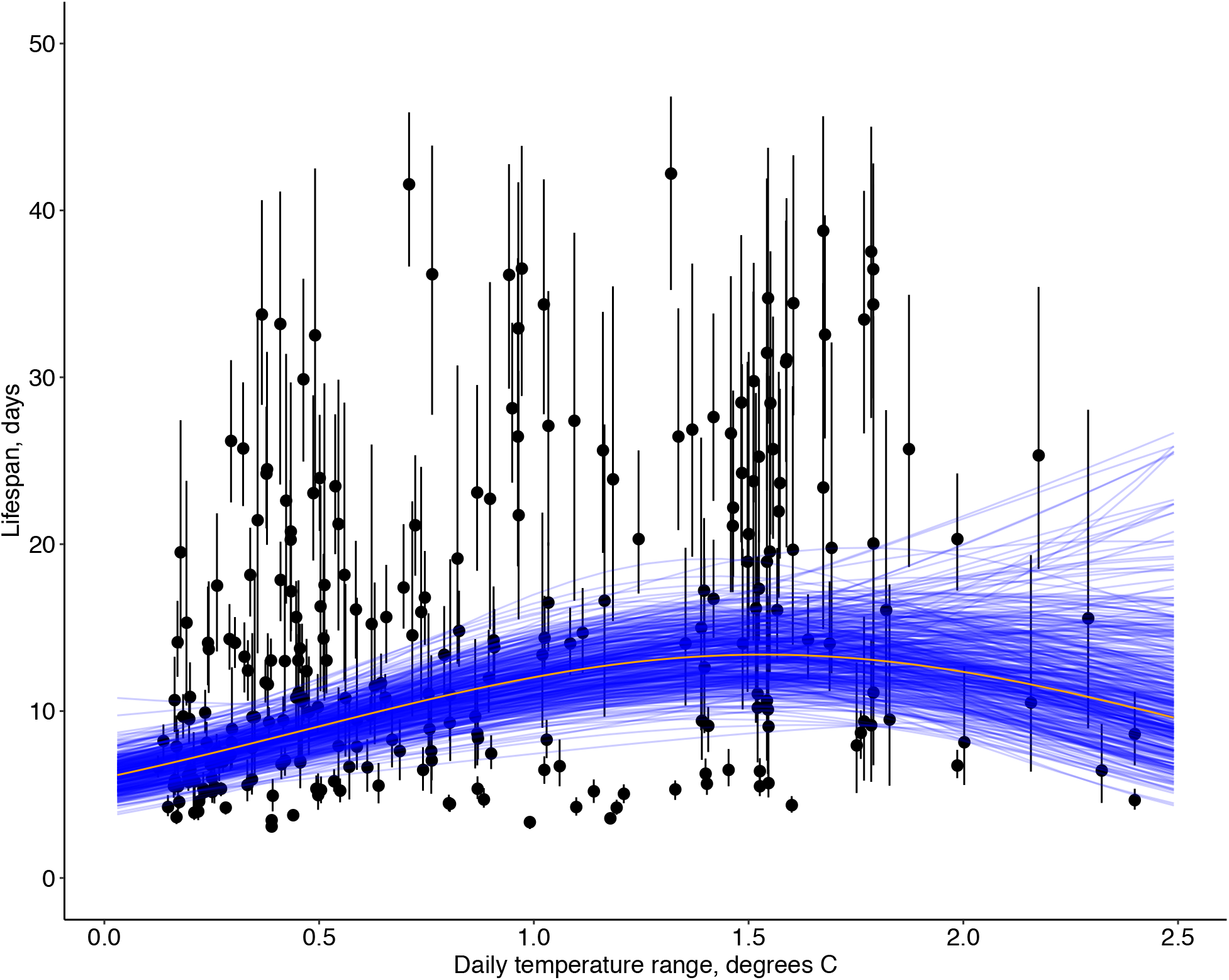
Detinova dissection: day-night temperature range versus lifespan estimates for *Anopheles gambiae s*.*l*.. Points indicate the posterior mean estimates and the whiskers show 25% and 75% quantiles. Each blue line indicates a posterior estimate of the relationship between lifespan and temperature range; the orange line indicates the posterior median.

### Most mosquitoes do not live long enough to transmit disease

The extrinsic incubation period of a vector-borne disease (EIP) is the time required for a pathogen ingested in one blood meal to become transmissible during a future feeding event (at the minimum it must be greater than one gonotrophic cycle). In order to transmit a disease, a mosquito must live for at least as long as the EIP, with additional time required for finding a host and possibly a mate. We next use our lifespan results, from all three methodologies, to estimate how many mosquitoes are likely to survive beyond this lower bound for different vector-disease combinations (Fig. 9).

**Figure 9:**
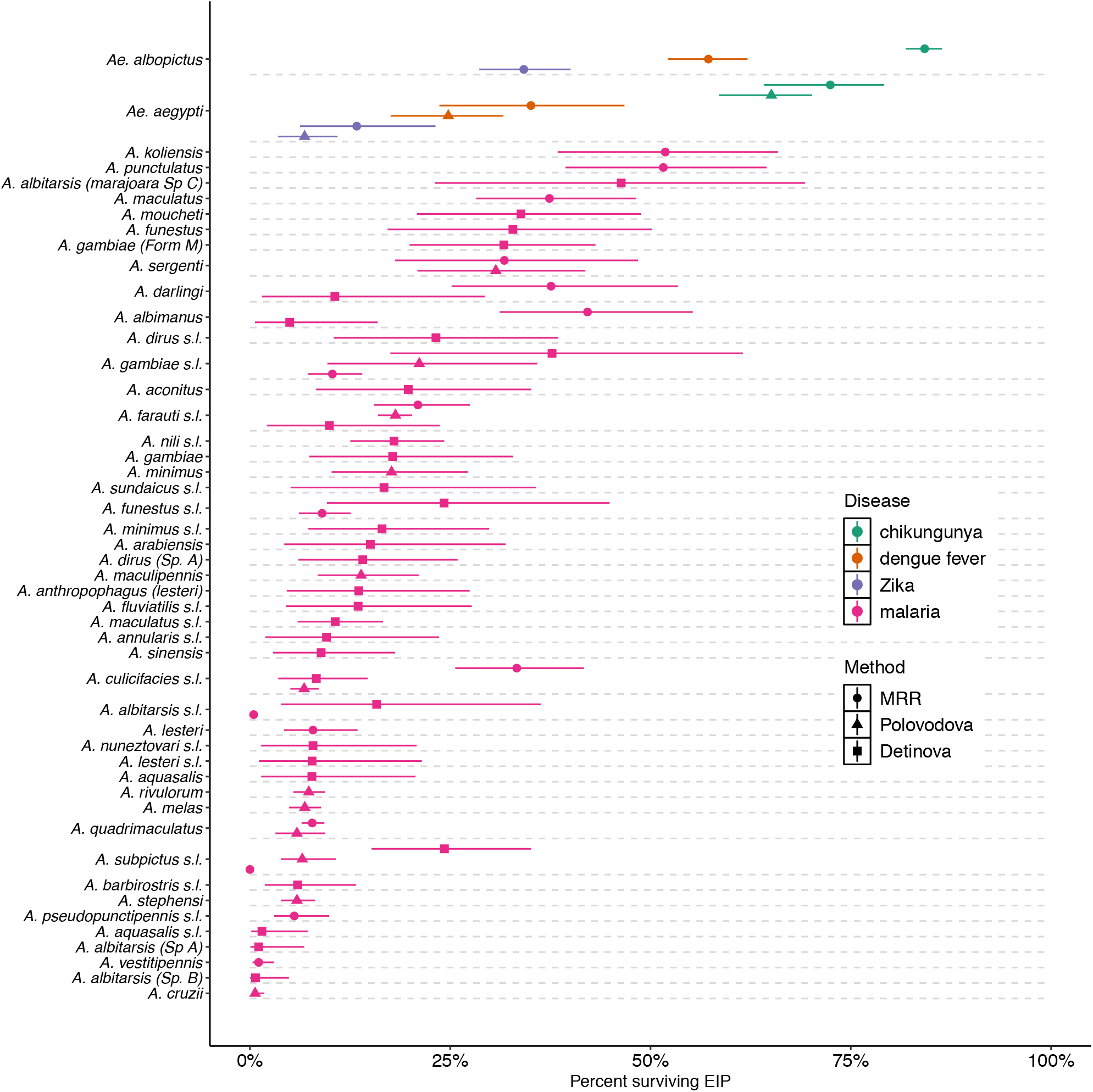
Proportions of mosquitoes surviving EIPs of malaria, dengue, chikungunya and Zika. Colours indicate the measures for the different diseases; shapes indicate the dataset used to produce the estimate. We assumed that the EIPs were: malaria-10 days, chikungunya-2 days, dengue fever-6.5 days and Zika-12.5 days. The left and right whiskers indicate the 25% and 75% posterior quantiles, and the points indicate the mean. All estimates were obtained using the exponential survival model.

Our results suggest that for most anopheline species only a small proportion of individuals live long enough to transmit the malaria pathogen. For the major malaria vector *An. gambiae s*.*l*., lifespan estimates from the Detinova, Polovodova and MRR methods suggested that 38% (25%-75% CI: 17%-62%), 21% (CI: 10%-36%) and 10% (CI: 7%-14%) of individuals survived long enough potentially to transmit malaria. For the longest-lived species, *An. koliensis* and *An. punctulatus*, 52% of individuals were predicted to live longer than the EIP, both according to MRR studies.

*Ae. aegypti* and *Ae. albopictus* are the main vectors of dengue, chikungunya and Zika viruses. These viruses have short EIPs in comparison with malaria, and hence we estimate that a greater fraction of mosquitoes will live long enough to transmit the diseases. The highest estimate (obtained from the MRR analysis) was 86% for *Ae. albopictus* transmitting chikungunya.

### The MRR and Polovodova data suggests that mosquito senescence may occur in a minority of wild mosquito populations

So far, we have assumed that mosquitoes die at a constant rate throughout their adult lifetimes. If mosquitoes tend to senesce, however, we should expect a better fit to the data from using models which allow the mortality rate to increase with age. We tested this using species level lifespan estimates obtained from MRR (Fig. 10), and the Polovodova method (Fig. S5). We included five senescence models in addition to the constant mortality model, allowing a wide range of age-dependent functional forms to be tested simultaneously. To handle ambiguities caused by the multiple comparisons, we created a ranking representing relative evidences for or against senescence at the species level: if all senescence models fitted the data better, we took this to represent evidence in favour of senescence; if the constant mortality model outperformed one or more of the senescence models, we took this to mean both senescence or constant mortality models could explain patterns seen in the data; if none of the senescence models fitted the data better, we took this to be evidence for age-independent mortality.

**Figure 10:**
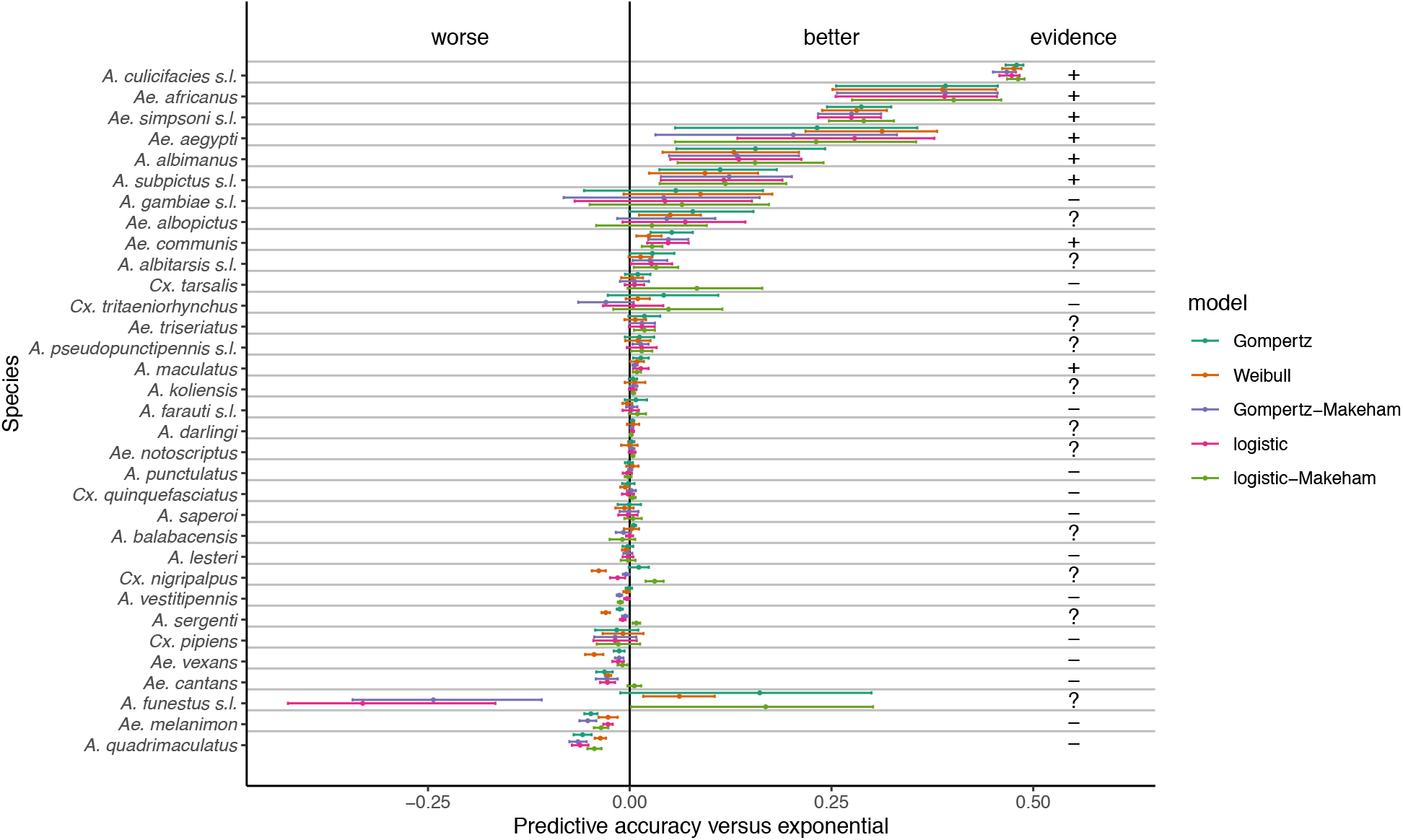
MRR: evidence for senescence. Here, we consider the predictive accuracy of age-dependent models of mortality versus the exponential model by species. The predictive accuracy was determined using K-Fold cross-validation as described in SOM; the measure of accuracy presented here by the central dots is the difference in estimated expected log pointwise predictive density compared to the exponential model. The lower and upper whiskers represent the lower and upper bounds of the 95% confidence interval in predictive accuracy. The species have been ordered so that those with the highest average predictive accuracy across all age-dependent models compared to the exponential model appear at the top. If all age-dependent models outperform the exponential, we deem this as evidence for age-dependent mortality (‘+’); if a subsample of models perform better, we deem the evidence ambiguous (‘?’); and if no age-dependent model exceeds the performance of the exponential, we conclude no evidence in favour of senescence (‘-’).

Using MRR estimates, we detected evidence for and against age-dependent mortality in 8 and 14 species respectively, and mixed evidence in 11 species (Fig. 10). Age-dependent mortality was supported in species including *Ae. aegypti*, the main vector of dengue fever, Zika and chikungunya.

Using Polovodova method estimates, we detected evidence of senescence in only two out of 25 species (*An. gambiae s*.*l*. and *An. minimus*; Fig. S5). There was no overlap between the species exhibiting evidence for senescence from each meta-analysis.

It is possible that some mosquito species do not live long enough in the wild to experience senescence. In support of this hypothesis, we detected a positive correlation between the ranked estimated lifespans of the species and the ranked mean predictive accuracy of age-dependent models for the MRR analysis (Spearman’s rank correlation test:*ρ* = 0.19, p<0.05). However, this was not significant for the Polovodova method analysis (*ρ* = 0.07,p=0.44).

### Discussion

In this study, we analysed a database of mark-release-recapture experiments and two other databases of female mosquito dissection experiments. Our approach enabled us to pool information from disparate experiments, which individually estimate lifespan with considerable uncertainty, to obtain estimates at the species and genus levels. Most of the estimated lifespans were less than 10 days, which is shorter than the EIPs of several important mosquito borne diseases including malaria and Zika. This suggests that relatively few mosquitoes live long enough to transmit these diseases (see, for example, Clements and Paterson, 1981) and highlights the importance of interventions that act to reduce adult lifespan. A further result, from Detinova dissection data, was that applying insecticides in the environment markedly reduces the lifespan of adult Anopheline mosquitoes, which is widely thought to be the main mode of action behind the success of insecticide treated bednets in sub-Saharan Africa (Bhatt, Weiss et al. 2015).

We used Bayesian hierarchical models to partition the variance in the data across different taxonomic ranks. The MRR analysis suggested that most variance in lifespan occurs at the level of the genus rather than the species, yet the reverse was true of the Polovodova analysis. Moreover, these two analyses did not agree on the ordering of genera in terms of lifespan, with MRR data indicating that *Culex* mosquitoes are on average the shortest lived and *Aedes* the longest, while Polovodova finding *Aedes* to be the shortest lived and *Anopheles* the longest. The Detinova data, which was limited to Anopheline species, indicated that there is more variation at the species than species-complex level. These inconsistencies may indicate that the assumptions underpinning the analyses were violated; it could also be that differences in the environment were more important than phylogeny in determining the lifespan of mosquitoes in a given population.

It is also possible that study design hampered our ability to reliably estimate lifespan. To investigate this, we used Monte Carlo simulations to determine how accurately mosquito lifespan could be estimated in an “ideal’’ MRR experiment, where mosquitoes are neither affected by marking nor emigrate out of the study area (see SOM for full details). Our analysis showed that many experiments included in the database had such short study durations or released so few marked mosquitoes that lifespan would be inaccurately estimated due to overwhelming sampling noise (Fig. S6). This may explain why there is so much study level variance in our MRR meta-analysis. Though statistical power can be increased by pooling data across experiments, as we have done here, this is an inefficient approach to studying mosquito longevity. Our Monte Carlo simulations suggest that it is more worthwhile to perform a small number of ambitious MRR experiments, where many mosquitoes are released and recapture efforts last for at least two weeks, than a large number of more modest experiments.

Even in larger scale experiments, MRR may underestimate lifespan for two reasons. First, laboratory experiments have demonstrated that marking can reduce survival (Verhulst, Loonen et al. 2013). It is difficult to quantify the effect of this on our results because the studies included in the meta-analysis used a variety of marking compounds and methods whose impacts may differ. Second, a marked mosquito that dies and another that disperses out of the study area are both not recaptured, meaning that lifespan will be underestimated by analysis of spatially pooled data. In this study, we did not find a significant association between trapping area and lifespan estimate, which would be expected if this was a major source of error.

The two dissection methods assume reproductive age (the number of gonotrophic cycles) can be estimated by examining ovariole structure. Polovodova’s method provides a direct count of the number of cycles and requires highly skilled dissection techniques (Hugo, Quick-Miles et al. 2014); even then, the accuracy of the method has been questioned (Fox and Brust 1994). In particular, it becomes harder for a dissector to distinguish the ovariole remnants of previous gonotrophic cycles as a mosquito ages, which may lead to the underestimation of lifespans. Detinova’s method provides less information, yet it is simpler, and dissections can be carried out reliably and routinely by most field entomologists, which probably explains its more widespread use.

Lifespan estimates from both dissection methods are sensitive to the assumption that population size is stable; lifespan will tend to be underestimated if the mosquitoes are collected from a growing population and vice versa. We will have reduced, but not eliminated, this bias by pooling data that were collected at different times of year for each study site where multiple collections were made. It is also possible that lifespan estimates will be affected by the trapping method, which may over-represent some life-stages at the expense of others. Typically, mosquitoes are caught when they attempt to blood-feed, and there are differing opinions about whether nulliparous females are more (Clements and Paterson 1981) or less (Gillies and Wilkes 1965) likely to be sampled. A further challenge to estimating chronological lifespan from dissection data is uncertainty in the length of the gonotrophic cycle. This is typically calculated using MRR studies or by observation of wild-caught females, though we found evidence that the method chosen influenced the estimated duration. We found significant variation in gonotrophic cycle length across the three mosquito genera for which we had data.

We used meteorological records to investigate the effect of temperature on lifespan, something that has been shown to be important in laboratory studies (Yang, Macoris et al. 2009, Murdock, Paaijmans et al. 2012, Beck-Johnson, Nelson et al. 2013, Brady, Johansson et al. 2013). We found no relationship which may be because our meteorological variables were too imprecise, ignoring weather variation around average temperatures. Alternatively, the effects of temperature may be smaller in the field than the laboratory if mosquitoes adjust their behaviour to buffer themselves against temperature extremes, for example by seeking out cool, shady micro-habitats during periods of high temperature. We did find that longevity was significantly correlated with the daily temperature range in *An. gambiae s*.*l*. (there was a similar non-significant trend in *An. funestus s*.*l*.) but are uncertain how to interpret this finding.

There is evidence that mosquito mortality rates rise with age (senescence) from laboratory studies (Styer, Carey et al. 2007, Dawes, Churcher et al. 2009) and a single field experiment (Harrington, Vermeylen et al. 2014). By fitting a range of survival models to the data in two meta-analyses, we tested for age-dependent mortality. We found limited evidence for senescence. To help interpret these results, we conducted a simulation power analysis of a typical MRR study to understand the factors affecting the likelihood of detection of senescence (Section S3). This revealed that study duration is very important, but the initial release size much less so. For the ‘moderate’ senescence that we modelled, the recapture efforts must continue for more than 18 days to detect senescence 80% of the time (Fig. S7), yet the median duration of experiments in the MRR dataset was only 10 days. It may be that mosquitoes in the field seldom live long enough to experience the senescence observed under laboratory conditions, but if they do then carefully designed and extended MRR experiments will be required to detect it. (Clements and Paterson 1981) conducted a meta-analysis of MRR and dissection field experiments and determined that mortality increased with age at a rate comparable to the moderate senescence we consider in the power analysis. To our knowledge, the MRR study of (Harrington, Vermeylen et al. 2014) on *Ae. aegypti* in Thailand has been the sole field experiment that has detected senescence: here, laboratory-reared mosquitoes of different ages were marked and released. It is possible that this study was able to detect senescence because mosquitoes of ages up to 20 days old were released, which considerably exceeds our estimates of wild lifespan.

As has been found in laboratory studies (Styer, Carey et al. 2007, Dawes, Churcher et al. 2009), our analysis of the MRR data indicates that female mosquitoes outlive males in the field, although the magnitude of this difference is not as great. Because of ethical concerns about releasing female vectors, many recent MRR experiments use only males (Fig. SM1). Our results suggest caution in extrapolating results from one sex to the other.

## Conclusion

We applied modern statistical methods to analyse field data collected over 99 years to produce a set of lower bound estimates of mosquito lifespan. Though the application of tailored statistical models to particular data sets may maximise the information obtained from each, our approach of using the same methods across all data sets allows different studies to be pooled and broader patterns to be examined. We found considerable variation among populations, and relatively few, and then generally weak, emergent patterns. There was weak or no concordance between lifespan estimates obtained by MRR and dissection methods. These results combined with our power analyses suggest that the majority of field studies conducted to estimate lifespan are of sufficient scale to obtain robust results, and we suspect that temporal and spatial variation in lifespan, something that is rarely assessed, may also frustrate simple conclusions about lifespan patterns. For MRR in particular, allocating resources to large-scale experiments rather than a large number of small-scale experiments is likely to be more fruitful. The challenges of statistically estimating lifespan from population data are thus very great and strongly justify continuing research into biochemical and genetic markers of individual age which, if successful, would revolutionise our ability to estimate this most important parameter.

## Materials and Methods

In recent years, many important vectors of disease have been shown to be complexes of closely related species, biotypes or forms that cannot be distinguished morphologically (for example, the morphospecies *Anopheles gambiae sensu lato* is now separated into the widespread *gambiae, coluzzii, arabiensis* and a number of more local species). In the MRR and Polovodova-dissection analyses, most data were collected before molecular techniques allowed these taxa to be separated, and, for these, we work chiefly with morphospecies. In the Detinova-dissection analyses, more detailed species-level information was often available, and we estimate lifespans for both species and morphospecies. A detailed description of methods is provided in the SOM file.

### Mark-release-recapture (MRR)

Data from MRR experiments in the database of (Guerra, Reiner et al. 2014) were examined. Of the 232 data sets, 179 involved only females, 35 males, and 18 both sex releases. For 102 data sets, the age of the released mosquitoes was known (the average age of released mosquitoes was 4.0 days) while in the other cases it was unknown or unrecorded; in these cases, we assumed the mosquitoes were newly emerged at the time of release. See Tables SM1 & SM2 for a summary of data characteristics.

We analysed all MRR experiments within the same statistical framework. In the simplest case, mosquitoes are released on day zero, and the probability that they remain in the recapture area until day *t* is *S*(*t*) when they are recaptured with probability *ψ*. We model the number of mosquitoes recaptured on day *t* using a negative binomial sampling model with mean (*N*_*R*_ − *Y* (*t* − 1)) *S* (*t*) *ψ*, where *Y* (*t* − 1) is cumulative captures before day *t*, and inverse-overdispersion parameter *κ*. The negative binomial has been used previously in analyses of mosquito count data (Service, 1971; Nedelman, 1983) because of its ability to represent temporal over-dispersion in recaptures most likely caused by variable weather. A slight modification was required for studies with multiple releases (see SOM).

The simplest model for *S*(*t*) assumes there is a constant probability (*λ*) that a mosquito dies or leaves the recapture area so that the numbers remaining after time *t* are given by the exponential distribution, exp(−*λt*). We utilised this form extensively but in testing for senescence used five other models where *λ*(*t*) varies with time so that,

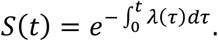

Details of the five models (Gompertz, Weibull, Gompertz-Makeham, Logistic and Logistic-Makeham), which vary in their ability to detect different forms of age-dependent mortality, are given in the SOM. Using multiple different types of models increased our chances of detecting senescence though also increased the likelihood of false positives.

We used two analyses to estimate lifespan. One considered each study separately; the other involved a hierarchical approach which grouped studies according to taxonomy. Parameters were estimated using Bayesian techniques with relatively uninformative priors for *κ* and the parameters of *λ*(*t*), but assuming a prior for *ψ* indicating a low recapture probability (bounded in part by knowledge of the maximum daily recapture rates; see SOM).

Using the hierarchical model, we estimate distributions of lifespan at the species and the genus levels, and across a dataset where experiments with fewer than six recaptures and those species within only a single study were removed. This procedure assumes that there is a distribution of lifespan parameters for each species from which those governing individual MRR time series are sampled, and similarly a distribution at the genus level from which those for individual species are derived (rather akin to random effects in classical statistics). Within this framework, we can also allow the parameters for individual time series to be influenced by covariates such as differences in experimental methodology. Posterior distributions were derived using Markov Chain Monte Carlo (MCMC) methods with convergence assessed using the 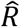 statistic (Gelman and Rubin 1992). The predictive power of the model was assessed using *K* -fold cross validation which tests the ability of the model fitted to part of the data to predict the rest using multiple different partitions. Further details of the prior specification, fitting and validation through posterior predictive checks (Lambert 2018) are given in the SOM.

Two studies of *Anopheles balabacensis* reported capture rates increasing with time, presumably reflecting a violation of our assumption of constant recapture probabilities. We omitted this species from the analysis.

## Dissection

### Polovodova’s method

Studies using Polovodova’s dissection method to determine reproductive age were located in literature databases using relevant keyword, citation and author searches, and by checking previous studies cited by the papers located (see SOM). The list of studies included with associated metadata is available as a Supplementary data file.

Most dissection studies recorded the distribution of reproductive age (nulliparous, uniparous, biparous and so on) in wild-caught mosquito samples collected over a specific period of time. Overall, we found 568 physiological age cross-sections recorded at distinct times in 72 published articles. Our statistical approach assumes stable population sizes. To guard against the effect of fluctuating population sizes on our analysis, we aggregated the data at a given location across cross-sections taken at different times. We further omitted time series with fewer than 100 mosquitoes and for species with only one data set, leaving 131 studies of mosquitoes in the genera *Anopheles, Aedes, Culex* and *Mansonia*.

The dissection data which we use provides information on the distribution of ages within each population. By assuming that population sizes were fixed throughout the period of investigation, this allows us to estimate mean lifespan using a statistical model of mortality incorporating the probability of mosquito capture. We modelled the number of mosquitoes found by dissection to be of age *a* using the negative binomial distribution with mean Ψ*S*(*a*) and shape parameter *κ*, where Ψ is the product of the recruitment rate of adult mosquitoes and the probability of being captured for dissection, and *S* (*a*) is the probability of surviving until age *a*. We used the number of females that have yet to lay eggs (nulliparous) to estimate the recruitment rate as described further in the SOM. Initial examination revealed that in some data sets the number of nulliparous females was anomalously low, something that has been noticed before (Gillies and Wilkes 1965). In data sets where the fraction of nulliparous females was less than 90% the uniparous females (completed one gonotrophic cycle), we excluded the nulliparous observation. Data was analysed using a Bayesian framework similar to that used to analyse the MRR data, with minor differences in the specification of the priors (see SOM).

### Detinova’s method

Detinova and Bertram (1962) provide an alternative dissection method to estimate the age of a given female mosquito, which results in a binary observation for each specimen: nulliparous or parous. As for the analysis of dissection data from Polovodova’s method, we assume that constant population sizes are constant and with knowledge of a given gonotrophic cycle duration (see below), we can estimate mean population lifespan (see SOM).

Massey, Garrod et al. (2016) provides a database estimates of different anopheline bionomic variables, which includes 1490 observations of parity using Detinova’s method. As for the other two analyses, we use a Bayesian framework. The likelihood assumed is a binomial distribution with sample size given by the number of specimens dissected and probability parameter representing the proportion parous in the wild population. The probability parameter is allowed to vary according to experiment but are assigned hierarchical beta priors (see SOM) that allow partial pooling of observations according to a grouping (species, morphospecies, genus and so on).

### Gonotrophic cycle duration estimates

To compare lifespan estimates from the MRR and dissection analyses, we need to convert physiological age into chronological age. To do this, we conducted a meta-analysis of previously published studies that estimate the duration of the gonotrophic cycle (see SOM). This was supplemented by references in the review by (Silver 2007). Whilst compiling our dataset on gonotrophic cycles, Massey et al. (2016) published a database of bionomic quantities for malaria vectors (that is, including only anopheline species). Included in this dataset were estimates of gonotrophic cycle duration. After removing duplicates with our dataset, we were left with 120 estimates of gonotrophic cycle duration.

Most published estimates of gonotrophic cycle duration were obtained by observing wild-caught specimens or their progeny in the laboratory or by dissecting females recaptured in MRR studies. Studies differed greatly in how (if at all) they represented uncertainty in their estimates. Where confidence limits were given, we treated these as the relevant quantiles of a normal distribution; where a range was stated (e.g. “4-6 days”), we interpreted the bounds as the 2.5% and 97.5% quantiles of a normal distribution; and where a single figure was quoted, we assumed this was the mean of this distribution. Using the quantiles of the normal distribution, we estimated its mean and standard deviation separately for the genera *Anopheles, Aedes* and *Culex* by regression (see SOM).

We converted physiological age to chronological age by sampling from this distribution to obtain a particular gonotrophic cycle length for each mosquito (see SOM).

## Supporting information

Polovodova dissection papers summary

Polovodova dissection series

Gonotrophic cycle data

Posterior predictive checks for MRR series

Aggregated Polovodova dissection series

## Acknowledgements

The authors would like to thank Mike Bonsall, Austin Burt, Thomas Churcher and Steve Lindsay for useful conversations throughout the course of this work.

## References

Beck-Johnson, L. M., W. A. Nelson, K. P. Paaijmans, A. F. Read, M. B. Thomas and O.N. Bjørnstad (2013). “The effect of temperature on Anopheles mosquito population dynamics and the potential for malaria transmission.” PLOS one 8(11): e79276.

Bhatt, S., D. Weiss, E. Cameron, D. Bisanzio, B. Mappin, U. Dalrymple, K. Battle, C. Moyes, A. Henry and P. Eckhoff (2015). “The effect of malaria control on Plasmodium falciparum in Africa between 2000 and 2015.” Nature 526(7572): 207–211.

Brady, O. J., M. A. Johansson, C. A. Guerra, S. Bhatt, N. Golding, D. M. Pigott, H. Delatte, M. G. Grech, P. T. Leisnham, R. Maciel-de-Freitas and vectors (2013). “Modelling adult Aedes aegypti and Aedes albopictus survival at different temperatures in laboratory and field settings.” Parasites & vectors 6(1): 1–12.

Carter, R. and K. N. Mendis (2002). “Evolutionary and historical aspects of the burden of malaria.” Clinical microbiology reviews 15(4): 564–594.

Clements, A. and G. Paterson (1981). “The analysis of mortality and survival rates in wild populations of mosquitoes.” Journal of applied ecology: 373–399.

Dawes, E. J., T. S. Churcher, S. Zhuang, R. E. Sinden and M.-G. Basáñez (2009). “Anopheles mortality is both age-and Plasmodium-density dependent: implications for malaria transmission.” Malaria journal 8(1): 1–16.

Detinova, T. S. and D. S. Bertram (1962). Age-grouping methods in Diptera of medical importance, with special reference to some vectors of malaria, World Health Organization.

Fox, A. and R. Brust (1994). “How do dilatations form in mosquito ovarioles?” Parasitology today 10(1): 19–23.

Gelman, A. and D. B. Rubin (1992). “Inference from iterative simulation using multiple sequences.” Statistical science 7(4): 457–472.

Gillies, M. T. and T. Wilkes (1965). “A study of the age-composition of populations of Anopheles gambiae Giles and A. funestus Giles in North-Eastern Tanzania.” Bulletin of entomological research 56(2): 237–262.

Guerra, C. A., R. C. Reiner, T. A. Perkins, S. W. Lindsay, J. T. Midega, O. J. Brady, C. M. Barker, W.K. Reisen, L. C. Harrington, W. Takken and vectors (2014). “A global assembly of adult female mosquito mark-release-recapture data to inform the control of mosquito-borne pathogens.” Parasites & vectors 7(1): 1–15.

Harrington, L. C., F. Vermeylen, J. J. Jones, S. Kitthawee, R. Sithiprasasna, J. D. Edman and T. W. Scott (2014). “Age-dependent survival of the dengue vector Aedes aegypti (Diptera: Culicidae) demonstrated by simultaneous release–recapture of different age cohorts.” Journal of medical entomology 45(2): 307–313.

Hugo, L. E., J. A. Jeffery, B. J. Trewin, L. F. Wockner, N. Thi Yen, N. H. Le, L. T. Nghia, E. Hine, P. A. Ryan and B. H. Kay (2014). “Adult survivorship of the dengue mosquito Aedes aegypti varies seasonally in central Vietnam.” PLOS neglected tropical diseases 8(2): e2669.

Hugo, L. E., S. Quick-Miles, B. Kay and P. Ryan (2014). “Evaluations of mosquito age grading techniques based on morphological changes.” Journal of medical entomology 45(3): 353–369.

Lambert, B. (2018). A student’s guide to Bayesian statistics, Sage.

Macdonald, G. (1957). The epidemiology and control of malaria, Oxford University Press.

Massey, N. C., G. Garrod, A. Wiebe, A. J. Henry, Z. Huang, C. L. Moyes and M. E. Sinka (2016). “A global bionomic database for the dominant vectors of human malaria.” Sci data 3: 160014.

Murdock, C., K. P. Paaijmans, A. S. Bell, J. G. King, J. F. Hillyer, A. F. Read and M. B. Thomas (2012). “Complex effects of temperature on mosquito immune function.” Proceedings of the Royal Society B: Biological Sciences 279(1741): 3357–3366.

Polovodova, V. (1949). “The determination of the physiological age of female Anopheles by the number of gonotrophic cycles completed.” Meditsin-skaia Parazitologiia Parazitar Bolezni 18: 352–355.

Silver, J. B. (2007). Mosquito ecology: field sampling methods, springer science & business media.

Sinka, M. E., M. J. Bangs, S. Manguin, M. Coetzee, C. M. Mbogo, J. Hemingway, A. P. Patil, W. H. Temperley, P. W. Gething and C. W. Kabaria (2010). “The dominant Anopheles vectors of human malaria in Africa, Europe and the Middle East: occurrence data, distribution maps and bionomic précis.” Parasites & vectors 3(1): 1-34 %@ 1756-3305.

Sinka, M. E., M. J. Bangs, S. Manguin, M. Coetzee, C. M. Mbogo, J. Hemingway, A. P. Patil, W. H. Temperley, P. W. Gething, C. W. Kabaria and vectors (2010). “The dominant Anopheles vectors of human malaria in Africa, Europe and the Middle East: occurrence data, distribution maps and bionomic précis.” Parasites & vectors 3(1): 1–34.

Styer, L. M., J. R. Carey, J.-L. Wang and T. W. Scott (2007). “Mosquitoes do senesce: departure from the paradigm of constant mortality.” The American journal of tropical medicine and hygiene 76(1): 111.

Verhulst, N. O., J. A. Loonen, W. Takken and vectors (2013). “Advances in methods for colour marking of mosquitoes.” Parasites & vectors 6(1): 1–7.

Yang, H., M. Macoris, K. Galvani, M. Andrighetti and D. Wanderley (2009). “Assessing the effects of temperature on the population of Aedes aegypti, the vector of dengue.” Epidemiology & infection 137(8): 1188–1202.

