## Supplementary methods for "A Meta-analysis of Longevity Estimates of Mosquito Vectors of Disease"

### A Meta-analysis of Longevity Estimates of Mosquito Vectors of Disease: Additional Methods

May 30, 2022

#### SM1 MRR experiments

##### SM1.1 Data

The MRR database in Guerra *et al.* (2014) contains 393 individual time-series, along with meta-data for a range of factors for each experiment (for example, species, geography, and date of study). We cleaned the data, making a number of amendments to it (see Section SM1.2) and transformed the data into a form amenable for our statistical model. We removed time-series with fewer than six separate recapture observations, resulting in 238 time series. For the species-level analyses, we required at least two time series per species, resulting in loss of six series, meaning that the data used in the Bayesian hierarchical analysis comprised 232 individual time series (see Section SM5 for a list of the studies included). Tables SM1 and SM2 summarise the data used in the Bayesian hierarchical analysis. This data encompassed time-series from MRR experiments across 33 different species and three genera: *Aedes* (91 separate time-series), *Anopheles* (94), and *Culex* (47), and spanning a wide geographical range (Fig. SM2).

##### SM1.2 Amendments by MRR ID

This section describes the amendments made by MRR ID (the unique identifier that Guerra *et al.* (2014) use for each MRR record). Where there was ambiguity in the data, we attempted to consult the original publication. The amendments to the dataset are described below:

- 69 and 70: Since the release was not disaggregated by sex, the recapture data were aggregated across both males and females, resulting in a single time series in each case.

- 482: There are actually two series (one with captured mosquitoes, the other with young reared) in the paper (Rawlings *et al.* , 1981), whereas the database contains only one recapture series. In this analysis, both time series are included.

- 483: There are three studies in the paper (Rawlings *et al.* , 1981), (by dye colour: there is one for 'M&Y', another for white, and a third for green), whereas the database only contains one release series. In this analysis, we analyse all three time series separately.

- 578: A null observation was removed on day 12, since it isn't present in the original paper (Eyles *et al.* , 1943a).

- 585: It is not clear from the original paper (Smith *et al.* , 1941) whether missing recapture observations on certain days are because no mosquitoes were found on those days, or because no efforts were made to recapture them on those days. We have assumed that the number recaptured were as stated in the database on those days (missing, because no recapture efforts were made on those days).

- 586: It was unclear from the database whether the missing observations were due to lack of recapture, or no recapture effort being made on those days. Have assumed that zero mosquitoes were captured on these days (the paper could not be accessed).

- 342: Changed number of released from 6000 to 12,000 following the original publication (Bryan *et al.* , 1991).

- 246: Moved the observations from captures to recaptures following the original publication (Arredondo-Jiménez *et al.* , 1998).

- 55: Corrected a mistake in release data; the actual number released is 739, not 749 (Midega *et al.* , 2007).

- 478-481: All four of these series give releases not disaggregated by gender, whereas the recaptures are split this way. Have combined the recapture series to yield a single release-recapture series in each case.

- 154: Moved captured male series to recaptured following original paper (Tsuda *et al.* , 2001).

- 334: Data starts on day 3, thus have added this number of days to our time variable in this case.

| **Data characteristic** | **Count** |
|---|---|
| Species | 33 |
| Genera | 3 |
| *Anopheles* | 94 |
| *Aedes* | 91 |
| *Culex* | 47 |
| Female-only MRR | 179 |
| Male-only MRR | 35 |
| Mixed sex MRR | 18 |
| Pre-release blood-feeding only MRR | 71 |
| Pre-release sugar-feeding only MRR | 41 |
| Pre-release both blood- and sugar-feeding MRR | 4 |
| Pre-release neither blood- and sugar-feeding MRR | 116 |
| MRR time-series | 232 |

Table SM1: **MRR: summary of time-series used in hierarchical analyses.** For the ‘Species’ and ‘Genera’ rows, the counts represent the number of distinct entries in the database; otherwise the counts represent the number of time series.

| **Data characteristic** | **Min** | **Mean** | **Median** | **Max** | **Standard deviation** |
|---|---|---|---|---|---|
| Study duration, days | 6.0 | 11.8 | 10.0 | 71.0 | 4.7 |
| Number of days on which collections took place | 6.0 | 10.1 | 9.0 | 47 | 9.1 |
| Number of separate release days | 1.0 | 1.9 | 1.0 | 23 | 3.0 |
| Number released | 66 | 4,929 | 1,297 | 86,200 | 12,043 |
| Number recaptured | 2 | 163 | 63 | 4,090 | 399 |
| Recapture percentage | 0% | 8.6% | 5.2% | 57.1% | 10.04% |
| Age at release, days | 0 | 1.8 | 0 | 13 | 2.8 |

Table SM2: **MRR: summary of data from individual time-series used in hierarchical analyses.**

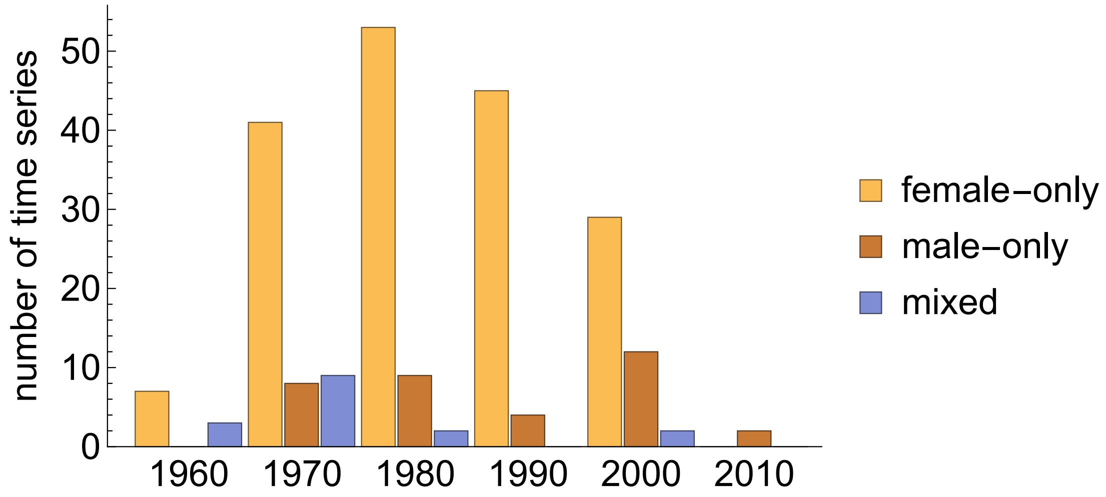


Figure SM1: **MRR: time-series by date and sex of released mosquitoes.** The numbers represent by-decade totals of the 232 time-series we used in the main analysis.

#### SM1.3 Statistical model

Data for a typical MRR experiment consists of a single release of $N_R$ marked mosquitoes followed by a series of marked mosquito recaptures on subsequent days (Fig. SM3 shows three such example series). We model the number $y(t)$ of marked mosquitoes recaptured on day $t$ using a negative binomial sampling model,

$$y(t) \sim \text{NB}\left((N_R - Y(t-1))S(t)\psi, \kappa\right), \tag{1}$$

where $Y(t-1)$ is the cumulative number of mosquitoes caught on all days before $t$, $S(t)$ is the probability that an individual mosquito survives and remains in the study area until time $t$, $\psi$ is the daily recapture probability for an individual mosquito which is assumed constant through time, and $\kappa$ is the time-independent shape parameter of the negative binomial distribution that controls the extent of variance in recapture rate likely due mostly to environmental heterogeneity. We chose a parameterisation of the negative binomial such that its mean is given by $\mu(t) = (N_R - Y(t-1))S(t)\psi$ and its variance by $\sigma(t)^2 = \mu(t) + \frac{\mu(t)^2}{\kappa}$.

For some MRR experiments, there were a number of releases ($q \geq 2$) of marked mosquitoes throughout the duration of the study. In contrast to single releases, with multiple releases over time, it is not in general possible to determine the particular release to which a recaptured mosquito belongs. To avoid the complication of directly inferring this quantity, we choose to represent previous recaptures probabilistically. This results in a slightly different mean to that of the single release

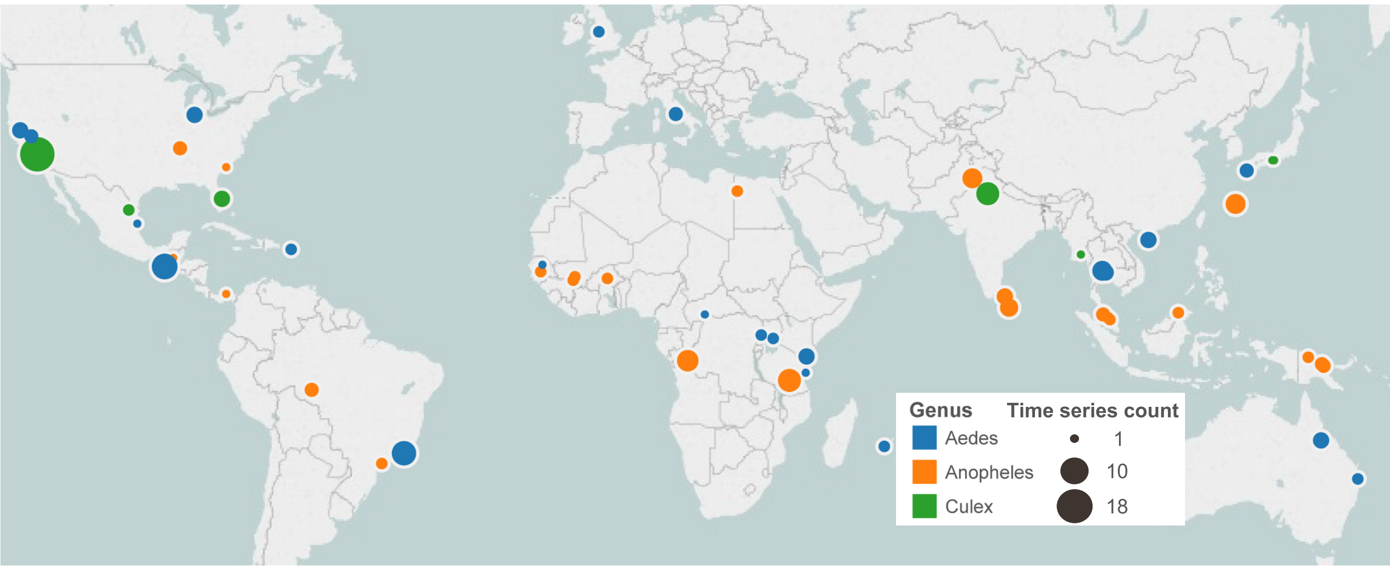


Figure SM2: **MRR: location of studies included in the analysis.** The area of each circle indicates the number of time-series at each study site. The colour shows the genus of the mosquitoes in each study.

model,

$$\mu(t) = N_{Released}(1-\psi)^{t-1}S(t)\psi, \tag{2}$$

where the factor of $(1-\psi)^{t-1}$ represents the probability that a mosquito is *not* recaptured on a previous day. Where experiments consisted of two or more releases occurring at distinct points in time, we assumed that recaptures of individual mosquitoes from either batch were independent of one another, although with the same sampling parameters ($\psi$ and $\kappa$). This results in an overall mean composed of the sum of those from all $q$ releases,

$$\mu(t) = \mu_1(t) + ... + \mu_q(t). \tag{3}$$

The simplest assumption is that death and dispersal rates are independent of mosquito age, giving an exponential 'survival' function,

$$S(t) = e^{-\lambda t}, \tag{4}$$

where $\lambda$ is the sum of the rates of death and dispersal from the study area. To estimate the lifespan of mosquitoes, we use this model, partly because of the limited evidence we find for senescence.

We assume that the number of mosquitoes recaptured on one day is independent of the number recaptured on any other day, apart from their joint dependence

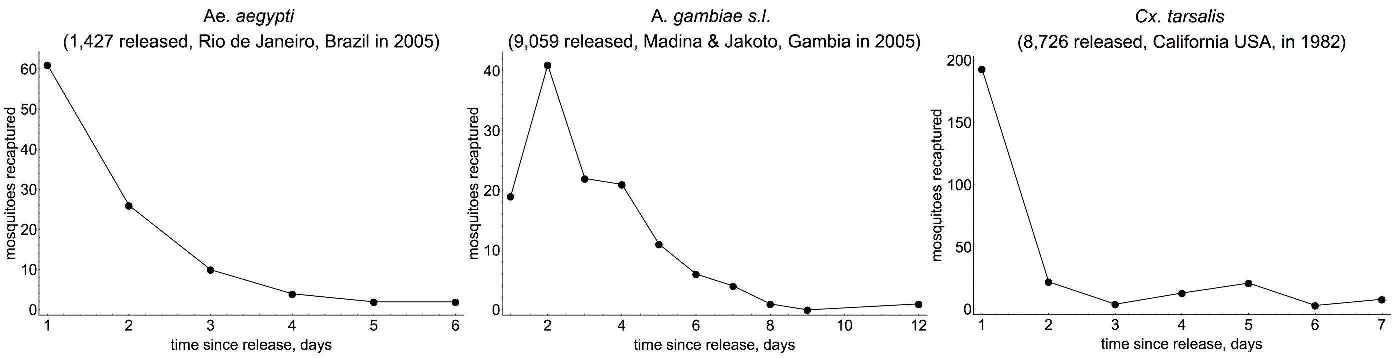


Figure SM3: **MRR: time-series for three different studies from the Guerra *et al.* (2014) database.** In all the above experiments, there was a single release of marked mosquitoes on day zero.

on $\lambda$, $\psi$ and $\kappa$. This results in a likelihood of the data of the form,

$$\mathcal{L}(y(t_1), y(t_2), ..., y(t_R)|\lambda, \psi, \kappa) = \prod_{i=1}^{R} p(y(t_i)|\psi, \lambda, \kappa), \tag{5}$$

where $R$ is the number of individual days where collections took place, and $p(y(t_i)|\psi, \lambda, \kappa)$ is the probability of recapturing $y(t_i)$ mosquitoes on day $t_i$ determined from a negative binomial distribution with parameters $\mu = e^{-\lambda t}\psi$, and $\kappa$.

#### SM1.4 Individual time-series estimates

We first treat each time-series separately, and estimate individual $(\lambda, \psi, \kappa)$ parameters of the statistical model (described in Section SM1.3) for each time-series. To use a Bayesian methodology, we must specify priors on this set of parameters. Here, we choose to specify independent priors on each parameter of the form: $\lambda \sim \mathcal{N}(-2.32, 1)$, $\psi \sim \exp(50)$ (with $\psi$ constrained to be less than 0.1, a threshold below which the majority of daily recapture probabilities in the data are observed) and $\kappa \sim$ log-normal$(2, 1)$. This prior on the rate parameter of the exponential distribution ($\lambda$) corresponds to a wide range of possible lifespans (Fig. SM4A), with a mean of 10 days. It was necessary to use an informative prior on $\psi$ to bound it to sensible values and to prevent unreasonably short estimates of $\lambda$; the prior on $\kappa$ is fairly uninformative and allows a wide range of values (Fig. SM4B,C).

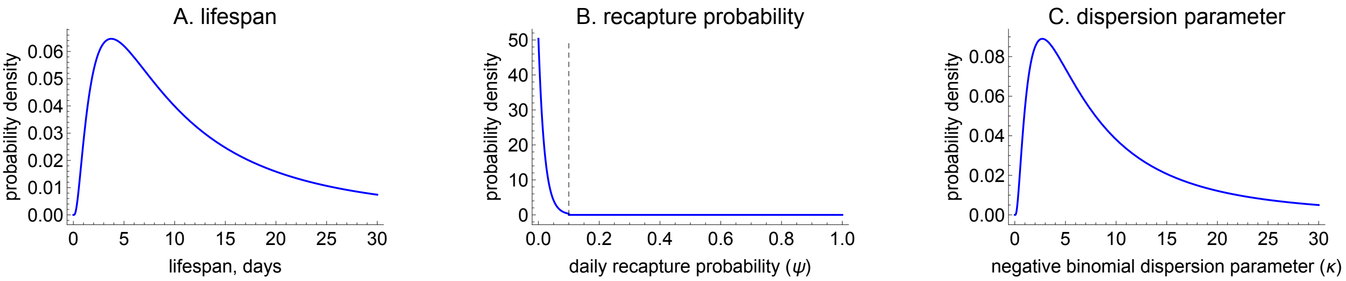


Figure SM4: **MRR: prior probability distributions used for lifespan (A), recapture probability (B) and dispersal parameter (C) in the individual time-series analysis.** Note: the prior for the recapture probability was truncated at $\psi = 0.1$ since most recapture fractions on the first day were significantly below this threshold.

#### SM1.5 Estimating lifespan at the species, genus and overall groupings

We synthesise information from across all MRR experiments to produce more robust estimates of mosquito lifespan than can be obtained from considering the individual time-series separately. However, there exists considerable heterogeneity across the experiments. This heterogeneity has two sources: that arising from variability in experimental methodology; but also that from actual differences in lifespan across the different mosquito cohorts – for example, due to genetic differences between mosquito populations or due to climatic differences. Because of this heterogeneity, we use a Bayesian hierarchical model which is akin to a 'random effects' model in classical statistics. This type of model assumes that there is random variation in parameters at the individual time-series level, although each of the parameters is drawn from a common 'population-level' distribution. In our case, we separately estimate three different model groupings where the population-level distributions correspond to the species, genus and overall (across all studies) levels respectively (Fig. SM5). This allows us to estimate mosquito lifespan for a given species or genus by independent sampling from the posterior predictive distribution from which the individual $\lambda_i$ are drawn.

Whilst we assume that the variation in mosquito mortality across experiments is random in nature, we recognise that there may exist systematic sources of variation that we do not include in our model. To account for this potential source of bias, we examined how the individual estimates of lifespan correlated with experimental covariates (number and density of traps). From this, we determined a slight but insignificant positive correlation between trapping number and density with individual lifespan estimates – suggesting that downward bias due to insufficient

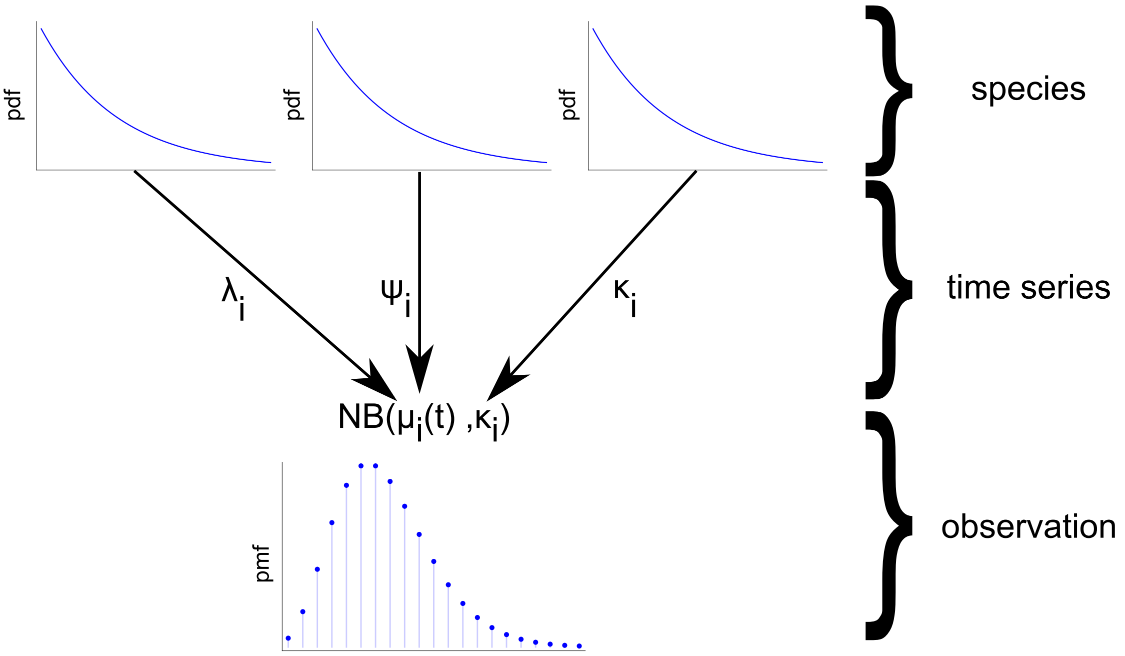


Figure SM5: **MRR: structure of hierarchical Bayesian model.** This diagram represents the structure of the model for the constant mortality model case with the species-level grouping. Here 'pdf' indicates 'probability density function', and $\lambda_i$, $\psi_i$, and $\kappa_i$ represent the force of mortality for the exponential distribution, daily recapture probability and over-dispersion parameter of the negative binomial distribution for time-series $i$ in the dataset; $\mu_i(t) = \psi_i e^{-\lambda_i t}$ is the mean number of mosquitoes recaptured on day $t$.

trapping may be relatively small.

In our hierarchical model, we must specify prior distributions at two levels of the analysis. The first of these links the individual time-series estimates with the overarching group-level distributions. The parameters of these group-level distributions are then set prior distributions (Fig. SM5). As an example, suppose that the rate parameter $\lambda_i$ of our exponential survival model in experiment $i$ is given by:

$$\lambda_i = \exp(c_i + \delta^{m}_{\text{genus}[i]}\text{sex}^{\text{m}}_i + \delta^{\text{mix}}_{\text{genus}[i]}\text{sex}^{\text{mix}}_i + \quad (6)$$

$$\delta^{\text{sugar}}_{\text{genus}[i]}\text{sugar}_i + \delta^{\text{blood}}_{\text{genus}[i]}\text{blood}_i), \quad (7)$$

where $\delta^{m}_{\text{genus}[i]}$ measures a genus-specific effect of male-only releases ($\text{sex}^{\text{m}}_i = 1$) versus female-only ($\text{sex}^{\text{m}}_i = 0$); $\delta^{\text{mix}}_{\text{genus}[i]}$ is the analogous effect for mixed releases versus female-only; $\delta^{\text{sugar}}_{\text{genus}[i]}$ is the analogous effect for mosquitoes that were sugar-fed prior to release versus those that were not fed; and $\delta^{\text{blood}}_{\text{genus}[i]}$ is the analogous effect for mosquitoes that were blood-fed prior to release versus those that were not fed. The parameter $c_i$ is modelled hierarchically, being drawn from a group-level distribution assumed to be normal,

$$c_i \sim \mathcal{N}(\mu_\lambda, \sigma). \tag{8}$$

We then specify priors on these group-level parameters, $\mu_\lambda \sim \mathcal{N}(-2.32, 1)$ and $\sigma \sim \text{log-normal}(-3, 1)$. On all $\delta$ parameters, except $\delta^{\text{mix}}_{\text{genus}[i]}$, we specify standard normal priors. We specify a uniform prior on $\delta^{\text{mix}}_{\text{genus}[i]} \sim U(0, \delta^{m}_{\text{genus}[i]})$, so that the effect of mixed releases lies between the female-only release effect (0) and the male-only release effect ($\delta^{m}_{\text{genus}[i]}$).

The relatively complex nature of priors in hierarchical models make it important to determine their influence on inferences. We chose the above – somewhat uninformative – priors, to allow a range of mosquito lifespans in order to minimise their effect on the estimates we report (Fig. SM6). We also chose to set hierarchical priors on the remaining parameters in our models – $\psi$ the probability of daily recapture, and $\kappa$ the 'over-dispersion' parameter of a negative binomial distribution. For $\psi$, we chose an informative prior that placed all probability mass $\psi \leq 10\%$, since in most experiments the fraction of mosquitoes captured was significantly below this value. For $\kappa$, we chose a fairly wide prior that had most of its probability mass below $\kappa = 20$ (Fig. SM7). In all cases, the hierarchical priors were chosen to have similar implications on lifespan, recapture probability and over-dispersion at the individual time-series level as for the non-hierarchical analysis described previously.

#### SM1.6 Testing for age-dependent mortality in wild mosquitoes

Previous work has found evidence of age-dependent mortality in lab populations (Styer *et al.* , 2007; Dawes *et al.* , 2009) and less commonly in wild mosquitoes (Clements & Paterson, 1981; Harrington *et al.* , 2008). Furthermore, modelling work has examined the implications of departures from a constant risk of mortality (Styer *et al.* , 2007; Hancock *et al.* , 2009; Novoseltsev *et al.* , 2012). To determine whether senescence occurs in wild mosquitoes, we re-estimate the model using a range of different survival functions, $S(t)$, that permit mortality to increases with age. Specifically, we re-estimate our model using five other models that were previously used in the literature (see Table SM3 for a description of these). Each of these models makes different assumptions about how the rate of death and dispersal varies with age, but all can be represented by a general form,

$$S(t) = e^{-\int_0^t \lambda(\tau) \mathrm{d}\tau}. \tag{9}$$

| Survival function | Hazard rate | Interpretation | Mean time spent in study area, $\overline{T}$ | Papers assuming this function |
|---|---|---|---|---|
| Exponential | $\lambda$ | Constant mortality risk | $\frac{1}{\lambda}$ | Ross (1910); Anderson & May (1992); Smith & McKenzie (2004) |
| Gompertz | $\alpha e^{\beta t}$ | Mortality increases with age at an ever-increasing rate | $\frac{e^{\frac{\alpha}{\beta}}\Gamma(0,\frac{\alpha}{\beta})}{\beta}$ | Clements & Paterson (1981); Styer *et al.* (2007);Novoseltsev *et al.* (2012) |
| Weibull | $\alpha t^{\beta};\ \beta \geq 1$ | Mortality increases with age at an ever-increasing rate | $\alpha^{-\frac{1}{\beta}}\Gamma(1+\frac{1}{\beta})$ | Hancock *et al.* (2009); and Carey (2001) considering general insect demography |
| Gompertz-Makeham | $\alpha e^{\beta t} + c$ | Two additive mortality risks: one that increases with age, and another that is age-independent | No simple analytic form | Styer *et al.* (2007) |
| Logistic | $\frac{\alpha e^{\beta t}}{1+\frac{\alpha s}{b}(e^{bt}-1)}$ | Mortality risk increases with age, although at a declining rate | No simple analytic form | Styer *et al.* (2007); Novoseltsev *et al.* (2012) |
| Logistic-Makeham | $\frac{\alpha e^{\beta t}}{1+\frac{\alpha s}{b}(e^{bt}-1)} + c$ | Two separate additive mortality risks: an age-dependent risk of the same form as the logistic model, and a constant hazard | No simple analytic form | Styer *et al.* (2007); Bellan (2010) |

Table SM3: **A description of the survival functions used in this study, arranged in rough order of model complexity (simple-complex from top-bottom.)** All parameters are defined to be non-negative. $\Gamma(\theta)$ and $\Gamma(\theta_1,\theta_2)$ refer to the Euler gamma function, and incomplete gamma function respectively. The 'mean time spent in study area' is an estimate of the combined effects of mosquito mortality and dispersal from the area of the study where collections take place, since we do not analyse spatially-resolved data.

where we constrain the parameters of our models to preclude the possibility of a hazard that decreases with age ($\frac{\mathrm{d}\lambda(t)}{\mathrm{d}t} \geq 0$). Whilst this could occur if older mosquitoes disperse less than younger mosquitoes, it is unlikely this would outweigh any declines in survival associated with old age.

The parameters of the survival function of each model were assigned hierarchical priors that allowed considerable variation in mosquito lifespan (see Fig. SM6) but had similar means as to the constant mortality case. Otherwise, the statistical model we used remained the same as for the constant mortality case. In testing for age-dependent mortality, we did not account for sex or pre-release feeding effects due to the complexity of including these elements in more complex survival models.

| Survival function | time-series-level priors | Group-level priors |
|---|---|---|
| Exponential | $c \sim \mathcal{N}(\mu_\lambda, \sigma)$ | $\mu_\lambda \sim N(-2.32, 1)$, $\sigma \sim$ log-normal$(-3, 1)$ |
| Gompertz | $\alpha, \beta \sim$ log-normal$(\mu_{\alpha\|\beta}, 0.2)$ | $\mu_\alpha \sim N(-3, 1)$, $\mu_\beta \sim N(-3, 1.1)$ |
| Weibull | $\alpha, (\beta - 1) \sim$ log-normal$(\mu_{\alpha\|\beta}, 0.2)$ | $\mu_\alpha \sim N(-4.8, 1.75)$, $\mu_\beta \sim N(-4, 2)$ |
| Gompertz-Makeham | $\alpha, \beta, c \sim$ log-normal$(\mu_{\alpha\|\beta\|c}, 0.2)$ | $\mu_\alpha \sim N(-3, 1)$, $\mu_\beta \sim N(-3, 0.5)$, $\mu_c \sim N(-4.5, 0.5)$ |
| Logistic | $\alpha, \beta, s \sim$ log-normal$(\mu_{\alpha\|\beta\|s}, 0.2)$ | $\mu_\alpha \sim N(-2.8, 1)$, $\mu_\beta \sim N(-3, 1)$, $\mu_s \sim N(-3, 1)$ |
| Logistic-Makeham | $\alpha, \beta, s, c \sim$ log-normal$(\mu_{\alpha\|\beta\|s\|c}, 0.2)$ | $\mu_\alpha \sim N(-3.2, 1)$, $\mu_\beta \sim N(-3.2, 1)$, $\mu_s \sim N(-3, 0.5)$, $\mu_c \sim N(-4, 0.5)$ |

Table SM4: **MRR: priors used on parameters of each survival model.** For the exponential model, the 'group-level' priors were the same for the genus and 'overall' models that were also estimated. The exponential model was the only model that was simple enough to allow the scale parameter of the log-normal ($\sigma$) to be estimated by the data. The notation $\alpha, \beta \sim$ log-normal$(\mu_{\alpha|\beta}, 0.2)$ means that $\alpha$ and $\beta$ were assigned independent log-normal priors with location parameters $\mu_\alpha$ and $\mu_\beta$ respectively, and a scale parameter of 0.2 in both cases.

#### SM1.7 Model estimation by MCMC

The likelihood and priors used result in posterior distributions whose analytic form cannot be calculated with existent computational methods. Instead, we use Markov Chain Monte Carlo (MCMC) methods to sample from each posterior distribution. Specifically, we used *Stan* software (Carpenter *et al.* , 2016) that implements an efficient MCMC algorithm known as NUTS (Hoffman & Gelman, 2014).

To judge convergence of the sampling algorithm to the posterior density, we calculated $\hat{R}$ across all Markov chains – a ratio that compares between-chain variation to that within each chain that is commonly used to measure convergence in MCMC (Gelman & Rubin, 1992). For each class of model, we used the following MCMC parameters,

- Individual lifespan models (Section SM1.4): 12 independent chains with 1000 iterations per chain,
- Hierarchical lifespan estimates (Section SM1.5): 12 independent chains with 15,000 iterations per chain,
- Age-dependent analysis (Section SM1.6): 12 independent chains with 3000 iterations per chain.

In all cases, we discarded the first half of these iterations as warm-up (Gelman *et al.* , 2014). At the end of all runs for lifespan estimation, $\hat{R} < 1.1$ for all model parameters. For the age-dependent cases, there remained a handful of parameters where $1.2 > \hat{R} > 1.1$ after running simulations for 3000 iterations but this did not affect our inferences, since repeated runs demonstrated the same pattern. We also ensured that across each MCMC run, the number of divergent iterations (that can bias the MCMC away from the true posterior density) was minimal. In the majority of cases, the number of divergent iterations was far fewer than 1% of the total number of samples.

The Stan scripts for lifespan estimation (i.e. those using an exponential survival model) for the MRR analyses are provided below. The script for the individual time series analysis (Section SM1.4):

```
data {
// Single release parameters
int<lower=0> N; // number of observations
int<lower=0> K; // number of groups
int Y[N]; // observations for the releases
int S[K]; // time series lengths
vector[N] t; // time observations (days)
int Pos[K]; // starting position of each dataset

// Multiple release parameters
int RelFreq[K]; // frequency of releases in each study
// starting position of releases in each study
// in RelNumber
int PosRel[K];
int<lower=0> NReleases; // total number of releases
// numbers released in each study
vector[NReleases] RelNumber;
// time of each release in each study
vector[NReleases] RelTime;

real Age[K];
}

parameters {
real<lower=0.3, upper=20> kappa[K];
real<lower=0> cDecay[K];
real<lower=0, upper=1> psi[K];
```

```
}

model {
// Likelihood
for (i in 1:K) {
real lambda[S[i]];

if (RelFreq[i]< 2) // Single release
{
   real numberCaught;
   numberCaught = 0;
   for (j in 1:S[i])
   {
     lambda[j]= (psi[i] *
                 (RelNumber[PosRel[i]] - numberCaught) *
                 exp(-cDecay[i] * (t[Pos[i] + j - 1] + Age[i])));
     //1e-5 to prevent loc=0
     Y[Pos[i] + j - 1] ~ neg_binomial_2(
                           lambda[j] + 1e-5, kappa[i]);
     numberCaught = numberCaught + Y[Pos[i] + j - 1];
   }
} else // Multiple release
  {
    for (j in 1:S[i])
    {
      real lambdaTemp;
      lambdaTemp = 0;
      for (kk in 1:RelFreq[i])
      {
        if (t[Pos[i] + j - 1]> RelTime[PosRel[i]+kk-1])
        {
           lambdaTemp = (
                   lambdaTemp +
                   RelNumber[PosRel[i] + kk - 1] *
                   ((1 - psi[i])^(t[Pos[i] + j - 1] -
                   RelTime[PosRel[i]+kk-1]-1)) *
                   psi[i] * exp(-cDecay[i] *
                   (t[Pos[i] + j - 1] -
                   RelTime[PosRel[i]+kk-1] + Age[i]))
           );
```

```
          }
    }
    lambda[j] = lambdaTemp;
    Y[Pos[i] + j - 1] ~ neg_binomial_2(lambda[j]+1e-5, kappa[i]);
    }
  }
}

// Priors
for (i in 1:K)
{
    cDecay[i] ~ lognormal(-2.32, 1);
    kappa[i] ~ lognormal(2, 1);
    psi[i] ~ exponential(50);
}
}

generated quantities {
vector[K] lifespans;
  for(i in 1:K)
    lifespans[i] = 1 / cDecay[i];
}
```

The script for the hierarchical lifespan estimation (Section SM1.5) is shown below:

```
data {
// Single release parameters
int<lower=0> N; // number of observations
int<lower=0> K; // number of groups
int Y[N]; // observations
int S[K]; // group sizes
vector[N] t; // time observations (days)
int Pos[K]; // starting position of each dataset

// Multiple release parameters
int RelFreq[K]; // release frequency in each study
int PosRel[K]; // starting position of releases in each study
int<lower=0> NReleases; // release frequencies
vector[NReleases] RelNumber; // numbers released
vector[NReleases] RelTime; // release times
```

```
int<lower=0> nSpecies; // number of individual species
int SpeciesIndex[K]; // species index of each study

real Age[K];
int SexM[K];
int SexMix[K];
int Genus[K];
int Blood[K];
int Sugar[K];
}

transformed data{
int SpeciesToGenus[nSpecies];
for(i in 1:K){
  SpeciesToGenus[SpeciesIndex[i]] = Genus[i];
}
}

parameters {
real<lower=0.3,upper=100> kappa[K];
real a[nSpecies];
real<lower=0> p[nSpecies];
real<lower=0> r[nSpecies];
real cDecay[K];
real<lower=0,upper=0.1> psi[K];
real<lower=0> sigma_a[nSpecies];
real delta_M[3];
real delta_Blood[3];
real delta_Sugar[3];
real<lower=0, upper=1> u_mixed[3];
}

transformed parameters{
real delta_Mixed[3];
for(i in 1:3)
  delta_Mixed[i] = u_mixed[i] * delta_M[i];
}

model {
```

```
for (i in 1:K) {
  real lambda[S[i]];
  real decay_temp = exp(cDecay[i]+delta_M[Genus[i]] * SexM[i] +
    delta_Mixed[Genus[i]] * SexMix[i] + delta_Sugar[Genus[i]] *
    Sugar[i] + delta_Blood[Genus[i]] * Blood[i]);

  if (RelFreq[i]< 2) // Single release
  {
    real R_times_psi;
    real numberCaught;
    numberCaught = 0;
    R_times_psi = psi[i];
    for (j in 1:S[i])
    {
    lambda[j]= R_times_psi *
            (RelNumber[PosRel[i]]-numberCaught) *
            exp(-decay_temp * (t[Pos[i] + j - 1] + Age[i]));
    Y[Pos[i] + j - 1] ~ neg_binomial_2(lambda[j] + 1e-5,
                                        kappa[i]);
    numberCaught = numberCaught + Y[Pos[i] + j - 1];
    }
  }
  else // Multiple release
  {
    for (j in 1:S[i])
    {
    real lambdaTemp;
    lambdaTemp = 0;
    for (kk in 1:RelFreq[i])
    {
      if (t[Pos[i] + j - 1]> RelTime[PosRel[i] + kk - 1])
      {
        lambdaTemp = (lambdaTemp +
          RelNumber[PosRel[i] + kk - 1] *
          ((1-psi[i])^(t[Pos[i] + j - 1] -
          RelTime[PosRel[i] + kk - 1] - 1)) *
          psi[i] * exp(-decay_temp *
          (t[Pos[i] + j - 1] -
          RelTime[PosRel[i] + kk - 1] + Age[i]));
         );
```

```
        }
      }
      lambda[j] = lambdaTemp;
      Y[Pos[i] + j - 1] ~ neg_binomial_2(lambda[j] + 1e-5,
                                          kappa[i]);
      }
    }
}

// Priors
for (i in 1:K)
{
  int speciesTemp;
  speciesTemp = SpeciesIndex[i];
  cDecay[i] ~ normal(a[speciesTemp], sigma_a[speciesTemp]);
  kappa[i] ~ exponential(p[speciesTemp]);
  psi[i] ~ exponential(r[speciesTemp]);
}
a ~ normal(-2.32, 1);
sigma_a ~ lognormal(-3, 1);
p ~ lognormal(1.5, 1);
r ~ gamma(100, 2);
delta_M ~ normal(0, 1);
delta_Blood ~ normal(0, 1);
delta_Sugar ~ normal(0, 1);
}

generated quantities{
real overallcDecay[nSpecies];
real overallFLife[nSpecies];
real overallMLife[nSpecies];
real overallMixedLife[nSpecies];
real overallFBloodLife[nSpecies];
real overallFBothLife[nSpecies];
real overallFSugarLife[nSpecies];
real overallMSugarLife[nSpecies];

for(i in 1:nSpecies){
  overallcDecay[i] = normal_rng(a[i], sigma_a[i]);
  overallFLife[i] = 1 / exp(overallcDecay[i]);
```

```
  overallMLife[i] = 1 / exp(overallcDecay[i] +
                            delta_M[SpeciesToGenus[i]]);
  overallMixedLife[i] = 1 / exp(overallcDecay[i] +
                            delta_Mixed[SpeciesToGenus[i]]);
  overallFBloodLife[i] = 1 / exp(overallcDecay[i] +
                             delta_Blood[SpeciesToGenus[i]]);
  overallFSugarLife[i] = 1 / exp(overallcDecay[i] +
                             delta_Sugar[SpeciesToGenus[i]]);
  overallFBothLife[i] = 1 / exp(overallcDecay[i] +
                             delta_Sugar[SpeciesToGenus[i]] +
                             delta_Blood[SpeciesToGenus[i]]);
  overallMSugarLife[i] = 1 / exp(overallcDecay[i] +
                             delta_M[SpeciesToGenus[i]] +
                             delta_Sugar[SpeciesToGenus[i]]);
}
}
```

##### SM1.8 K-Fold cross validation

One way to estimate a model's out-of-sample predictive capability is to use the Akaike Information Criterion that explicitly penalises a model in accordance to its complexity (typically indexed by its number of free parameters) to correct for model over-fit. The way in which this correction takes places however, is fairly heuristic, and less appropriate for hierarchical models. An alternative method, common in the machine learning literature, is known as 'cross-validation' (see for example, Kohavi *et al.* (1995)). Here to estimate out of sample predictive capability, the original data set is partitioned into training and test sets. The model is then fitted to the training set, and its predictive performance measured on the independent test set.

We use cross-validation to compare the predictive power of the species, genus and overall models, and later to determine whether age-dependent mortality occurs. In particular, we use K-Fold cross validation (with $K = 4$ folds for the MRR analysis), where we repeatedly partition our dataset into a test set composed of a number of individual time-series, and a training set of the remainder (see for example, Marsland (2015)). We then use the fitted model to measure the predictive performance on the test set. The particular measure we use is the expected log point-wise predictive density (elpd)(Vehtari *et al.* , 2015) which sums the predictive performance for each data point averaged across all posterior samples.

Since the models are hierarchical, we must specify how to draw parameters for a given test sample time-series. Here we chose to draw values of the individual time-series parameters as samples from the population distribution that corresponds to that particular sample's respective grouping. So if the particular test sample time-series pertains to *Ae. aegypti* (and we are using the species-level model), then we draw independent samples for the statistical model's parameters from the estimated *Ae. aegypti* population distribution,

$$\theta_i \sim p(\theta^{\text{Ae. aegypti}} | data), \tag{10}$$

where $\theta_i$ is the value of the parameters used in test time-series $i$, and $p(\theta^{\text{Ae. aegypti}} | data)$ is the overall posterior distribution across all *Ae. aegypti* experiments.

#### SM1.9 Model checking

To check the fit of the model to the data, we simulated from the posterior predictive distribution for each data point within each time series of recapture observations (see Figures SM8, SM9 & SM10 and the attached file, `mrr_ppcs_all.pdf` for the full graphs). In the majority of cases, the data lay within the 95% predictive intervals indicating that the model was a good fit to the data.

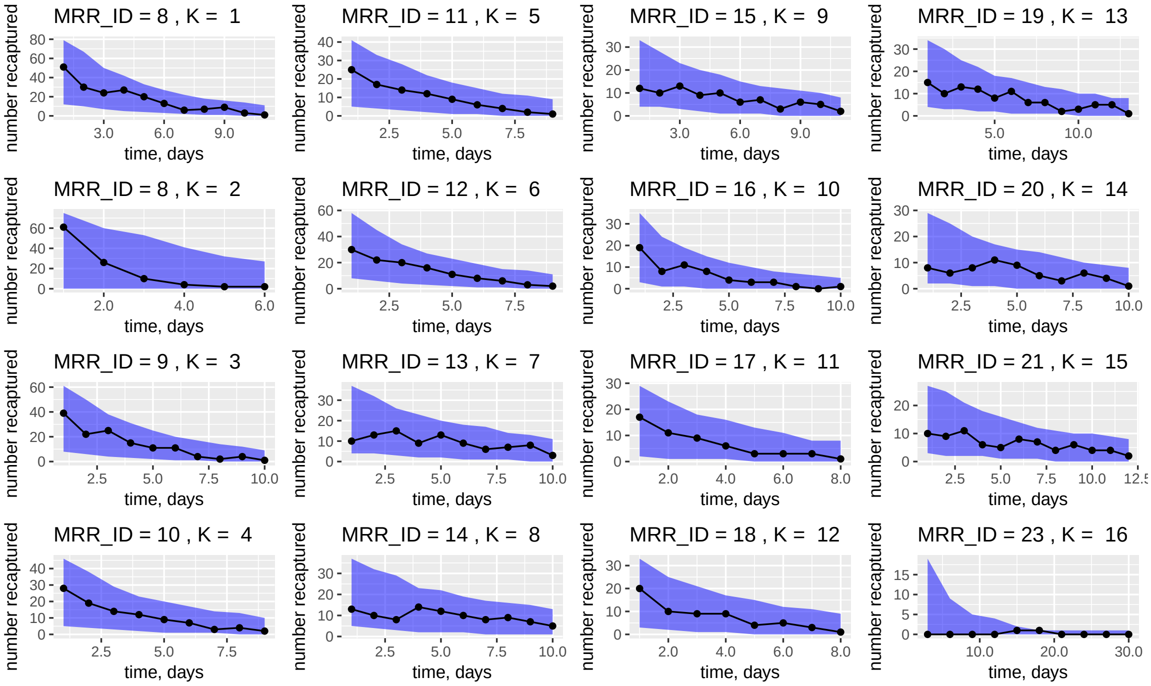


Figure SM8: **MRR: posterior predictive distribution versus actual data.** The figure shows the numbers of mosquitoes recaptured (black lines) versus the 95% central posterior interval of the posterior predictive distribution (blue shading) for a selection of the time series. For the rest of the posterior predictive checks for the MRR models, see the file referenced in the text.

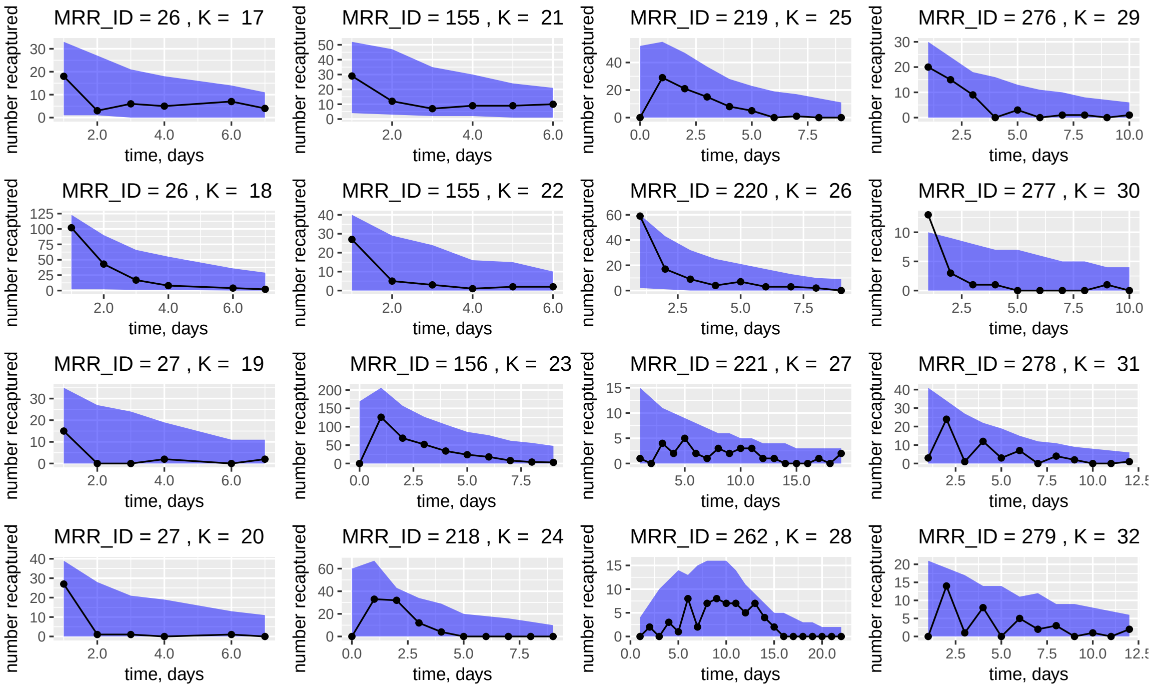


Figure SM9: **MRR: posterior predictive distribution versus actual data.** The figure shows the numbers of mosquitoes recaptured (black lines) versus the 95% central posterior interval of the posterior predictive distribution (blue shading) for a selection of the time series. For the rest of the posterior predictive checks for the MRR models, see the file referenced in the text.

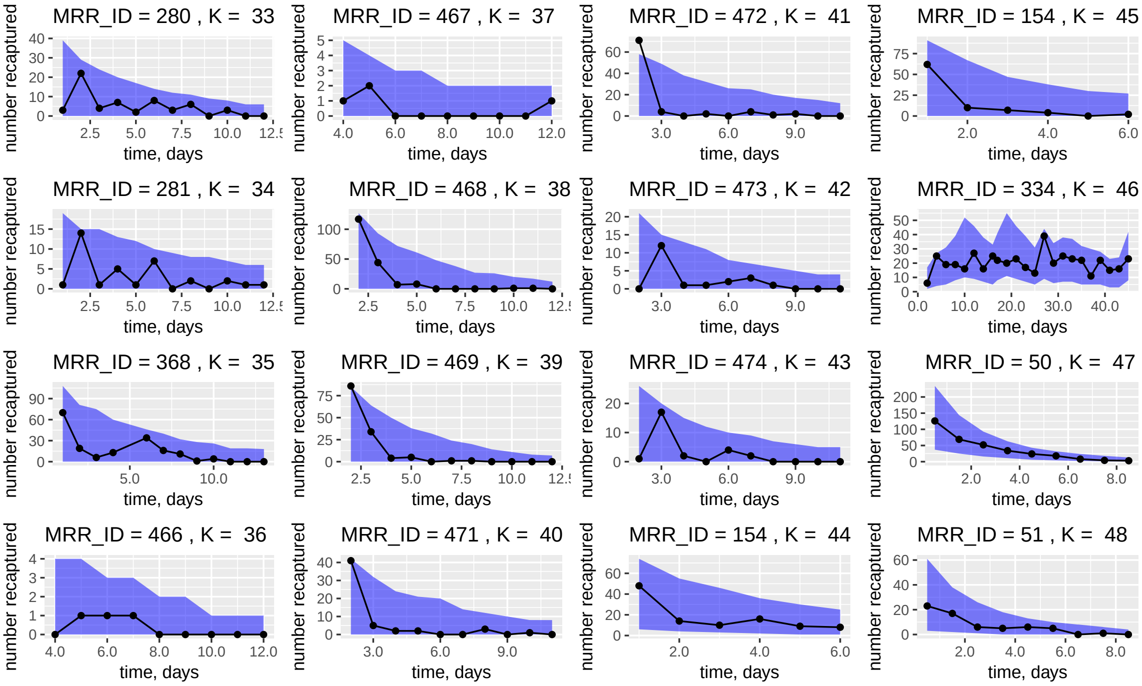


Figure SM10: **MRR: posterior predictive distribution versus actual data.** The figure shows the numbers of mosquitoes recaptured (black lines) versus the 95% central posterior interval of the posterior predictive distribution (blue shading) for a selection of the time series. For the rest of the posterior predictive checks for the MRR models, see the file referenced in the text.

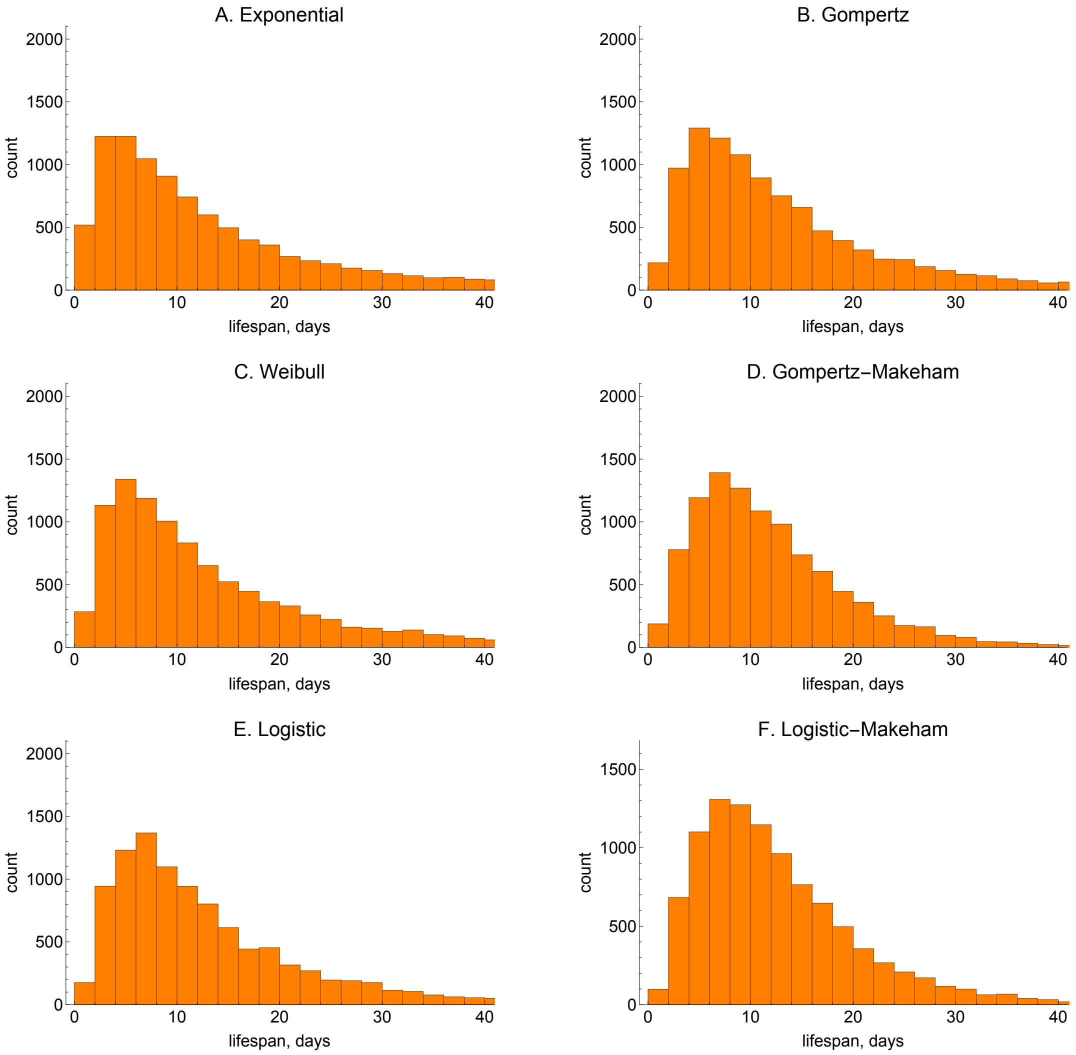


Figure SM6: **MRR: prior predictive distribution of mean lifespan across the six models of mosquito mortality.** In all cases, 10,000 samples were generated using the priors indicated in Table SM4, resulting in distributions with a mean close to 10 days.

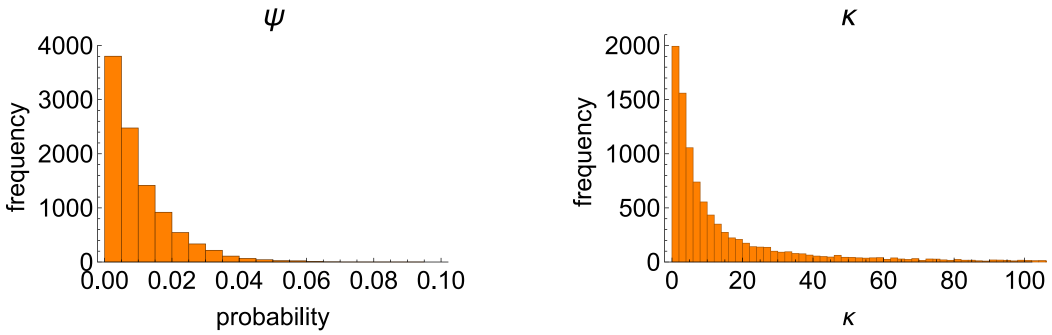


Figure SM7: **MRR: samples from the priors for daily recapture probability ($\psi$) and over-dispersion parameter ($\kappa$).** In both panels, 10,000 samples were generated using the following hierarchical priors: for the individual time series parameters, $\psi_i \sim \exp(r)$ (with $\psi$ constrained to be less than 0.1 - encompassing the upper limit of day one recapture fractions observed in most data series) and $\kappa_i \sim \exp(p)$; for the top-level parameters: $r \sim \text{gamma}(100, 2)$ and $p \sim \text{log-normal}(1.5, 1)$.

### SM2 Polovoda dissection experiments

#### SM2.1 Collection of dissection data on physiological age

A comprehensive search of the literature using Google Scholar (`scholar.google.co.uk`) was performed using various combinations of the following keywords: dissection, mosquito, parity, parous, age, and physiological age. No constraints were placed on publication date, location or type. This list was supplemented with a number of author-specific searches for those individuals most prevalent in the literature. In particular, we searched for all articles authored by: Charlwood, Muller, Schlein, Samarawickrema, Reisen, Detinova, Polodova, Gillies, and Wilkes. An additional list of potential articles was provided by doing a forward article search on some of the most highly-cited articles in the database: Polovodova (1949); Detinova *et al.* (1962); Gillies & Wilkes (1965); Clements & Paterson (1981). The list of results was then filtered manually by examining the titles and abstracts to produce a candidate list of the articles most likely to contain data on the physiological age of dissected mosquitoes caught in the wild, as determined by the Polovodova (1949) method.

A relational database was used to store the raw data from the actual experiments, along with the meta-data associated with each of the experiments. In many of the published articles, wild mosquitoes were caught and dissected over a period of time, and the raw data thus consists of snapshots of the age structure of the population at regular intervals in time. In the cases where the data was more sparse (fewer than ten individuals, on average, per date), we aggregated across dates and recorded this as a single entry; otherwise we recorded the snapshots of the population at each different date. We record separate series for each species that was captured, or for those that were recorded at separate capture locations (potentially with an alternative collection method), and do not aggregate over these datasets.

For each individual series, we recorded the following meta-data: genus, species, collection method and the start and end dates of the experiment. At the article level, we recorded the following meta-data: author, year, title, country, location (within country), start and end date, whether an MRR experiment occurred alongside the dissection study, and the collection location (indoor or outdoor). At either the individual series or article levels, we recorded additional meta-data describing the nature of data collection, for example explaining where the data was contained within the article, whether it was obtained by digitising graphs, and the number of separate dated series. For those few cases ($n = 16$ series) where the data was obtained by digitising graphs, we used the WebPlotDigitizer online tool (Rohatgi, 2017).

The data collection method resulted in 568 separate dissection series (see polovodova_full.csv), across 73 published articles (see parity_papers_summary.csv). The published datasets cover the period from 1960-2015 with comparable numbers of studies across each decade (Fig. SM11). The statistical approach applied to the data relies on the assumption that there is a constant rate of recruitment into the adult population (see Section SM2.2). If populations fluctuate strongly from month-to-month, this assumption will likely be violated. To try to mitigate against such a risk, we aggregated the data across all dates to obtain a single series for each identifier, resulting in 201 such series (see polovodova_aggregated.csv). To obtain a reasonable level of accuracy on estimates, we removed all those individual (aggregated) series where fewer than 100 mosquitoes were collected. Finally, we remove data for any species where there was only one series resulting in 131 series across four genera (*Anopheles*, *Culex*, *Aedes*, and *Mansonia*) and 25 species (Table SM5). These studies are distributed across a wide range of geographies (Fig. SM12). The raw data for the dissection analysis are shown in Figures SM13, SM14 and SM15.

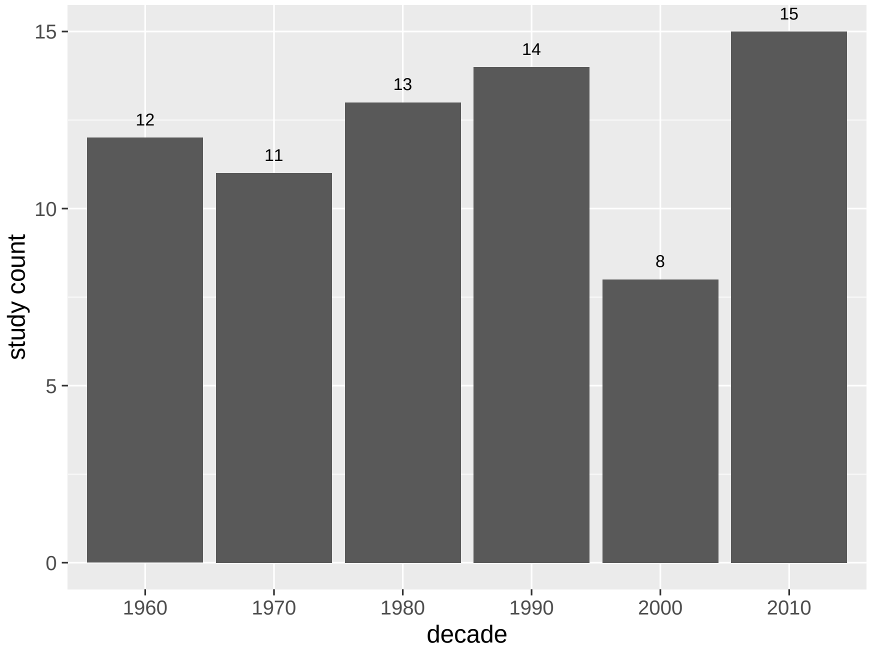


Figure SM11: **Polovodova dissection: numbers of published studies that estimate gonotrophic age by this method.**

| **Genus** | **Species** | **Frequency** |
|---|---|---|
| *Anopheles* | *gambiae s.l.* | 19 |
| *Culex* | *quinquefasciatus* | 12 |
| *Anopheles* | *maculipennis* | 11 |
| *Anopheles* | *farauti s.l.* | 10 |
| *Anopheles* | *sergentii* | 8 |
| *Aedes* | *polynesiensis* | 8 |
| *Culex* | *pipiens* | 6 |
| *Anopheles* | *culicifacies* | 5 |
| *Anopheles* | *darlingi* | 5 |
| *Anopheles* | *quadrimaculatus* | 5 |
| *Anopheles* | *stephensi* | 4 |
| *Anopheles* | *melas* | 4 |
| *Culex* | *annulirostris* | 4 |
| *Aedes* | *aegypti* | 3 |
| *Aedes* | *samoanus* | 3 |
| *Anopheles* | *minimus* | 3 |
| *Anopheles* | *rivulorum* | 3 |
| *Culex* | *thalassius* | 3 |
| *Mansonia* | *uniformis* | 3 |
| *Anopheles* | *subpictus* | 2 |
| *Aedes* | *sollicitans* | 2 |
| *Aedes* | *vexans* | 2 |
| *Anopheles* | *bellator* | 2 |
| *Anopheles* | *cruzii* | 2 |
| *Culex* | *tritaeniorhynchus* | 2 |
| ***Anopheles*** | | **83** |
| ***Culex*** | | **27** |
| ***Aedes*** | | **18** |
| ***Mansonia*** | | **3** |
| **Total** | | **131** |

Table SM5: **Polovodova dissection: numbers of dissection series included in the overall dataset.** Note, these counts represent numbers of series analysed by the hierarchical models – that is, after aggregating the individual series as described in Section SM2.1.

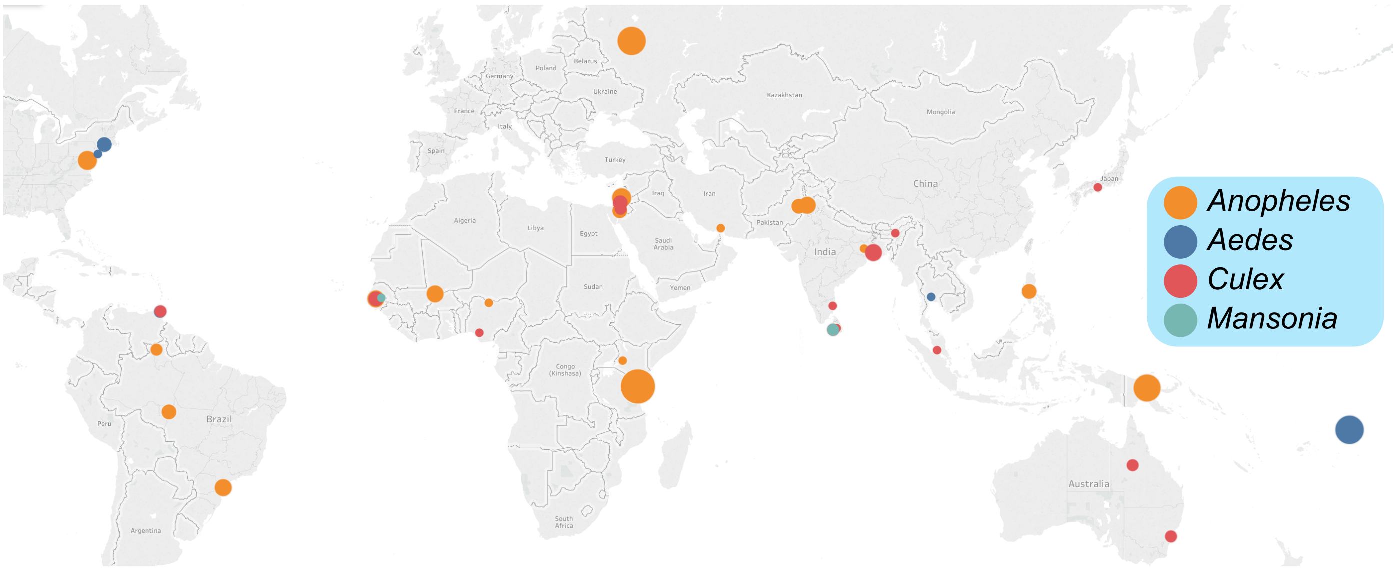


Figure SM12: **Polovodova dissection: the geographic location of each of the datasets included in the meta-analysis.** The area of the bubbles indicates the number of unique data series available.

#### SM2.2 Statistical analysis of Polovodova dissection data

We suppose that the number of individuals recruited into the adult female population is constant over time, meaning that the age-structure of the population is stable. Whilst environmental heterogeneity will naturally lead to variation in adult recruitment over time, we hope that by aggregating data across studies undertaken over a range of dates this will reduce the impact of this effect. Suppose that the probability an individual female mosquito survives until age $a$ is given by the survival function $S(a)$, then the number of individuals in the population

| **Variable** | **Min** | **Median** | **Mean** | **Max** | **Standard deviation** |
|---|---|---|---|---|---|
| Min age of captures, gonotrophic cycles | 0 | 0 | 0.02 | 1.00 | 0.12 |
| Median age of captures, gonotrophic cycles | 0 | 1.00 | 0.63 | 2.00 | 0.62 |
| Mean age of captures, gonotrophic cycles | 0.17 | 0.90 | 1.00 | 3.03 | 0.58 |
| Max age of captures, gonotrophic cycles | 1.00 | 5.00 | 5.77 | 13.00 | 2.94 |
| Number captured | 100 | 565 | 1,317 | 14,012 | 1,885 |

Table SM6: **Polovodova dissection: summaries of the data series included in this analysis.** Note when series are censored (see Section SM2.2.1) we assume that all mosquitoes dissected over the threshold are of this age when calculating summary statistics.

surviving to this age is given by,

$$A(a) = A(0)S(a), \tag{11}$$

where $A(0)$ is the number of adult female mosquitoes recruited into the population per unit time. Consider one individual experiment where we randomly sample from a wild population that is structured as per eq. (11). In this case, the number of individuals sampled at each age is binomially-distributed,

$$y(a) \sim \mathcal{B}(A(a), p(a)), \tag{12}$$

where $p(a)$ is the probability of recapturing a single mosquito of age $a$. In what follows, we assume that the probability of capturing a given mosquito is independent of their age, so that $p(a) = p = const.$ Since, in general, $A(a)$ is large and $p(a)$ is small, we can approximate the above using a Poisson distribution,

$$y(a) \sim \text{Poisson}(A(a)p). \tag{13}$$

However, the assumption of *independent* captures of individual mosquitoes, which underlies the binomial and Poisson models, is likely suspect for the same reasons as for the MRR analysis (see Section SM1.3). As before, we instead use a negative binomial sampling distribution that allows for non-independent captures,

$$y(a) \sim \text{NB}(A(a)p, \kappa), \tag{14}$$

where we use the parameterisation such that the mean is $A(a)p$, and over-dispersion parameter is $\kappa$, where, as $\kappa \to \infty$, the above sampling distribution approaches a Poisson.
In the field, unfortunately, we do not know the number of mosquitoes recruited into the adult population, $A(a)$, nor the probability of capturing an individual at a given point in time, $p$. Instead, we model their product $\Psi = A(0)p$ (the population of adult female mosquitoes of age zero that are expected to be captured) probabilistically resulting in a model,

$$y(a) \sim \text{NB}(\Psi S(a), \kappa). \tag{15}$$

The resultant likelihood of a data series consisting of counts: $(y(a_1), y(a_2), ..., y(a_R))$, is then calculated by assuming (conditional) independence of the observations,

$$\mathcal{L}(y(a_1), y(a_2), ..., y(a_R)|S(.), \Psi, \kappa) = \prod_{a=a_1}^{a_R} p(y(a)|S(a), \Psi, \kappa), \tag{16}$$

where $p(y(a)|S(a), \Psi, \kappa)$ corresponds to the negative binomial probability mass function for a count of $y(a)$ mosquitoes aged $a$ as specified in eqn. (15), and $R$ is the number of separate physiological age classes that we include: for series with no censoring (see below for how censored data were handled), we include 14 separate age classes (from nulliparous to 13-parous). We recognise that the choice of a "maximum" age class is an approximation, since, in theory, we should include a contribution from each "0" observed until $\infty$. In reality, however, the limit was chosen to match the maximum number of cycles observed in the data, and the majority of series had far maxima that were less than 5. As such, we believe that our truncation should provide a good approximation to the infinite sum. Since we do not know $\Psi$, we must learn it from the data. One approach to estimate this parameter could be use the number of captures of nulliparous mosquitoes, $X(0)$ in place of $\Psi$. We prefer to allow for some uncertainty in this parameter and use the data to estimate its value. However, unfortunately, the negative binomial likelihood allows too much variation in this parameter (because the data are over-dispersed), and instead we specify a likelihood of the form,

$$X(0) \sim \mathcal{N}(\Psi, \sqrt{\Psi}), \tag{17}$$

solely for the first data point $X(0)$. The above allows for some freedom in $\Psi$ whilst ensuring that the parameter's probability mass lies near enough to $X(0)$ to allow useful model estimates. $\Psi$ is set a uniform prior over the range $[0, \frac{3}{2}X(0)]$.

Since $S(a)$ is monotonically-decreasing, we know that, if recruitment to the adult population is constant, then the numbers of nulliparous individuals should exceed the count in subsequent parity states. However, in a number of data series there is a relative dearth of nulliparous mosquitoes versus uniparous individuals. This deficiency has been noted in a previously-published study where the authors hypothesise that it is due to the issue of sampling the nulliparous population, since they are more likely to rest outside, compared with parous individuals (Gillies & Wilkes, 1965). However, there is conflicting evidence that suggests that, on average, the first gonotrophic cycle is longer than subsequent cycles (see Section SM2.5), meaning that a relative surplus of nulliparous mosquitoes may exist in captured samples (Clements & Paterson, 1981).

In our analysis, we chose to remove the counts of nulliparous individuals from the series where the count was low relative to uniparous or higher parity individuals since this may indicate issues with sampling the nulliparous population. Specifically, we stipulated that the nulliparous count should exceed 90% of the count for the uniparous individuals. For those cases where this condition was not met, we removed the nulliparous observation and analyse the series of counts for all subsequent pars (uniparous and subsequent parous states).

Here, we estimate our model with one of six different survival functions (the same as for the MRR case; Table SM3), each of which makes different assumptions

| Survival function | Physiological age series-level priors | Group-level priors |
|---|---|---|
| Exponential | $\lambda \sim \text{log-normal}(\mu_\lambda, \sigma)$ | $\mu_\lambda \sim N(-1.2, 1)$, $\sigma \sim \text{log-normal}(-2, 1)$ |
| Gompertz | $\alpha, \beta \sim \text{log-normal}(\mu_{\alpha\|\beta}, 0.2)$ | $\mu_\alpha \sim N(-1.5, 1)$, $\mu_\beta \sim N(-2.5, 0.5)$ |
| Weibull | $\alpha, (\beta - 1) \sim \text{log-normal}(\mu_{\alpha\|\beta}, 0.2)$ | $\mu_\alpha \sim N(-2.5, 1)$, $\mu_\beta \sim N(-4, 0.5)$ |
| Gompertz-Makeham | $\alpha, \beta, c \sim \text{log-normal}(\mu_{\alpha\|\beta\|c}, 0.2)$ | $\mu_\alpha \sim N(-1.4, 1)$, $\mu_\beta \sim N(-3, 0.4)$, $\mu_c \sim N(-4.5, 0.5)$ |
| Logistic | $\alpha, \beta, s \sim \text{log-normal}(\mu_{\alpha\|\beta\|s}, 0.2)$ | $\mu_\alpha \sim N(-1.4, 1)$, $\mu_\beta \sim N(-2, 1)$, $\mu_s \sim N(-3, 1)$ |
| Logistic-Makeham | $\alpha, \beta, s, c \sim \text{log-normal}(\mu_{\alpha\|\beta\|s\|c}, 0.2)$ | $\mu_\alpha \sim N(-3, 1)$, $\mu_\beta \sim N(-1.3, 1)$, $\mu_s \sim N(-2, 0.5)$, $\mu_c \sim N(-4, 0.5)$ |

Table SM7: **Polovodova dissection: priors used on parameters of each different survival model.** For the exponential model the 'group-level' priors were the same for the genus and 'overall' models that were also estimated. The exponential model was the only model that was simple enough to allow the scale parameter of the log-normal ($\sigma$) to be estimated by the data. The notation $\alpha, \beta \sim \text{log-normal}(\mu_{\alpha|\beta}, 0.2)$ means that $\alpha$ and $\beta$ were assigned independent log-normal priors with location parameters $\mu_\alpha$ and $\mu_\beta$ respectively, and a scale parameter of 0.2 in both cases.

regarding how the force of mortality is affected by mosquito age. To estimate lifespan at the species, genus and overall groupings we then use a hierarchical Bayesian model of the same mathematical form as for the analysis of MRR experiments (i.e. assuming exponential survival functions; see Section SM1.5). However, the priors for the analysis of dissection data were modified so that they represented a mean prior lifespan of three gonotrophic cycles, although allowed considerable variation in this parameter (Fig. SM16). The priors used for the over-dispersion parameter ($\kappa$) for the negative binomial likelihood were the same as for the MRR analysis.

To determine whether mosquitoes experience age-dependent mortality, we compared the predictive power of each of the models that incorporate a hazard function that increases with age with that from the exponential model. As for the MRR analysis, we also used K-Fold cross-validation to perform this comparison, where the data are randomly partitioned into training and test sets (see Section SM1.8). The model is then fitted to each training set and used to predict the data in the independent test set. Since there are fewer series than for the MRR dataset, we used two partitions for each species, where each partition had roughly the same count of individual series.

The estimates of mean mosquito lifespan that we present here resulted from the use of the exponential survival model (i.e. no age dependence). This choice was

made because we found limited evidence in support of age-dependent mortality (see 'Results'). However, since the priors for each of the survival functions were specified to allow for a wide variety of lifespans (Fig. SM16), the specific choice of survival function made little difference to resultant estimates (data not shown).
To demonstrate that the results we obtained are not sensitive to the particular form of hierarchical prior structure chosen, we also provide estimates of the lifespan for each series analysed separately (i.e. non-hierarchically). To do so, we assume priors on the rate parameter of the exponential distribution that provides support over a wide range of possible lifespans (Fig. SM17A) but had a mean close to the hierarchical case equivalent. We also specified a prior on the over-dispersion parameter $\kappa$ that was comparable to the hierarchical case (Fig. SM17B).

##### SM2.2.1 Analysis of censored data series

Some published studies do not distinguish the number of gonotrophic studies beyond a threshold, $a_T$ (the more ovariole dilations there are the harder it is to count them) which is akin to censoring the data. This censoring involves only a relatively small number of mosquitoes in each time series (median = 2%) and so we are confident that these few cases do not substantially affect our inferences. We now describe the statistical model used to describe these instances. In these cases, we only know the number, $y(a_T)$, of individuals who were captured and dissected with an estimated physiological age that is greater than or equal to the threshold. Since we do not know the estimated physiological age of individual specimens $i$, we represent this as a parameter, $\aleph_i$, in our statistical model. This means that the joint distribution is a function of $y(a_T)$ different $\aleph_i$ parameters. Since these parameters are not directly of interest, and our chosen MCMC engine, Stan (Stan Development Team, 2014), does not directly allow discrete parameters in models, we marginalise these out of the joint distribution. To do this exactly would require an infinite sum over all the possible ages for all $y(a_T)$ mosquitoes that have been caught whose age exceeds the threshold. Rather than carry out this intractable number of summations, we instead make the approximation that all mosquitoes in this group are of the same age $\aleph_i = \aleph$, $\forall i \in (1, ..., y(a_T))$. We believe this assumption is justifiable, particularly since the numbers of mosquitoes in each subsequent age category is a strongly-decreasing function of age. This means that whilst we do not know with certainty individuals' ages, it is likely that most will be of the threshold age $a_T$.
This approximation means that to marginalise the parameter $\aleph$ out of the joint distribution, we are only required to do a single summation. Specifically if we have $y(a_T)$ captured individuals of age equal to, or exceeding, some threshold age, $a_T$,

the probability of these observations is given by,

$$q(y(a_T)|S(.), \Psi, \kappa) = \sum_{\aleph=a_T}^{\infty} p(y(a_T), \aleph|S(\aleph), \Psi, \kappa) \tag{18}$$

$$= \sum_{\aleph=a_T}^{\infty} p(y(a_T)|S(\aleph), \Psi, \kappa) \times p(\aleph) \tag{19}$$

where $q(y(a_T)|S(\aleph), \Psi, \kappa)$ is the probability of observing $y(a_T)$ counts for mosquitoes of an age $\aleph$. In practice, since we do not believe mosquitoes live for longer than 20 gonotrophic cycles (the maximum observed in the data was 13), we cut-off the summation at this point, and assume that the discrete prior probability distribution $p(\aleph)$ is uniform over this range. The overall likelihood for the cases where the series are censored therefore has the form,

$$\mathcal{L}(y(a_1), y(a_2), ..., y(a_T{-}1), y(a_T)|S(.), \Psi, \kappa) = \left(\prod_{a=a_1}^{a_T-1} p(y(a)|S(a), \Psi, \kappa)\right) q(y(a_T)|S(\aleph), \Psi, \kappa). \tag{20}$$

In practice, the number of mosquitoes captured in those series that are censored represents a small percentage of total captures, so the effect of the $q(.)$ term above is likely minimal on resultant inferences.

#### SM2.3 Model estimation by MCMC

As for the MRR analysis, we used *Stan* software (Carpenter *et al.* , 2016) to sample from the posterior distributions of model parameters. To judge convergence of the sampling algorithm to the posterior density, we calculated $\hat{R}$ across all Markov chains (Gelman & Rubin, 1992). For each model, we ran 16 independent Markov chains with 200 iterations per chain, discarding the first half of these iterations as warm-up (Gelman *et al.* , 2014). After running the algorithm for the given number of observations, $\hat{R} < 1.1$ for all model parameters. We also ensured that across each MCMC run, the number of divergent iterations (that can bias the MCMC away from the true posterior density) was minimal. In the majority of cases, the number of divergent iterations was far fewer than 1% of the total number of samples.

The model used to estimate the lifespan in terms of gonotrophic cycles is provided below.

```
data{
  int N; // number of obs
  int K; // number of time series
  int Y[N]; // number captured
  int S[K]; // length of each series
  int threshold[K]; // age of censoring; if -1 then no censoring
  int Pos[K]; // position of start of each series in Y
  int species[K]; // index variable of species
  int nSpecies; // number of species
}

parameters{
  real<lower=0> PopSize[K];
  real<lower=0> alpha[K];
  real alpha_mean[nSpecies];
  real<lower=0> alpha_sigma[nSpecies];
  real<lower=0> kappa[K];
  real<lower=0> p[nSpecies];
}

model{
// Likelihood
for(i in 1:K){
  for(t in 2:S[i]){
    // uncensored
    if(threshold[i] < 0)
      Y[Pos[i]+t-1] ~ neg_binomial_2(PopSize[i] *
                        exp(-alpha[i] * (t-1)), kappa[i]);
    // censoring
    else{
      if((t-1) < threshold[i])
        Y[Pos[i]+t-1] ~ neg_binomial_2(PopSize[i] *
                        exp(-alpha[i] * (t-1)), kappa[i]);
      else{
        real lLogProb[20];
        for(j in 1:20){
          lLogProb[j] = neg_binomial_2_lpmf(Y[Pos[i]+t-1] |
                        PopSize[i] * exp(-alpha[i] * (t-1)),
                        kappa[i]);
        }
        target += log_sum_exp(lLogProb);
      }
    }
  }
}
```

```
// Priors
for(i in 1:K){
  int aSpecies;
  aSpecies = species[i];
  Y[Pos[i]] ~ normal(PopSize[i], sqrt(PopSize[i]));
  alpha[i] ~ lognormal(alpha_mean[aSpecies],
                       alpha_sigma[aSpecies]);
  kappa[i] ~ exponential(p[aSpecies]);
}

alpha_mean ~ normal(-1.2, 1);
alpha_sigma ~ lognormal(-2, 1);
p ~ lognormal(1, 1);
}

generated quantities{
real alpha_average[nSpecies];
for(i in 1:nSpecies)
  alpha_average[i] = lognormal_rng(alpha_mean[i],alpha_sigma[i]);
}
```

#### SM2.4 Model checking

As for the MRR experiments, we simulated from the posterior predictive distribution for each series of dissection data. In the majority of the cases, the capture data lay within the 95% posterior predictive intervals, indicating that our model was a good fit to the data (see Figures SM18, SM19 & SM20 and the attached file, `dissection_ppcs_all.pdf` for the full graphs).

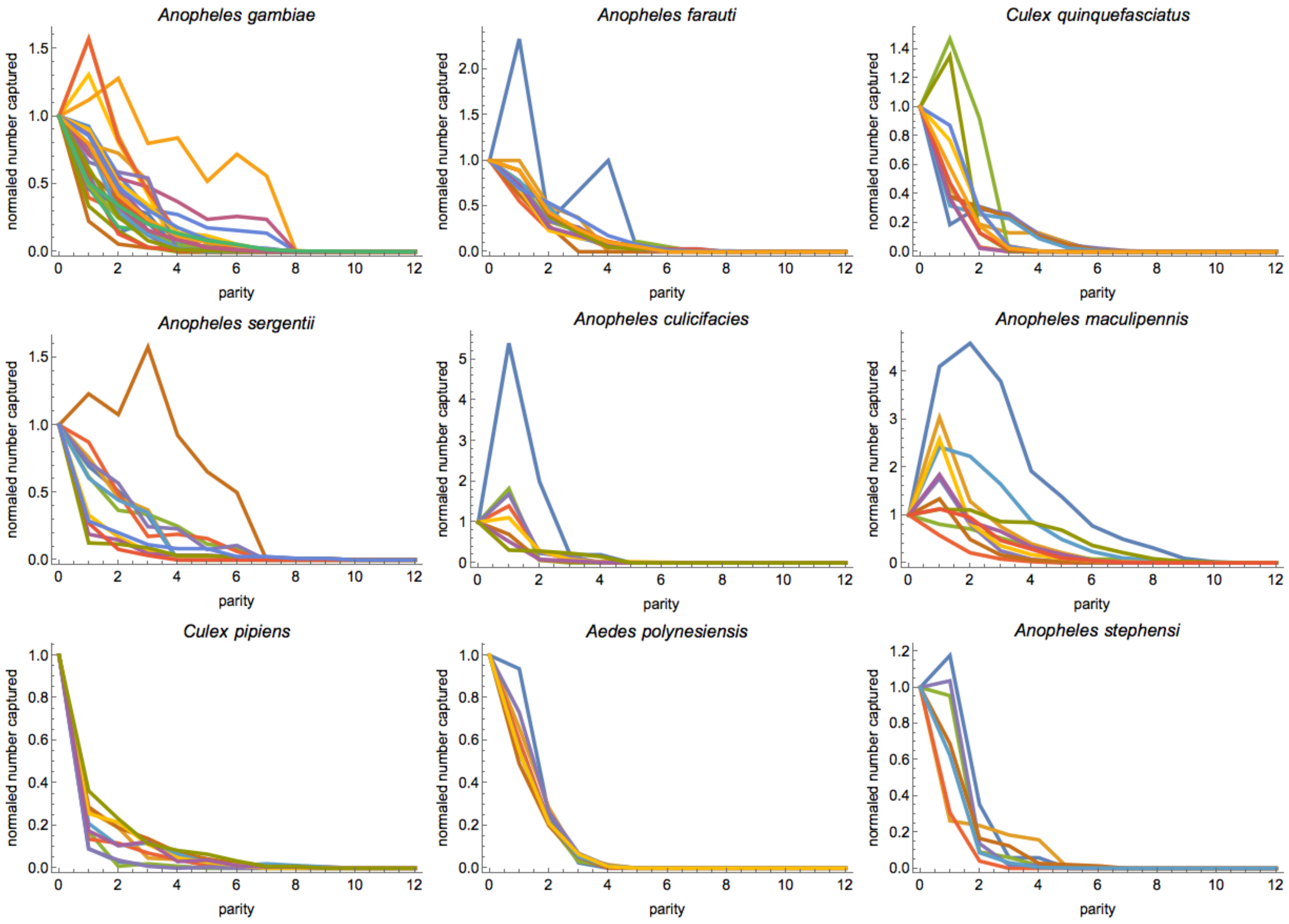


Figure SM13: **Polovodova dissection: normalised physiological age series for nine species in the database.** Each different coloured line represents an individual series. In each cas,e the count for all ages has been normalised by the nulliparous count. In all cases, we do not include any data for censored observations (see Section SM2.2.1).

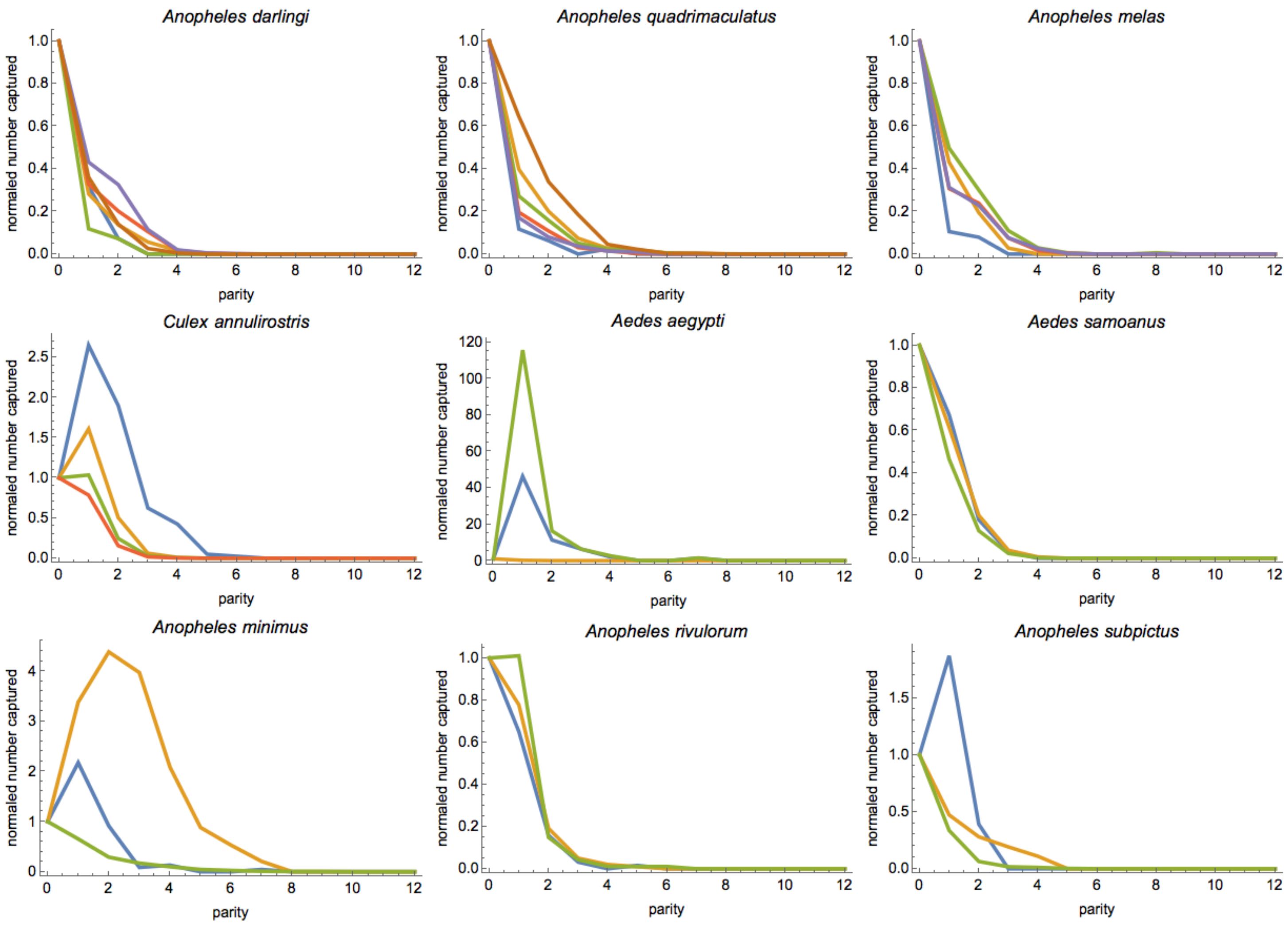


Figure SM14: **Polovodova dissection: normalised physiological age series for nine species in the database.** Each different coloured line represents an individual series. In each case, the count for all ages has been normalised by the nulliparous count. In all cases, we do not include any data for censored observations (see Section SM2.2.1).

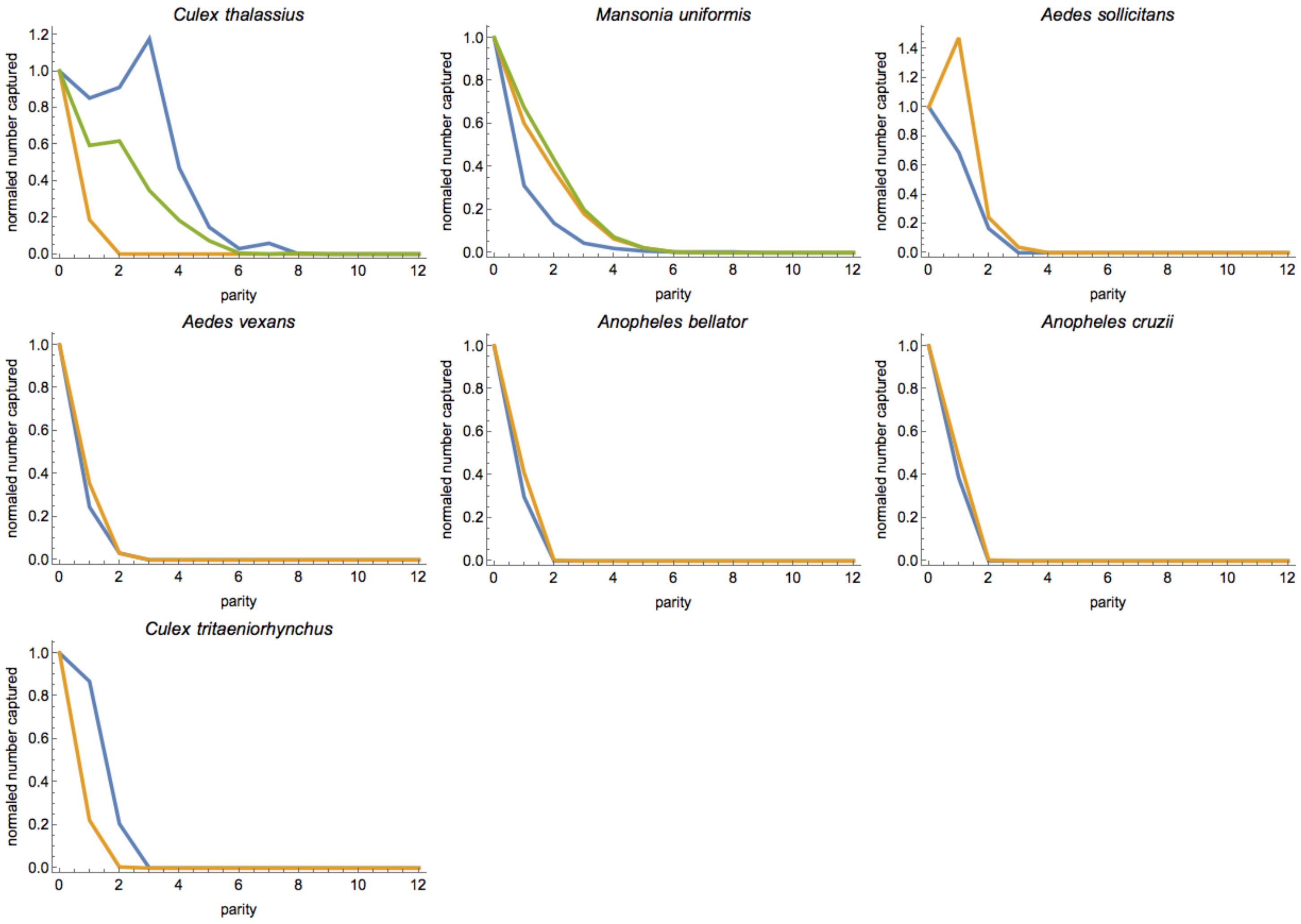


Figure SM15: **Polovodova dissection: normalised physiological age series for nine species in the database.** Each different coloured line represents an individual series. In each case, the count for all ages has been normalised by the nulliparous count. In all cases, we do not include any data for censored observations (see Section SM2.2.1).

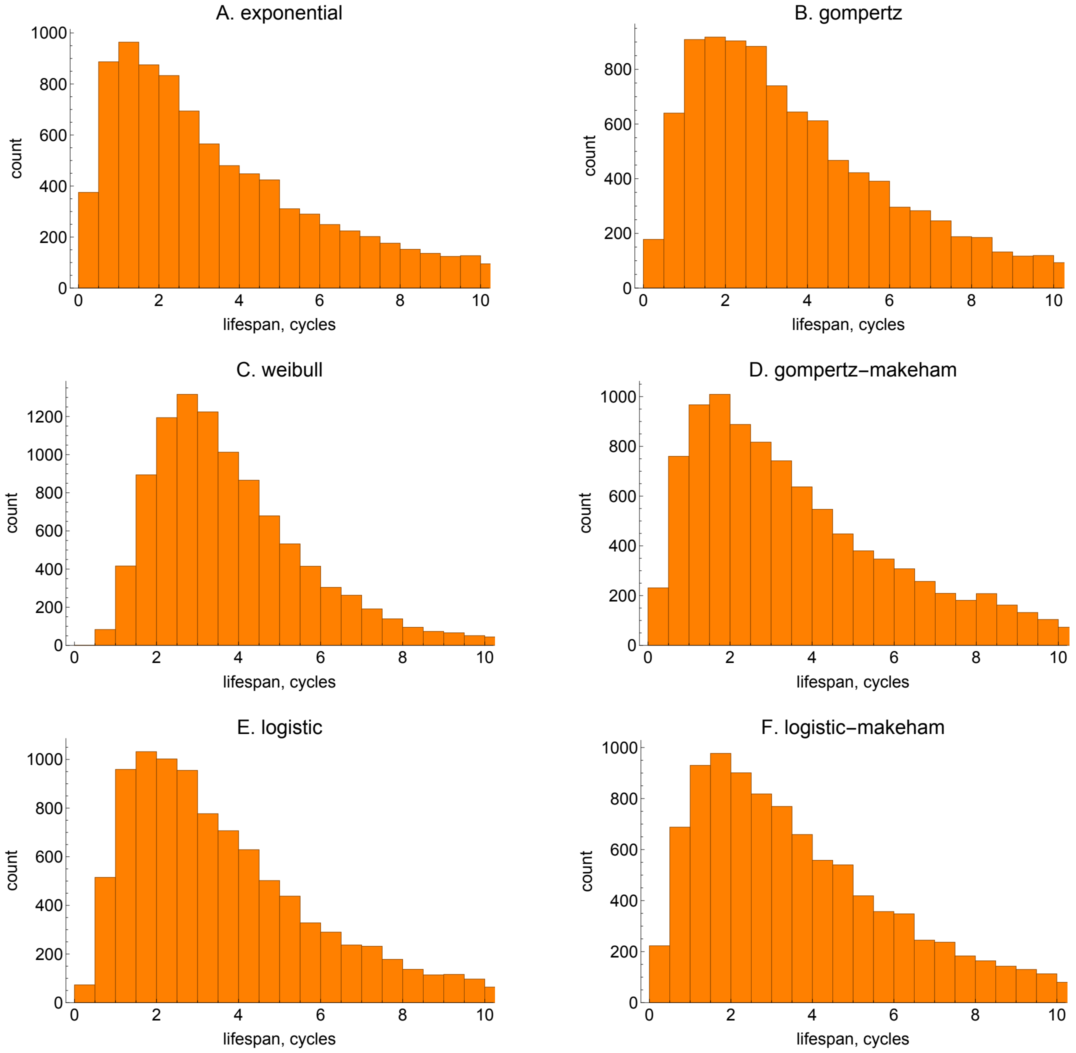


Figure SM16: **Polovodova dissection: prior predictive mean lifespan for each mortality model.** The distributional form of the priors is given in Table SM7. In all cases, the plots show data for 10,000 samples from the relevant prior distribution.

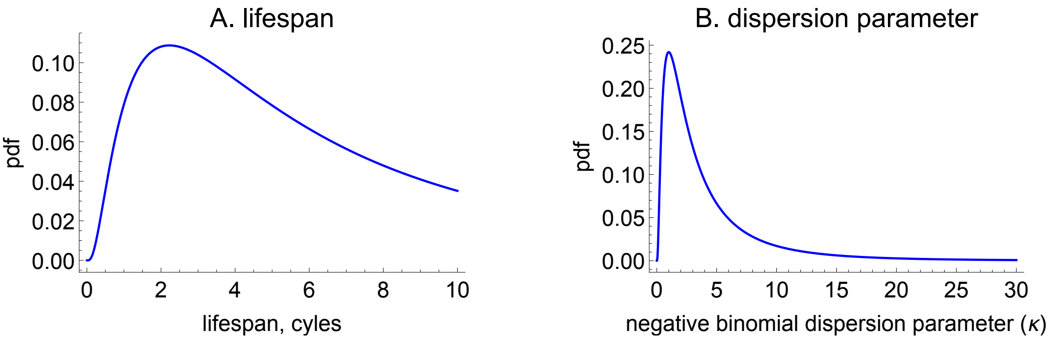


Figure SM17: **Polovodova dissection: priors for A. mean lifespan and B. the over-dispersion parameter used to analyse the individual dissection series.** The prior on the rate parameter of the exponential distribution was $\lambda \sim \text{log-normal}(-1.8, 1)$. The prior on the dispersion parameter was $\kappa \sim \text{log-normal}(1, 1)$.

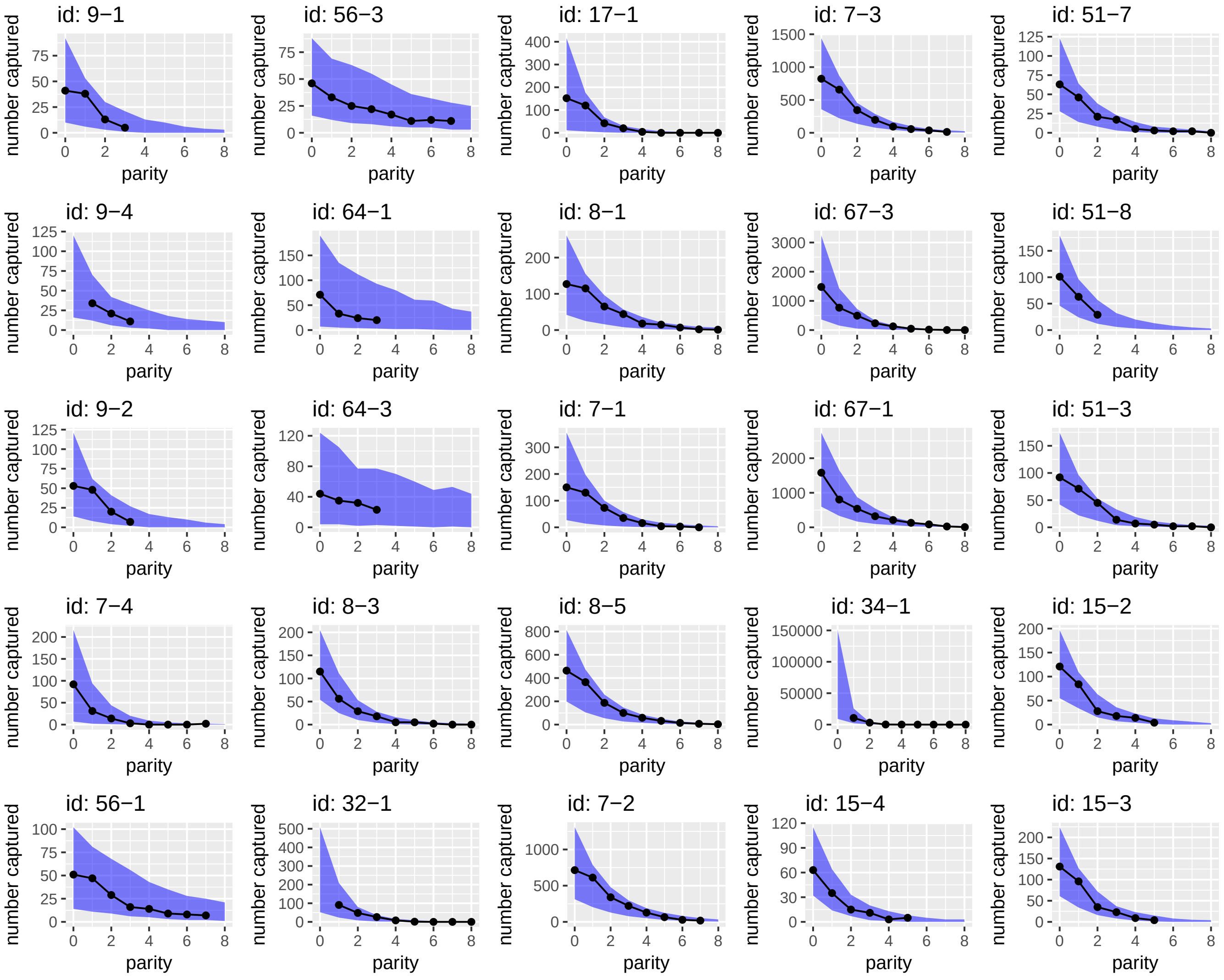


Figure SM18: **Polovodova dissection: posterior predictive checks.** The numbers of mosquitoes recaptured (black lines) versus the 95% central posterior interval of the posterior predictive distribution (blue shading) for a selection of the Polovodova dissection series. The titles correspond to the unique combination of "identifier - physiological-experiment-id" in the database. For the rest of the posterior predictive checks for the dissection data models, see the file referenced in the text.

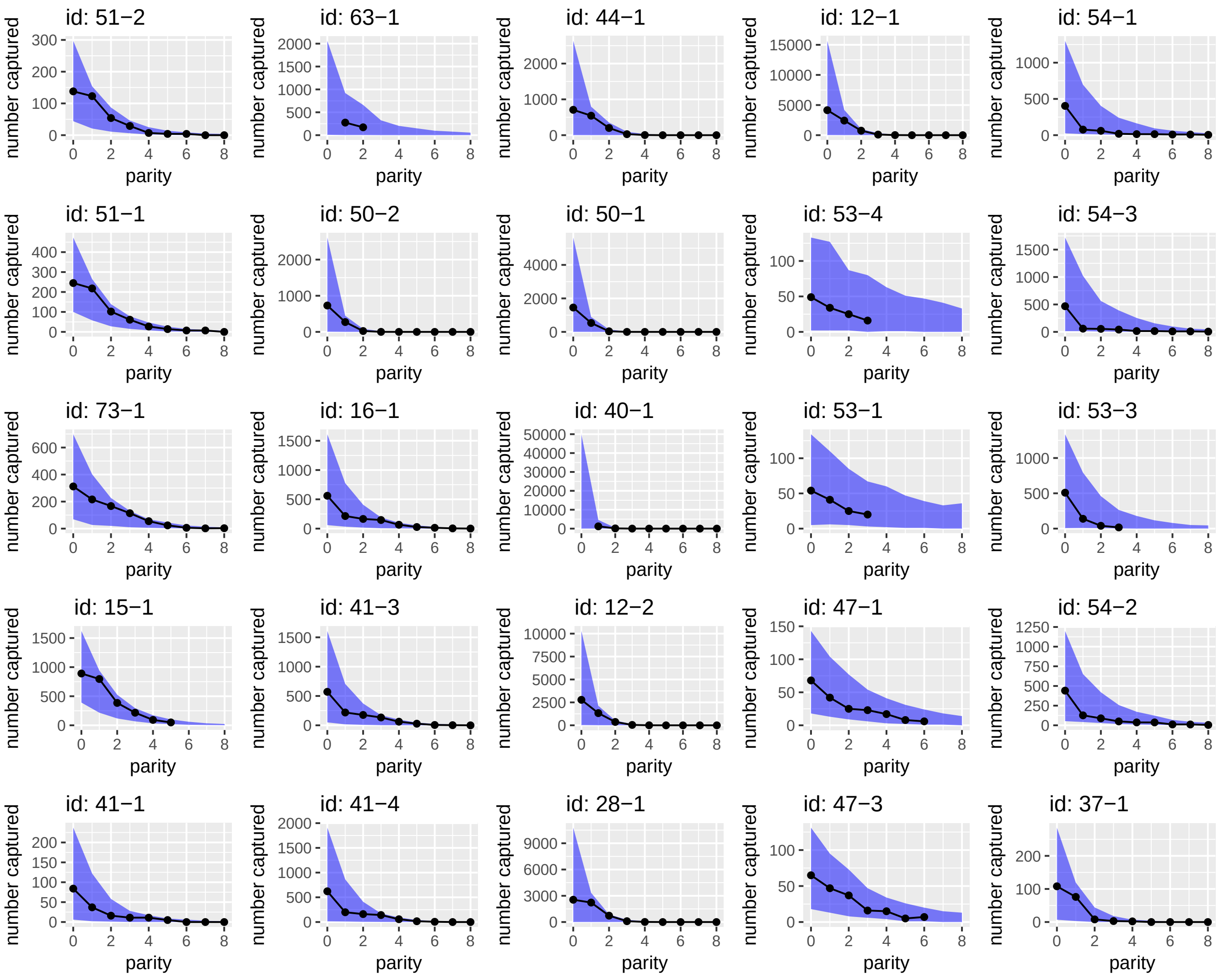


Figure SM19: **Polovodova dissection: posterior predictive checks.** The numbers of mosquitoes recaptured (black lines) versus the 95% central posterior interval of the posterior predictive distribution (blue shading) for a selection of the Polovodova dissection series. The titles correspond to the unique combination of "identifier - physiological-experiment-id" in the database. For the rest of the posterior predictive checks for the dissection data models, see the file referenced in the text.

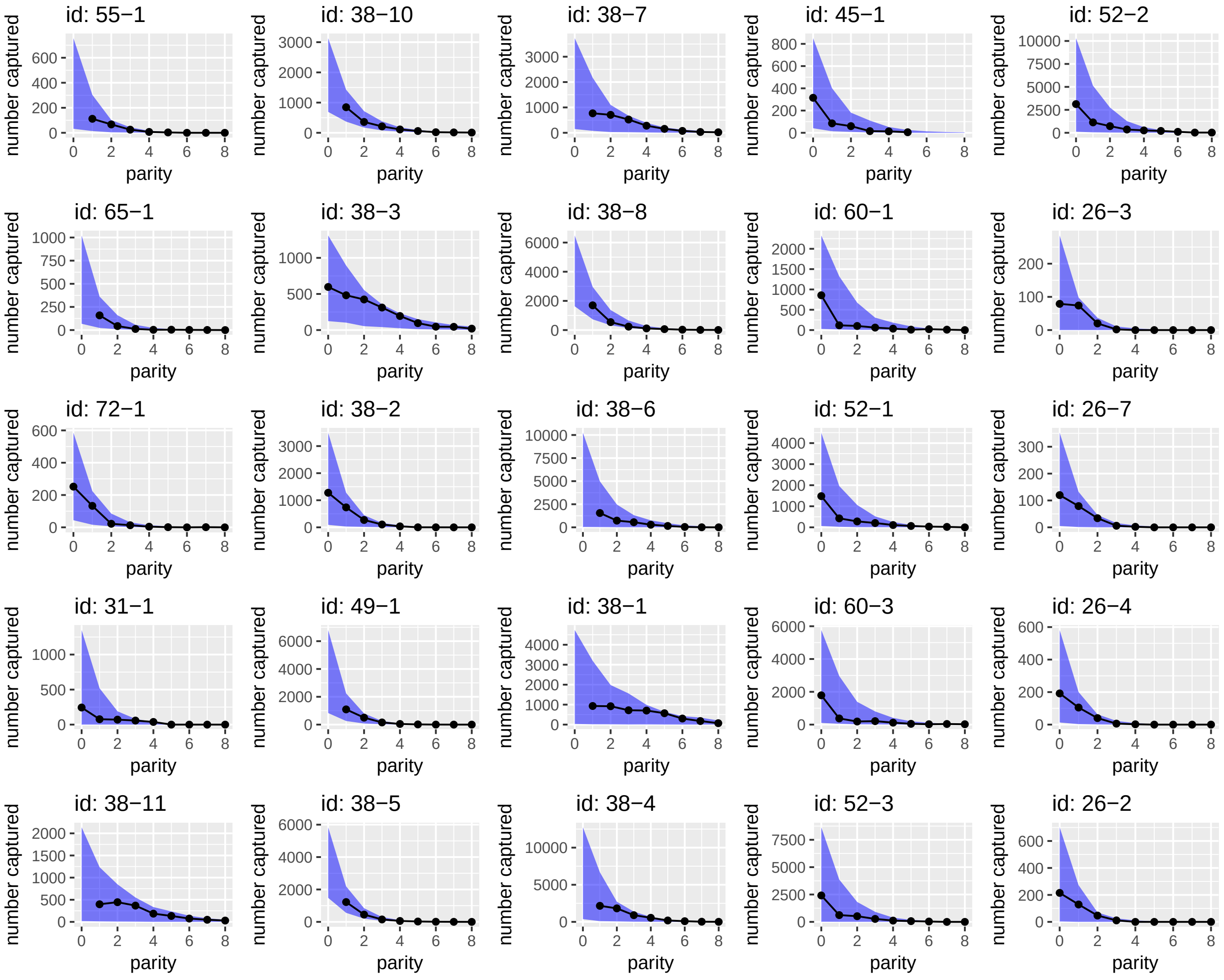


Figure SM20: **Polovodova dissection: posterior predictive checks.** The numbers of mosquitoes recaptured (black lines) versus the 95% central posterior interval of the posterior predictive distribution (blue shading) for a selection of the Polovodova dissection series. The titles correspond to the unique combination of "identifier - physiological-experiment-id" in the database. For the rest of the posterior predictive checks for the dissection data models, see the file referenced in the text.

#### SM2.5 Data collection for gonotrophic cycle duration

To convert the estimates of lifespan in physiological age into chronological age requires a duration for the gonotrophic cycle. To determine this characteristic, we conducted a meta-analysis of previously-published studies that estimate the duration of the gonotrophic cycle. A search of the literature using Google Scholar (`scholar.google.co.uk`) was performed using the search term: 'gonotrophic cycle duration'. The list of articles was then supplemented with a list of references discussed by Silver (2007). Based on the abstracts of the resultant list of published studies, we then decided whether to search each article for estimates of the duration of the gonotrophic cycle. Overall 79 separate estimates of this parameter were found across 42 published articles (see `gonotrophic.csv`). Along with information about the estimates, we also recorded study and series meta-data, including the location of the study, the method used for estimation, species and genus.

Whilst compiling our dataset on gonotrophic cycles, Massey *et al.* (2016) published a database of bionomic quantities for malaria vectors (that is, including only anopheline species). Included in this dataset were a number of estimates of gonotrophic cycle duration. After removing duplicates with our dataset, we were left with 120 estimates of gonotrophic cycle duration. The majority of these estimates were based on laboratory experiments conducted on wild-caught specimens or their $F_1$ or $F_2$ descendents ($n = 45$), followed by MRR studies ($n = 36$), then experiments on laboratory colonies ($n = 11$). In MRR studies, estimates are made of the duration of gonotrophic cycles by dissecting recaptured mosquitoes at each time point using the method of Polovodova (1949) to count ovariolar dilations. For laboratory studies, duration of gonotrophic cycles is determined by direct observation.

Along with point estimates of the parameter we also collected information about the uncertainty in the estimates (if available). In many articles, the duration of the gonotrophic cycle was estimated separately for the first versus subsequent cycles, and these estimates were recorded separately. If there was no disaggregation of durations between 1st and subsequent cycles, we assumed that each of these were the same. Raw estimates of gonotrophic cycle duration were obtained for species across three different genera (*Aedes*, *Anopheles* and *Culex*), although there was a bias towards *Anopheles* with $n = 92$ estimates (Fig. SM21).

#### SM2.6 Statistical analysis of gonotrophic cycle data

As discussed in Section SM2.5, there was considerable study-level heterogeneity in the information provided for the estimates of the gonotrophic cycle duration. A number of studies ($n = 42$) provided no estimates of uncertainty in gonotrophic cycle duration, whereas the rest gave some indication of confidence or alternatively

a range of possible estimates. However, the types of uncertainty intervals that were specified varied considerably across studies: from the vague (but common), '4-6 days'; to the more helpful, '5±1 day (95% confidence interval)'.
The heterogeneous nature of the gonotrophic cycle estimates requires a method that explicitly accounts for this characteristic of the data. We decided to model the estimates as representing quantiles from an underlying normal distribution that represents uncertainty over possible durations of the gonotrophic cycle. In those circumstances where lower, central and upper bounds were given explicitly, the data was fairly symmetric, so we believe assuming the observations come from an unskewed normal distribution is reasonable. Explicitly we treated each type of uncertainty interval as follows,

- Simple range, for example, '4-6 days': treat the lower and upper bounds as the 2.5% and 97.5% quantiles of a normal distribution.
- Confidence intervals, '5±1 day (95% confidence interval)': treat the lower and upper bounds as the relevant quantiles of a normal distribution.
- Point estimates, for example '5 days': treat this as the median of a normal distribution.

The benefit of this approach is that we can convert the quantiles from our normal distribution parameterised by a mean ($\mu$) and standard deviation ($\sigma$) to equivalent quantiles from a standard normal,

$$Z_x = \frac{G_x - \mu}{\sigma}, \tag{21}$$

where $Z_x$ and $G_x$ indicate $x\%$ quantiles from a standard normal distribution and a normal distribution respectively. By manipulating the above expression we obtain the following,

$$G_x = \sigma Z_x + \mu, \tag{22}$$

which forms a straight line in $(Z, G)$ space with slope $\sigma$ and y-intercept $\mu$. Therefore by estimating a linear regression of $Z$ on $G$, we can characterise the underlying $\mathcal{N}(\mu, \sigma)$ distribution from which they are assumed to be drawn.
An ANOVA using the central estimates of 1st gonotrophic cycle duration indicated that there was significant variation in this quantity between genera (ANOVA: $F_{2,116} = 8.7, p < 0.01$; Kruskal-Wallis rank sum test: $\chi^2_2 = 23.9, p < 0.01$), so we estimated separate durations for each genus. The duration of the 1st gonotrophic cycles were estimated as follows: for *Anopheles*, $\mathcal{N}(3.65, 0.40)$; for *Aedes*, $\mathcal{N}(4.62, 0.51)$; and, for *Culex*, $\mathcal{N}(5.23, 0.47)$. For the subsequent durations, the estimated durations were: for *Anopheles*, $\mathcal{N}(3.21, 0.38)$; for *Aedes*,

$\mathcal{N}(4.78, 0.59)$; and, for *Culex*, $\mathcal{N}(5.15, 0.47)$. Pooling data across all genera, the 1st cycle was estimated as $\mathcal{N}(3.95, 0.43)$ and the subsequent as $\mathcal{N}(3.61, 0.42)$. For the *Mansonia* observations on physiological lifespan collected, we converted these using the overall (i.e. across all genera) estimates of gonotrophic cycle durations.

#### SM2.7 Conversion of lifespan from physiological to calendar age

The estimates of lifespan produced from analysing the dissection data are in terms of physiological age (the number of gonotrophic cycles undertaken). To allow comparison with the estimates from the MRR studies and to produce more useful estimates to inform disease transmission dynamical models, we convert these estimates to calendar days. To do this, we use the estimated parameters of the normal densities that we have assumed represent uncertainty in gonotrophic cycle duration (see Section SM2.6) and use them to convert our posterior samples of mean physiological lifespan $(L_1^p, ..., L_S^p)$ into lifespan in calendar ages $(L_1^c, ..., L_S^c)$. To do this, we iterate the following for all $i \in (1, ..., S)$ posterior samples of physiological lifespan for each genus $(j)$,

1. Sample $G_{1i} \sim \mathcal{N}(\mu_{1j}, \sigma_{1i})$, to obtain a duration for the 1st gonotrophic cycle (here, $\mu_{1j}$ and $\sigma_{1j}$ are parameters characterising the mean and std. dev. of a normal representing 1st cycle duration for genus $j$).

2. Sample $G_{2i} \sim \mathcal{N}(\mu_{2j}, \sigma_{2j})$, to obtain a duration for subsequent gonotrophic cycles (here, $\mu_2$ and $\sigma_2$ are parameters characterising the mean and std. dev. of a normal representing subsequent cycle duration for genus $j$).

3. If $L_i^p > 1$, the mean lifespan is longer than one gonotrophic cycle:

    then $L_i^c = G_{1i} + G_{2i} \times (L_i^p - 1)$.

4. Else:

    then $L_i^c = G_{1i} \times L_i^p$.

In using the above methodology to convert from physiological to calendar age, we implicitly assume that all gonotrophic cycles after the first are of the same length (i.e. we choose one $G_{2i}$ per sample). However, since we allow a different $G_{2i}$ for each sample, we nonetheless believe that the above approach allows for sufficient uncertainty in the estimates of subsequent gonotrophic cycle durations. If instead we believed that significant variation in gonotrophic cycle occurred within a particular mosquito's life (as well as between mosquitoes) then we could draw a new value of $G_{2i}$ for each subsequent cycle. This produces results with indistinguishable uncertainty than the results we report (data not shown).

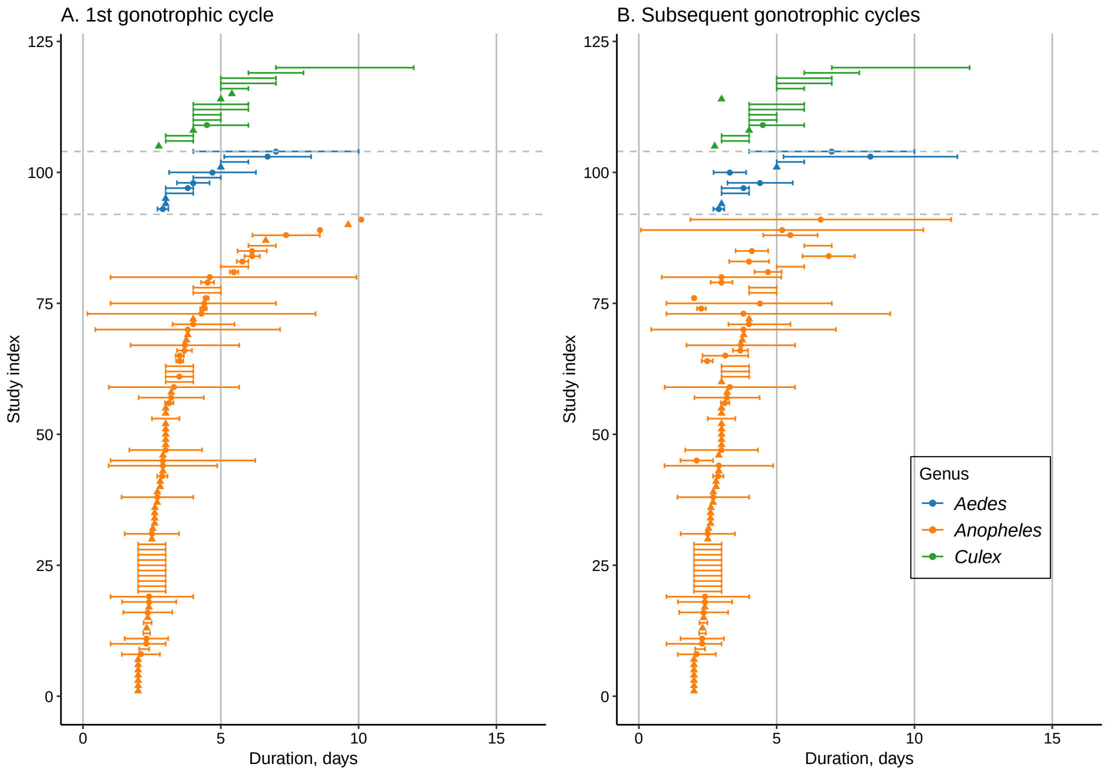


Figure SM21: **Raw gonotrophic cycle duration estimates for A. the 1st cycle and B. subsequent cycles.** When present, the confidence intervals represent our estimates of the 95% confidence interval on gonotrophic cycle duration (see Section SM2.6 for a discussion of how we handle this issue); if our estimate of the lower limit was below zero, we deemed this unbiological and set it to 1 day for this graph. If no statement of uncertainty was given in the estimates then we indicate this by a triangle marker. The studies are ordered within each Genus according to the central estimate of gonotrophic duration.

### SM3 Detinova dissection experiments

Detinova *et al.* (1962) provided an alternative approach to estimate the lifespan of individual specimens, based on sampling the parous rate in a population (the proportion of adult females that have laid eggs) using dissection to determine parity of each specimen. In what follows, we follow the derivation given by Davidson (1954), to determine an expression for the mean lifespan of a mosquito in terms of gonotrophic cycles (all assumed to be of the same duration, $G$). If the probability of dying in a given cycle, $p$, is assumed constant (i.e. there is no age-dependent mortality), then the probability a mosquito completes $Y$ cycles, is geometrically distributed,

$$Pr(Y = k) = (1 - p)^k p, \tag{23}$$

where $k = 0, 1, 2, ....$ This implies that the proportion of adults surviving to become parous is $Pr(Y \geq 1) = 1 - Pr(Y = 0) = 1 - p = \mathcal{P}$. Therefore, the proportion of parous mosquitoes in any given sample $i$ of wild-caught mosquitoes, $\mathcal{P}^{\{i\}}$ is a direct estimator of the probability of surviving a given cycle, $1 - p$. The mean lifespan of a mosquito in terms of completed gonotrophic cycles is then given by $(1 - p)/p = \mathcal{P}/(1 - \mathcal{P})$.

If, instead, we choose chronological time as our measure of lifespan, the above analysis still holds, although with $q$ now representing the probability of dying on a particular day, meaning $(1 - q)^G$ is the probability of surviving a cycle. The daily survival probability $1 - q$ is, hence, equal to $\sqrt[G]{\mathcal{P}}$, and the mean lifespan is given by $q/(1 - q) = \sqrt[G]{\mathcal{P}}/(1 - \sqrt[G]{\mathcal{P}})$.

#### SM3.1 Data processing

In this section, we describe the data processing carried out on the Massey *et al.* (2016) dataset, which provides a global database of bionomic quantities for the dominant vector species of malaria. Included in this database is the number of parous and nulliparous samples in studies where Detinova's dissection approach was applied to field caught specimens.

For each of the three datasets (Africa, Americas and Asia) contained with the SOM of Massey *et al.* (2016), we used only those observations which had a non-missing and positive total capture size. For each observation, we calculated observed parity (= parous count/total capture size) and compared this quantity with the 'parity percent' column in the datasets. If the difference between these two quantities was lower than 1%, we considered the discrepancy to be due to numerical rounding and did not investigate further. In those cases where the difference exceeded 1%, we did the following:

- Africa: considering the database entry for Ijumba *et al.* (2002), there was a difference of about 50%. Reading the paper, it looks like there is a mistake in their Table 5, and that the parous number should be 131 and, as such, we changed this number in the database; considering the entry for Service *et al.* (1995), there was a difference of around 5%. We could not access the paper here so assumed that the issue was with the 'parity percent' entry.

- Americas: no differences exceeding 1%.
- Asia: considering the entry for Kitthawee *et al.* (1992), there was a difference of around 3%. Reading the paper, it looks like there was a (small) error in the database, and the parous number should be 35 (rather than 36) – this adjustment was made for the ensuing analysis.

For the species- and complex-level estimates of lifespan, we considered only those groups with five or more parity experiments. For the continent-level and overall estimates, we used all data. In a number of cases, a species was indicated as a complex (for example, 'Anopheles gambiae complex'), presumably when insufficient information was available in the paper to identify the species. These data were dropped for the species-level analysis but were included for all other analyses. The Massey *et al.* (2016) database included a variable for each observation determining whether insecticide-based controls were in place at the specified location and time period of the study. For $n = 883$ observations, this information was recorded, allowing us to estimate the impact of these interventions on lifespan (see SM3.5). This analysis determined that insecticidal interventions reduced lifespan, and, thus, studies where insecticides were in place ($n = 364$) were removed from our main analysis.
For the species-level analyses, there were, hence, $n = 198$ observations for Africa across 4 species; $n = 107$ observations for the Americas across 6 species; and $n = 108$ observations for Asia across 4 species. For the complex-level analyses, there were, hence, $n = 679$ observations for Africa across 4 groups; $n = 121$ observations for the Americas across 5 groups; and $n = 318$ observations for Asia across 13 groups. For the continent-level and overall analyses, there were $n = 1126$ observations across Africa ($n = 679$), the Americas ($n = 126$) and Asia ($n = 321$).

#### SM3.2 Models

We estimated lifespan at the species and complex / subgroup level (hereafter denoted as *s.l.*) using a Bayesian hierarchical framework. We denote this grouping variable by $s$ in what follows. The number of parous individuals $y_i^{\{s\}}$ in sample $i$ (consisting of individuals belonging to group $s$) was modelled by a binomial sampling distribution,

$$y_i^{\{s\}} \sim \mathcal{B}(N_i, \mathcal{P}_i^{\{s\}}), \tag{24}$$

where $N_i$ is the sample size and $\mathcal{P}_i^{\{s\}}$ is the proportion parous in sample $i$ for group $s$. A hierarchical prior structure was used for the parous proportion in each sample,

$$\mathcal{P}_i^{\{s\}} \sim \text{beta}\left(\phi^{\{s\}}\kappa^{\{s\}}, (1-\phi^{\{s\}})\kappa^{\{s\}}\right), \tag{25}$$

where $0 \leq \phi^{\{s\}} \leq 1$ is the mean parous proportion and $\kappa^{\{s\}} > 1$ is a concentration parameter (both of which are group-specific). The priors we used in this model were: $\phi \sim \text{beta}(20, 15)$ and $\kappa \sim \text{Pareto}(5, 1.5)$. These were chosen because this yielded a prior predictive distribution with a mean lifespan close to 10 (see Fig. SM22).

The Stan code used to fit the hierarchical model to data is shown below.

```
data{
  int N; // total obs
  int X[N]; // number parous
  int num[N]; // number of samples collected
  int species[N]; // integer index array with group identity
  int K; // number of different groups
}

parameters{
  vector<lower=0, upper=1>[N] theta;
  vector<lower=0, upper=1>[K] phi;
  vector<lower=5>[K] kappa;
}

model{
  for(i in 1:N){
    X[i] ~ binomial(num[i], theta[i]);
    theta[i] ~ beta(phi[species[i]] * kappa[species[i]],
                    (1 - phi[species[i]]) * kappa[species[i]]);
  }
  kappa ~ pareto(5, 1.5);
  phi ~ beta(20, 15);
}

generated quantities{
  real lifespan[K];
  real survival[K];
  real theta_average[K];
  for(i in 1:K){
    real gon = normal_rng(3.65, 0.4);
    theta_average[i] = beta_rng(phi[i] * kappa[i],

    survival[i] = theta_average[i] ^ (1.0 / gon);
    lifespan[i] = survival[i] / (1.0 - survival[i]);
  }
}
```

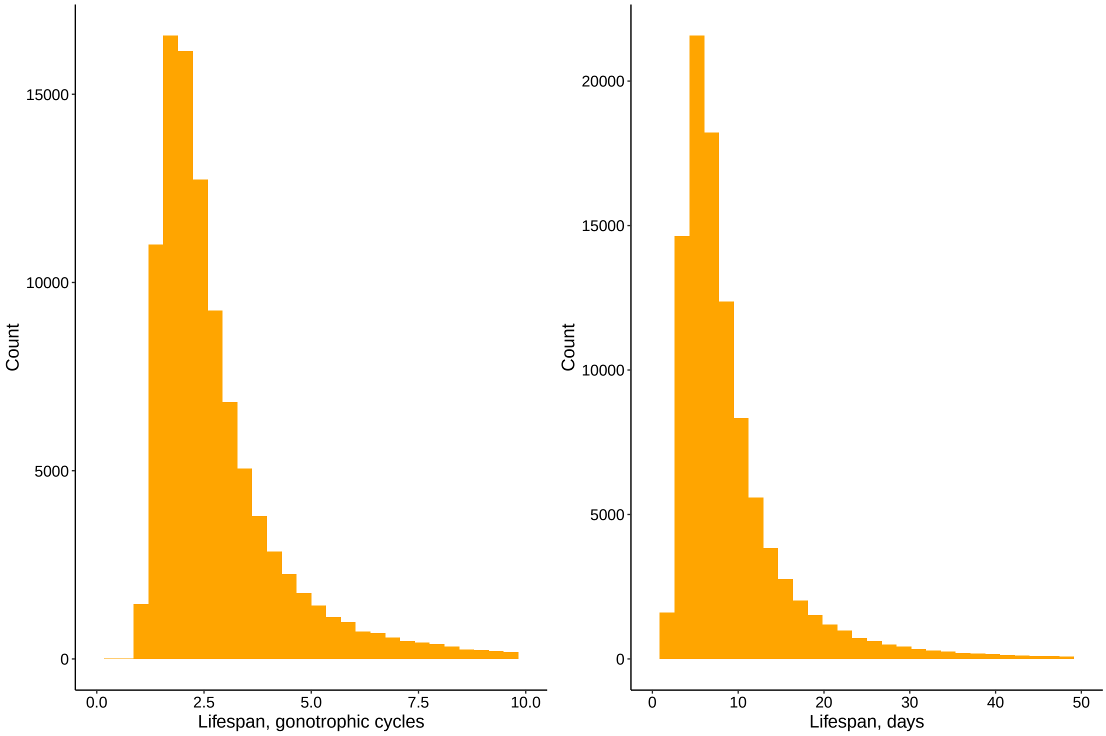


Figure SM22: **Detinova dissection: prior predictive distribution of lifespan in terms of (left) physiological age and (right) chronological age.** Here, we used the gonotrophic cycle duration estimated for *Anopheles*, $G \sim \mathcal{N}(3.65, 0.40)$, to convert physiological survival to calendar days. The figure shows 100,000 samples from the prior predictive distribution. The mean lifespan in terms of gonotrophic cycles is approximately 3; in terms of chronological age, it is approximately 10.

#### SM3.3 Model estimation by MCMC and cross validation

The statistical models were fit to data using Stan's NUTS algorithm Carpenter *et al.* (2016). In all cases, models were run for 4000 iterations on each of 12 chains, with the first half of the iterations discarded as warm-up. We monitored diagnosed convergence as per Gelman & Rubin (1992), with $\hat{R} < 1.1$ on all parameters. In all model runs, there were fewer than a handful of divergent transitions.
To evaluate model fit to data, we used K-fold cross validation with 4 folds for each analysis. This means that, in a run, approximately 3/4 of data was used to train models, and the remaining data used to evaluate out-of-sample predictive capability through the log-likelihood. To evaluate out-of-sample predictive log-likelihood for a given data point, we drew data-level parameters $\mathcal{P}_i^{\{s\}}$ from the relevant top-level distribution; for example, for a species-level fit, we drew data-

level parameters from the species-level distribution relevant to the species for the data point in question.
The folds were chosen so that, at all levels of analysis, there was always at least a handful of data points belonging to a particular group in the training set. This meant that, to generate meaningful comparisons between the species-level, complex-level and continent-level models, we selected only those data points with at least 5 observations for each species and complex. This reduced the number of data points used for model comparison versus those used for model estimation. For Africa, we had $n = 217$ data points; for the Americas, there were $n = 107$; and for Asia, there were $n = 108$.
For the comparison between continent-level grouping and overall, we were able to use more data points since we were less constrained by the minimal numbers of data belonging to each group. This meant we had $n = 1126$ data points, with $n = 679$ for Africa, $n = 126$ for the Americas, and $n = 321$, for Asia.

#### SM3.4 Model checking

To check the fit of models to the data, we simulated from the posterior predictive distribution for each grouping used in analysis. This means that we have two sets of posterior predictive figures: those where the check was performed at the species level; and those corresponding to the complex or subgroup level. In each of these figures, we plot observed empirical cumulative distribution function (eCDF) for the proportion parous versus the equivalent from each posterior predictive dataset.

- **Africa**: the complex-level posterior predictive distribution graph is shown in Figure SM23; the species-level graph in Figure SM24.
- **Americas**: the complex-level posterior predictive distribution graph is shown in Figure SM25; the species-level graph in Figure SM26.
- **Asia**: the complex-level posterior predictive distribution graph is shown in Figure SM27; the species-level graph in Figure SM28.

In all cases, the eCDF for the actual data was bounded by the range of the posterior predictive eCDFs. By this metric, we conclude that the model seems to represent the data well.

#### SM3.5 Association of insecticide interventions on lifespan

To determine the association of insecticidal interventions with lifespan, we modified the model described in §SM3.2 to estimate effects at the species level. To do so, we allowed a logit-linear impact of insecticide. Further, the species-level effects

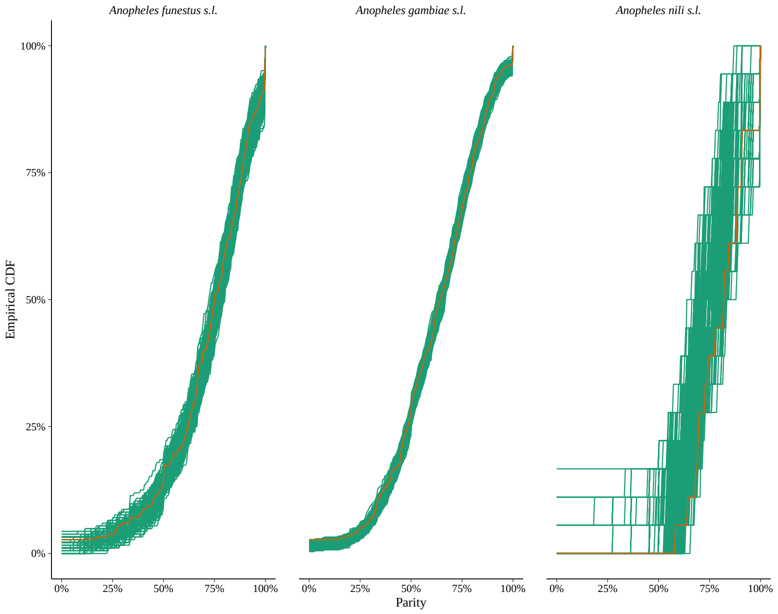


Figure SM23: **Detinova dissection: posterior predictive check for Africa, at the complex-level.** Plots show the empirical cumulative distribution functions for observations (orange) and posterior predictive simulations (green). In each case, we ran 4 Markov chains, with 200 iterations per chain; discarding the first 100 iterations as warm-up.

were modelled hierarchically using normal priors, allowing the impact of resistance to vary at the observation level within each study. The model to estimate this is shown below (where the variable metering the impact of insecticide at the observational level is 'delta'). This model was estimated using 1000 iterations per chain (500 of which were warm-up) across 12 chains.

```
data{
  int N;
  int X[N];
  int num[N];
  int species[N];
  int K;
  real z[N];
}

parameters{
  vector<lower=0, upper=1>[N] theta;
  vector<lower=0, upper=1>[K] phi;
  vector<lower=5>[K] kappa;
  vector[K] delta_raw;
  real delta_top;
  real<lower=0> delta_sigma;
}

transformed parameters{
  vector[K] delta;
  for(i in 1:K)
    delta[i] = delta_top + delta_sigma * delta_raw[i];
}

model{
  real theta_eff;
  for(i in 1:N){
    theta_eff = logit(theta[i]) + delta[species[i]] * z[i];
    X[i] ~ binomial_logit(num[i], theta_eff);
    theta[i] ~ beta(phi[species[i]] * kappa[species[i]],
                    (1 - phi[species[i]]) * kappa[species[i]]);
  }
  kappa ~ pareto(5, 1.5);
  phi ~ beta(20, 15);
  delta_raw ~ normal(0, 1);
  delta_top ~ normal(0, 1);
  delta_sigma ~ normal(0, 1);
}
```

```
generated quantities{
  real lifespan_without_insecticide[K];
  real lifespan_with_insecticide[K];
  for(i in 1:K){
    real theta_eff;
    real theta_eff_without;
    real gon = normal_rng(3.65, 0.4);
    real survival;
    real survival_without;
    theta_eff = logit(theta[i]) + delta[species[i]];
    theta_eff_without = logit(theta[i]);
    survival = inv_logit(theta_eff)^(1.0/gon);
    survival_without = inv_logit(theta_eff_without)^(1.0/gon);
    lifespan_with_insecticide[i] = survival / (1.0 - survival);
    lifespan_without_insecticide[i] = survival_without /
                                     (1.0 - survival_without);
  }
}
```

#### SM3.6 Association of weather covariates with temperature

To investigate the effect of measures of temperature, we extracted daily data from the ERA-interim re-analysis by ECMWF (Dee *et al.* , 2011) through their website using the Python API. This data included:

- Two-meter surface temperature of air at 2m above the surface of land, sea or in-land waters.
- Daily minimum two-meter surface temperature.
- Daily maximum two-meter surface temperature.

For this analysis, we included only those studies where the start month listed in the database coincided with the end date, meaning the study took place within a given month. We also omitted those data points where insecticide use was recorded. To ensure that we had sufficient data points for each fold used for model comparison, we considered only those species with over 100 observations. This resulted in $n = 366$ observations across 2 species complexes: *A. gambiae s.l.* ($n = 257$) and *A. funestus s.l.* ($n = 109$).
The temperature data was extracted for the GPS position of each study ($\pm 0.1$ degrees) for each day of the month over which mosquito sampling took place,

with a grid size of 0.05 degrees. The Python script to extract this information is given below.

```
from ecmwfapi import ECMWFDataServer
import time
def extract_meanminmax_temperature(a_area, a_date_start,
a_date_end):
server = ECMWFDataServer()
date_range = str(a_date_start) + "/to/" + str(a_date_end)
date_file = str(a_date_start) + "_to_" + str(a_date_end)
area=(str(a_area["north"]) + '/' + str(a_area["west"]) +
'/' + str(a_area["south"]) + '/' + str(a_area["east"]))
area_file = (str(a_area["north"]) + '_' + str(a_area["west"])
+ '_' + str(a_area["south"]) + '_' + str(a_area["east"]))
a_filename = ("../data/raw/weather/temperature_" +
str(area_file) + "_" + date_file + ".nc")
server.retrieve({
"class": "ei",
"dataset": "interim",
"date": str(date_range),
"expver": "1",
"grid": "0.05/0.05",
"levtype": "sfc",
"area" : str(area),
"param": "167.128/201.128/202.128",
"step": "3",
"stream": "oper",
"time": "00:00:00",
"type": "fc",
"target":  str(a_filename),
"format" : "netcdf",
})
```

To determine the impact of temperature on mosquito lifespan, we estimated a generalised linear model using a logit link, with hierarchical complex-level intercepts and either no temperature term, and linear temperature terms with or without quadratic temperature terms. In each case, these temperature terms were allowed to vary at the complex level (non-hierarchically). This meant our likelihood had the form,

$$X_i^{\{s\}} \sim \mathcal{B}(N_i, \text{inv-logit}(\alpha_i^{\{s\}})), \tag{26}$$

where,

$$\alpha_i^{\{s\}} = \text{logit}(\mathcal{P}_i^{\{s\}}) + \delta_1^{\{s\}} T_i + \delta_2^{\{s\}} T_i^2. \tag{27}$$

Here $T_i$ is a measure of temperature at study location $i$ (measures described below); $\mathcal{P}_i$ was set a prior according to eq. (25) and $\delta_1^{\{s\}} \sim \mathcal{N}(0,1)$ and $\delta_2^{\{s\}} \sim \mathcal{N}(0,1)$. The temperature measures ($T_i$), we investigated were summary statistics calculated across the daily data for all days within the month each study took place and across all grid points falling within the area centred on the GPS location of each study (described above),

- Average two meter temperature (average of mean daily data for each grid point),
- Daily range in two meter temperature (average of maximum minus minimum daily temperature for each grid point). This likely reflects the average day-night temperature range.

Each of these two models were estimated in Stan using 1000 iterations per chain (500 of each were warm-up) across 4 chains and thinned samples by a factor of 20. To determine whether each covariate was predictive for lifespan, we undertook K-fold cross-validation (using 4 folds with a given fold holding roughly equal numbers of each species).

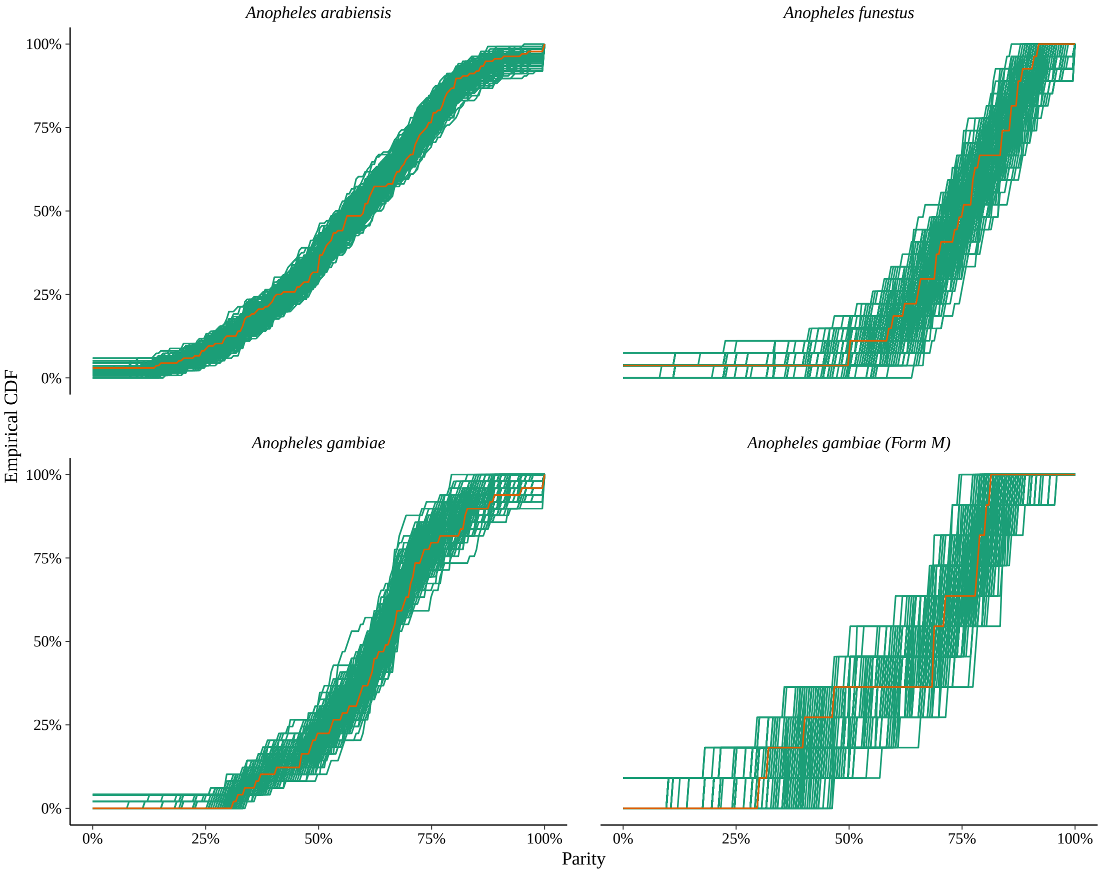


Figure SM24: **Detinova dissection: posterior predictive check for Africa, at the species-level.** Plots show the empirical cumulative distribution functions for observations (orange) and posterior predictive simulations (green). In each case, we ran 4 Markov chains, with 200 iterations per chain; discarding the first 100 iterations as warm-up.

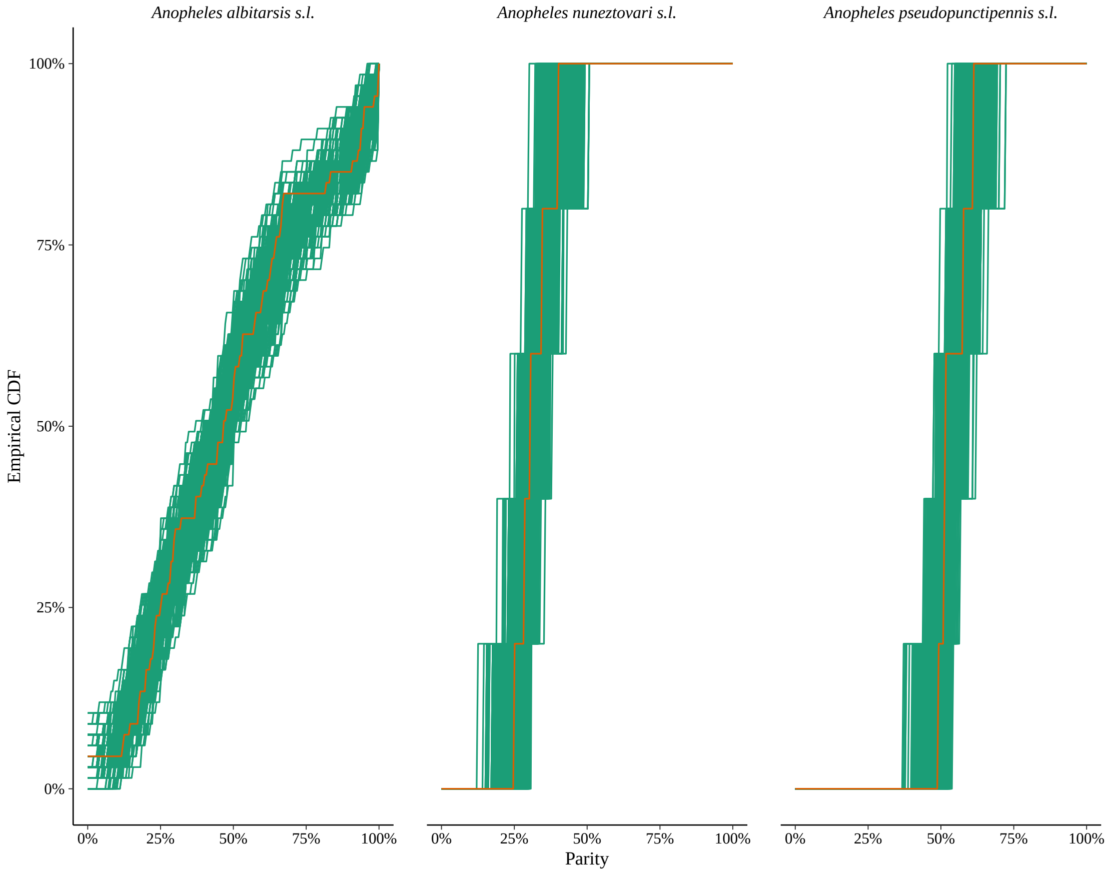


Figure SM25: **Detinova dissection: posterior predictive check for the Americas, at the complex-level.** Plots show the empirical cumulative distribution functions for observations (orange) and posterior predictive simulations (green). In each case, we ran 4 Markov chains, with 200 iterations per chain; discarding the first 100 iterations as warm-up.

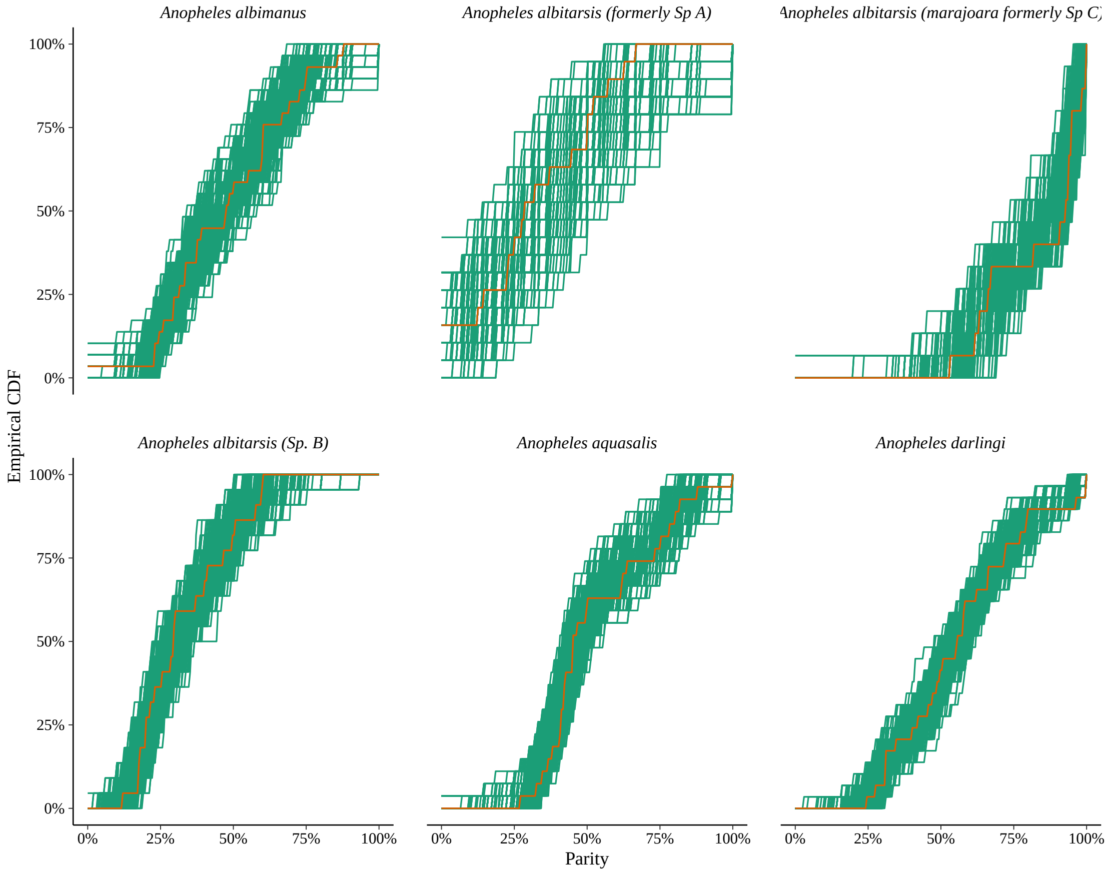


Figure SM26: **Detinova dissection: posterior predictive check for the Americas, at the species-level.** Plots show the empirical cumulative distribution functions for observations (orange) and posterior predictive simulations (green). In each case, we ran 4 Markov chains, with 200 iterations per chain; discarding the first 100 iterations as warm-up.

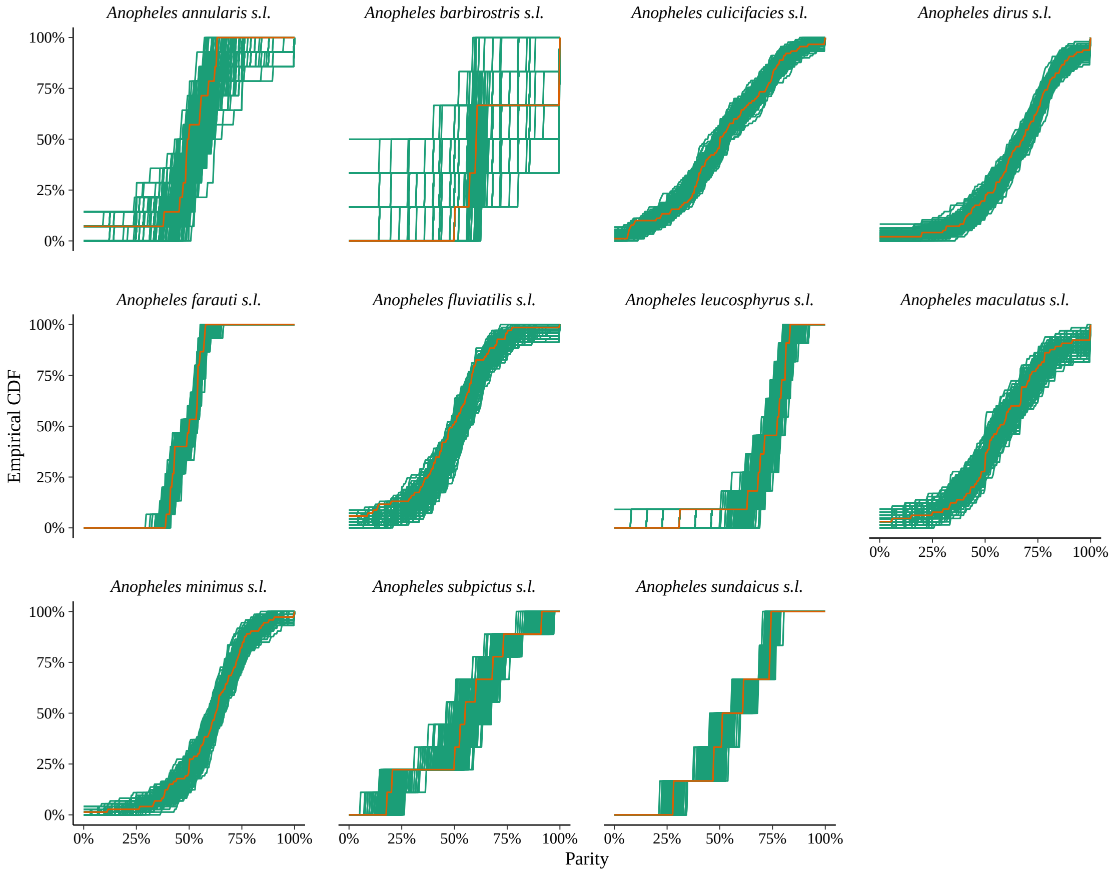


Figure SM27: **Detinova dissection: posterior predictive check for Asia, at the complex-level.** Plots show the empirical cumulative distribution functions for observations (orange) and posterior predictive simulations (green). In each case, we ran 4 Markov chains, with 200 iterations per chain; discarding the first 100 iterations as warm-up.

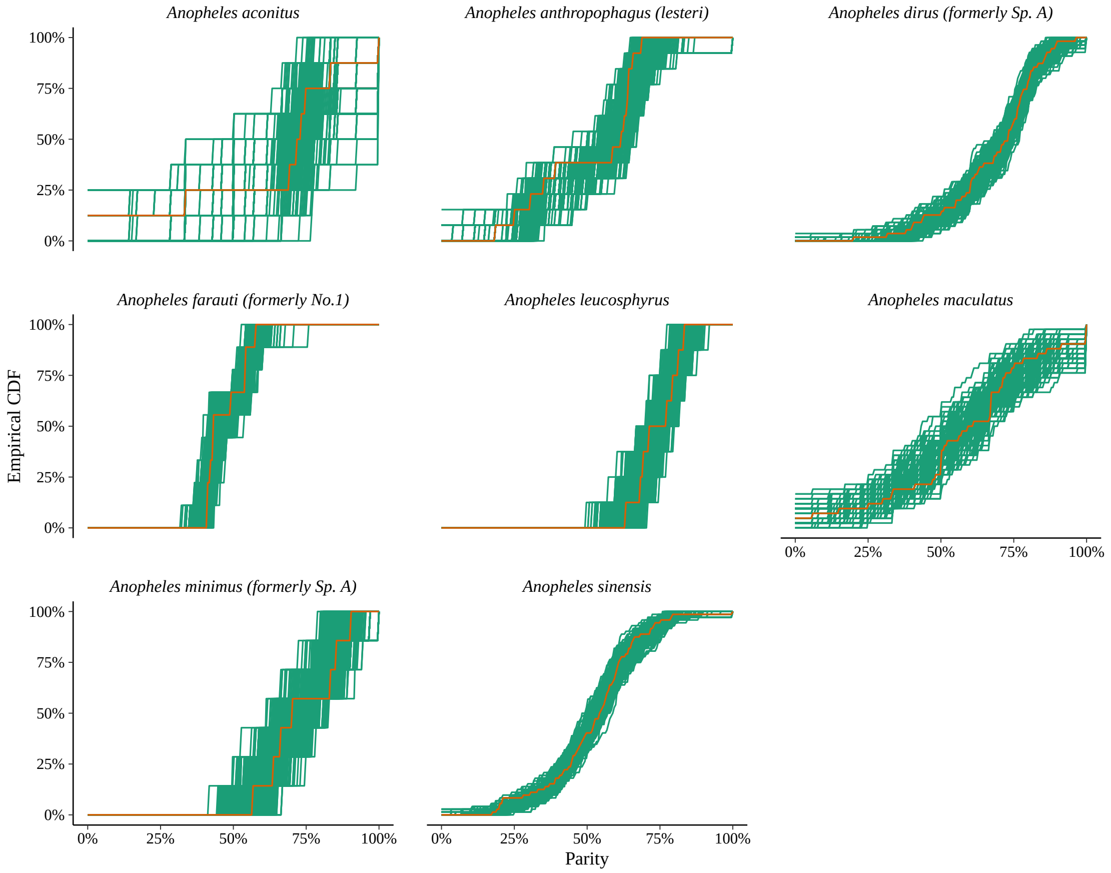


Figure SM28: **Detinova dissection: posterior predictive check for Asia, at the species-level.** Plots show the empirical cumulative distribution functions for observations (orange) and posterior predictive simulations (green). In each case, we ran 4 Markov chains, with 200 iterations per chain; discarding the first 100 iterations as warm-up.

### SM4 EIPs of vector species of malaria, dengue fever, chikungunya and Zika

The following anopheline species were identified as vector species of malaria,

- **Africa**: *A. gambiae s.l.*, *A. funestus* and *A. melas* (Sinka *et al.* , 2012), and *A. rivulorum* (Wilkes *et al.* , 1996).
- **Americas**: *A. albimanus*, *A. albitarsis*, *A. darlingi*, *A. pseudopunctipennis* and *A. quadrimaculatus* (Sinka *et al.* , 2012); *A. bellator* (Forattini *et al.* , 1999; Lorenz *et al.* , 2012); *A. cruzii* (Lorenz *et al.* , 2012); and *A. vestitipennis* (Sinka *et al.* , 2010).
- **Europe and the Middle-East**: *A. sergentii* (Sinka *et al.* , 2012); and *A. maculipennis* (Hackett *et al.* , 1935).
- **Asia**: *A. farauti s.l.*, *A. koliensis*, *A. lesteri*, *A. maculatus*, *A. minimus*, *A. punctulatus*, *A. stephensi* and *A. subpictus s.l.* (Sinka *et al.* , 2012); and *A. culicifacies* (Green & Miles, 1980).

For dengue fever, chikungunya and Zika, the main vector species are *Ae. aegypti* and *Ae. albopictus* (Kraemer *et al.* , 2015; Grard *et al.* , 2014; Benelli & Mehlhorn, 2016).

### SM5 Studies included in the MRR meta-analysis

The following is the subset of studies from the original Guerra *et al.* (2014) database which were used in the MRR meta-analysis: Marini *et al.* (2010); Baber *et al.* (2010); Lacroix *et al.* (2009); Maciel-de Freitas *et al.* (2008); Midega *et al.* (2007); Maciel-De-Freitas *et al.* (2007); Elizondo-Quiroga *et al.* (2006); Ba *et al.* (2005); Fabian *et al.* (2005); La Corte Dos Santos *et al.* (2004); Watson *et al.* (2000); Harrington *et al.* (2001); Tsuda *et al.* (2001); Muir & Kay (1998); Touré *et al.* (1998); Quinones *et al.* (1997); Costantini *et al.* (1996); Trpis *et al.* (1995); Jensen & Washino (1994); Fernandez-Salas *et al.* (1994); Jaal & MacDonald (1992); Rodriguez *et al.* (1992); Chiang *et al.* (1991); Jensen & Washino (1991); Eldridge & Reeves (1990); Macdonald *et al.* (1968); Pumpuni & Walker (1989); Charlwood & Bryan (1987); Charlwood & Alecrim (1989); Birley & Charlwood (1989); Arredondo-Jiménez *et al.* (1998); Hii *et al.* (1990); Renshaw *et al.* (1994); Milby & Reisen (1989); Maciel-de Freitas *et al.* (2007); Loong *et al.* (1990); Nelson & Milby (1980); Maciel-de Freitas *et al.* (2006); McDonald (1977); Curtis & Rawlings (1980); Rawlings & Davidson (1982); Conway *et al.* (1974);

Reisen *et al.* (1979); Nelson *et al.* (1978); Rawlings *et al.* (1981); Sempala (1981); Takagi *et al.* (1995); Buei *et al.* (1980); Eyles *et al.* (1943a); Ordóñez González *et al.* (2001); Pant & Yasuno (1973); Charlwood *et al.* (1988); Reisen *et al.* (1982); Nayar *et al.* (1980); Carnevale *et al.* (1979); Eyles *et al.* (1946); Reisen *et al.* (1984); Charlwood *et al.* (1986); Trpis & Hausermann (1975); Lutwama & Mukwaya (1994); Wada *et al.* (1969); Takken *et al.* (1998); Abdel-Malek *et al.* (1966); Valerio *et al.* (2012); Zetek (1915); Takagi *et al.* (1995); Yasuno *et al.* (1975); Eyles *et al.* (1943b); Germain *et al.* (1974).

### SM6 Studies included in the dissection study meta-analysis

The following is a list of studies included in the dissection study meta-analysis: Catangui *et al.* (1971); Charlwood & Wilkes (1979); Chang *et al.* (1991); Charlwood *et al.* (2000); Edalat *et al.* (2015); de Barros *et al.* (2011); Lines *et al.* (1991a); Magesa *et al.* (1991); Lines *et al.* (1991b); Forattini *et al.* (1996); Samarawickrema (1967, 1968); de Barros *et al.* (2007); Charlwood *et al.* (1985a); Chandra *et al.* (1996); Hoc & Wilkes (1995); Vythilingam *et al.* (1997); Russell (1986); Chadee & Ritchie (2010); Ebsary & Crans (1977); Charlwood (1980); Shriram & Krishnamoorthy (2011); Mahmood & Reisen (1981); Samarawickrema *et al.* (1987); Uttah *et al.* (2013); Ramaiah & Das (1992); Buei & Ito (1982); Chadee *et al.* (1995); Chandra (2008); Foll & Pant (1966); Kanda *et al.* (1975); Mala *et al.* (2014); Shalaby *et al.* (1962); World Health Organisation *et al.* (1960); Reisen *et al.* (1980); Detinova *et al.* (1962); Smith & Kurtz (1994); Jayanetti *et al.* (1987); Ch *et al.* (2013); Pant *et al.* (1962); Ghosh *et al.* (2010); Mahanta *et al.* (1999); Schlein & Müller (2012); Penilla *et al.* (2002); Schlein & Müller (2015); Snow & Boreham (1978); Detinova (1968); Nathan (1981); Charlwood *et al.* (1985b); Schlein & Müller (2008); Beier *et al.* (2012); Müller & Schlein (2005); Surendran *et al.* (2006); Müller *et al.* (2010); Mendis *et al.* (1998); Wilkes *et al.* (1996); Chatterjee & Chandra (2000); Müller *et al.* (2010); Tuchinda *et al.* (1969); Chan *et al.* (1971); Aigbodion *et al.* (2011); Qualls *et al.* (2015); Reisen *et al.* (1981); Gillies & Wilkes (1972, 1965); Kay (1979); Hitchcock Jr (1968).

### SM7 Studies included in the gonotrophic cycle duration meta-analysis

The following is a list of studies included in the gonotrophic study meta-analysis: Gillies & Wilkes (1965); Sheppard *et al.* (1969); de Meillon *et al.* (1967); Pant &

Yasuno (1973); Germain *et al.* (1974); Lowe *et al.* (1973); Fernandez-Salas *et al.* (1994); Buei *et al.* (1980); Rawlings & Curtis (1982); Mori *et al.* (1977); Sempala (1981); Reisen *et al.* (1983); Birley & Charlwood (1989); Suzuki (1978); Charlwood & Bryan (1987); Charlwood *et al.* (1986); Charlwood & Wilkes (1979); Charlwood *et al.* (1995); Chang *et al.* (1991); Edalat *et al.* (2015); de Barros *et al.* (2011); Samarawickrema (1968, 1967); Charlwood *et al.* (1985a); Chandra *et al.* (1996); Russell (1986); Chadee & Ritchie (2010); Mahmood & Reisen (1981); Ahumada *et al.* (2004); Kenawy (1991); Rajagopalan (1980); Chandra (2008); Scholl *et al.* (1979); Mala *et al.* (2014); Afrane *et al.* (2005); Gillies (1953); Quinones *et al.* (1997); Rúa *et al.* (2005); Delatte *et al.* (2009); Arredondo-Jiménez *et al.* (1998); Wong *et al.* (2014); Ijumba *et al.* (2002).

### References


Abdel-Malek, Albert A, Abdel-Aal, MA, *et al.* . 1966. Study of the dispersion and flight range of *Anopheles sergentii* Theo in Siwa Oasis using radioactive isotopes as markers. *Bulletin of the World Health Organisation*, **35**, 968–973.

Afrane, Yaw A, Lawson, Bernard W, Githeko, Andrew K, & Yan, Guiyun. 2005. Effects of microclimatic changes caused by land use and land cover on duration of gonotrophic cycles of *Anopheles gambiae* (Diptera: Culicidae) in western Kenya highlands. *Journal of Medical Entomology*, **42**(6), 974–980.

Ahumada, Jorge A, Lapointe, Dennis, & Samuel, Michael D. 2004. Modeling the population dynamics of *Culex quinquefasciatus* (Diptera: Culicidae), along an elevational gradient in Hawaii. *Journal of Medical Entomology*, **41**(6), 1157–1170.

Aigbodion, FI, Uyi, OO, Akintelu, OH, & Salau, LA. 2011. Pelagia Research Library. *European Journal of Experimental Biology*, **1**(4), 173–180.

Anderson, Roy M, & May, Robert M. 1992. *Infectious diseases of humans: dynamics and control.* Vol. 28. Wiley Online Library.

Arredondo-Jiménez, Juan I, Rodríguez, Mario H, & Washino, Robert K. 1998. Gonotrophic cycle and survivorship of *Anopheles vestitipennis* (Diptera: Culicidae) in two different ecological areas of southern Mexico. *Journal of Medical Entomology*, **35**(6), 937–942.

Ba, Yamar, Diallo, Diawo, Kebe, Cheikh Mouhamed Fadel, Dia, Ibrahima, & Diallo, Mawlouth. 2005. Aspects of bioecology of two Rift Valley fever virus vectors in Senegal (West Africa): *Aedes vexans* and *Culex poicilipes* (Diptera: Culicidae). *Journal of Medical Entomology*, **42**(5), 739–750.

Baber, Ibrahima, Keita, Moussa, Sogoba, Nafomon, Konate, Mamadou, Doumbia, Seydou, Traore, Sekou F, Ribeiro, Jose MC, Manoukis, Nicholas C, *et al.* . 2010. Population size and migration of *Anopheles gambiae* in the Bancoumana Region of Mali and their significance for efficient vector control. *PLoS One*, **5**(4), e10270.

Beier, John C, Müller, Günter C, Gu, Weidong, Arheart, Kristopher L, & Schlein, Yosef. 2012. Attractive toxic sugar bait (ATSB) methods decimate populations of *Anopheles* malaria vectors in arid environments regardless of the local availability of favoured sugar-source blossoms. *Malaria Journal*, **11**(1), 31.

Bellan, Steve E. 2010. The importance of age dependent mortality and the extrinsic incubation period in models of mosquito-borne disease transmission and control. *PLoS One*, **5**(4), e10165.

Benelli, Giovanni, & Mehlhorn, Heinz. 2016. Declining malaria, rising of dengue and Zika virus: insights for mosquito vector control. *Parasitology research*, **115**(5), 1747–1754.

Birley, MH, & Charlwood, JD. 1989. The effect of moonlight and other factors on the oviposition cycle of malaria vectors in Madang, Papua New Guinea. *Annals of Tropical Medicine & Parasitology*, **83**(4), 415–422.

Bryan, JH, Foley, DH, Geary, M, & Carven, CTJ. 1991. Anopheles annulipes Walker (Diptera: Culicidae) at Griffith, New South Wales. 3. Dispersal of two sibling species. *Australian Journal of Entomology*, **30**(2), 119–121.

Buei, K, Ito, S, Nakamura, H, & Yoshida, M. 1980. Field studies on the gonotrophic cycle of *Culex tritaeniorhynchus*. *Medical Entomology and Zoology*, **31**(1), 57–62.

Buei, Kazuo, & Ito, Sumiyo. 1982. The age-composition of field populations and the survival rates in *Culex tritaeniorhynchus Giles*. *Medical Entomology and Zoology*, **33**(1), 21–25.

Carey, James R. 2001. Insect biodemography. *Annual Review of Entomology*, **46**(1), 79–110.

Carnevale, Pierre, Bosseno, Marie-France, Molinier, Michel, Lancien, Jeannick, Le Pont, François, & Zoulani, Albert. 1979. Etude du cycle gonotrophique d?*Anopheles gambiae* (Diptera, Culicidae)(Giles, 1902) en zone de forêt degradée d?Afrique Centrale. *Cahiers Orstom. Série Entomologie Médicale et Parasitologie*, **17**, 55–75.

Carpenter, Bob, Gelman, Andrew, Hoffman, Matt, Lee, Daniel, Goodrich, Ben, Betancourt, Michael, Brubaker, Michael A, Guo, Jiqiang, Li, Peter, & Riddell,

Allen. 2016. Stan: A probabilistic programming language. *Journal of Statistical Software.*

Catangui, FP, *et al.* . 1971. Studies on the gonotrophic cycle of *Anopheles minimus flavirostris* and the application of physiological age grading technique on the same species. *Southeast Asian Journal of Tropical Medicine and Public Health*, **2**(3), 384–92.

Ch, Goutam, Paramanik, Manas, Mondal, Samir Kumar, Ghosh, Arup Kumar, *et al.* . 2013. Comparative Studies of Different Indices Related to Filarial Vector of a Rural and an Urban Area of West Bengal, India. *Tropical Medicine & Surgery.*

Chadee, Dave D, & Ritchie, Scott A. 2010. Oviposition behaviour and parity rates of *Aedes aegypti* collected in sticky traps in Trinidad, West Indies. *Acta Tropica*, **116**(3), 212–216.

Chadee, Dave D, Ganesh, R, Hingwan, JO, & Tikasingh, Elisha S. 1995. Seasonal abundance, biting cycle and parity of the mosquito Haemagogus leucocelaenus in Trinidad, west indies. *Medical and Veterinary Entomology*, **9**(4), 372–376.

Chan, Kai-Lok, *et al.* . 1971. Life table studies of *Aedes albopictus* (Skuse). *Sterility principles for insect control or eradication*, 131–144.

Chandra, G, Seal, B, & Hati, AK. 1996. Age composition of the filarial vector *Culex quinquefasciatus* (Diptera: Culicidae) in Calcutta, India. *Bulletin of Entomological Research*, **86**(3), 223–226.

Chandra, Goutam. 2008. Age composition of incriminated malaria vector in a rural foothills in West Bengal, India. *Indian Journal of Medical Research*, **127**(6), 607.

Chang, Moh-Seng, Chan, Kai-Lok, Ho, Beng-Chuan, & Hawley, William A. 1991. Comparative transmission potential of three *Mansonia spp.* (Diptera: Culicidae) for filariasis in Sarawak, Malaysia. *Bulletin of Entomological Research*, **81**(4), 437–444.

Charlwood, Jacques D, & Alecrim, WA. 1989. Capture-recapture studies with the South American malaria vector *Anopheles darlingi*, Root. *Annals of Tropical Medicine & Parasitology*, **83**(6), 569–576.

Charlwood, JD. 1980. Observations on the bionomics of *Anopheles darlingi* Root (Diptera: Culicidae) from Brazil. *Bulletin of Entomological Research*, **70**(4), 685–692.

Charlwood, JD, & Bryan, JH. 1987. A mark?recapture experiment with the filariasis vector *Anopheles punctulatus* in Papua New Guinea. *Annals of Tropical Medicine & Parasitology*, **81**(4), 429–436.

Charlwood, JD, & Wilkes, TJ. 1979. Studies on the age-composition of samples of *Anopheles darlingi* Root (Diptera: Culicidae) in Brazil. *Bulletin of Entomological Research*, **69**(02), 337–342.

Charlwood, JD, Birley, MH, Dagoro, H, Paru, R, & Holmes, PR. 1985a. Assessing survival rates of *Anopheles farauti* (diptera: Culicidae) from Papua New Guinea. *The Journal of Animal Ecology*, 1003–1016.

Charlwood, JD, Dagoro, H, & Paru, R. 1985b. Blood-feeding and resting behaviour in the *Anopheles punctulatus Dönitz* complex (Diptera: Culicidae) from coastal Papua New Guinea. *Bulletin of Entomological Research*, **75**(3), 463–476.

Charlwood, JD, Graves, PM, & Birley, MH. 1986. Capture-recapture studies with mosquitoes of the group of *Anopheles punctulatus* Dönitz (Diptera: Culicidae) from Papua New Guinea. *Bulletin of Entomological Research*, **76**(2), 211–227.

Charlwood, JD, Graves, PM, & Marshall, TF de C. 1988. Evidence for a memorized home range in *Anopheles farauti* females from Papua New Guinea. *Medical and Veterinary Entomology*, **2**(2), 101–108.

Charlwood, JD, Kihonda, J, Sama, S, Billingsley, PF, Hadji, H, Verhave, JP, Lyimo, E, Luttikhuizen, PC, & Smith, T. 1995. The rise and fall of *Anopheles arabiensis* (Diptera: Culicidae) in a Tanzanian village. *Bulletin of Entomological Research*, **85**(1), 37–44.

Charlwood, JD, Vij, R, & Billingsley, PF. 2000. Dry season refugia of malaria-transmitting mosquitoes in a dry savannah zone of east Africa. *The American Journal of Tropical Medicine and Hygiene*, **62**(6), 726–732.

Chatterjee, Soumendranath, & Chandra, Goutam. 2000. Role of *Anopheles subpictus* as a primary vector of malaria in an area in India. *Japanese Journal of Tropical Medicine and Hygiene*, **28**(3), 177–181.

Chiang, GL, Loong, KP, Chan, ST, Eng, KL, & Yap, HH. 1991. Capture-recapture studies with *Anopheles maculatus Theobald* (Diptera: Culicidae) the vector of malaria in peninsular Malaysia. *Southeast Asian Journal of Tropical Medicine and Public Health*, **22**, 643–647.

Clements, AN, & Paterson, GD. 1981. The analysis of mortality and survival rates in wild populations of mosquitoes. *Journal of Applied Ecology*, 373–399.

Conway, GR, Trpis, M, & McClelland, GAH. 1974. Population parameters of the mosquito *Aedes aegypti* (L.) estimated by mark-release-recapture in a suburban habitat in Tanzania. *The Journal of Animal Ecology*, 289–304.

Costantini, Carlo, LI, Song-Gang, Torre, Alessandra Della, Sagnon, N'Fale, Coluzzi, Mario, & Taylor, Charles E. 1996. Density, survival and dispersal of *Anopheles gambiae* complex mosquitoes in a West African Sudan savanna village. *Medical and Veterinary Entomology*, **10**(3), 203–219.

Curtis, CF, & Rawlings, P. 1980. A preliminary study of dispersal and survival of *Anopheles culicifacies* in relation to the possibility of inhibiting the spread of insecticide resistance. *Ecological Entomology*, **5**(1), 11–17.

Davidson, G. 1954. Estimation of the survival-rate of anopheline mosquitoes in nature. *Nature.*

Dawes, Emma J, Churcher, Thomas S, Zhuang, Shijie, Sinden, Robert E, & Basáñez, María-Gloria. 2009. *Anopheles* mortality is both age-and *Plasmodium*-density dependent: implications for malaria transmission. *Malaria Journal*, **8**(1), 228.

de Barros, Fábio Saito Monteiro, Arruda, Mércia Eliane, Vasconcelos, Simão D, Luitgards-Moura, José Francisco, Confalonieri, Ulisses, Rosa-Freitas, Maria Goreti, Tsouris, Pantelis, Lima-Camara, Tamara Nunes, & Honório, Nildimar Alves. 2007. Parity and age composition for *Anopheles darlingi* Root (Diptera: Culicidae) and *Anopheles albitarsis Lynch-Arribálzaga* (Diptera: Culicidae) of the northern Amazon Basin, Brazil. *Journal of Vector Ecology*, **32**(1), 54–68.

de Barros, Fabio Saito Monteiro, Honorio, Nildimar Alves, & Arruda, Mercia Eliane. 2011. Survivorship of *Anopheles darlingi* (Diptera: Culicidae) in relation with malaria incidence in the Brazilian Amazon. *PLoS One*, **6**(8), e22388.

de Meillon, Botha, Sebastian, Anthony, & Khan, ZH. 1967. Time of arrival of gravid *Culex pipiens fatigans* at an oviposition site, the oviposition cycle and the relationship between time of feeding and time of oviposition. *Bulletin of the World Health Organisation*, **36**(1), 39.

Dee, DP, Uppala, SM, Simmons, AJ, Berrisford, P, Poli, P, Kobayashi, S, Andrae, U, Balmaseda, MA, Balsamo, G, Bauer, d P, *et al.* . 2011. The ERA-Interim reanalysis: Configuration and performance of the data assimilation system. *Quarterly Journal of the royal meteorological society*, **137**(656), 553–597.

Delatte, Hélène, Gimonneau, Geoffrey, Triboire, Aurélie, & Fontenille, Didier. 2009. Influence of temperature on immature development, survival, longevity, fecundity, and gonotrophic cycles of *Aedes albopictus*, vector of chikungunya and dengue in the Indian Ocean. *Journal of Medical Entomology*, **46**(1), 33–41.

Detinova, Tatjana Sergeevna, *et al.* . 1962. Age grouping methods in Diptera of medical importance with special reference to some vectors of malaria. *Monograph series World Health Organisation.*

Detinova, TS. 1968. Age structure of insect populations of medical importance. *Annual Review of Entomology*, **13**(1), 427–450.

Ebsary, BA, & Crans, WJ. 1977. The biting activity of *Aedes sollicitans* in New Jersey. *Mosquito News.*

Edalat, Hamideh, Moosa-Kazemi, Seyed Hassan, Abolghasemi, Esmail, & Khairandish, Sedigheh. 2015. Vectorial capacity and age determination of *Anopheles stephens Liston* (Diptera: Culicidae), during the malaria transmission in Southern Iran. *Journal of Entomology and Zoology Studies*, **3**(1), 256–263.

Eldridge, BF, & Reeves, WC. 1990. Daily survivorship of adult Aedes communis in a high mountain environment in California. *Journal of the American Mosquito Control Association*, **6**(4), 662–666.

Elizondo-Quiroga, Armando, Flores-Suarez, Adriana, Elizondo-Quiroga, Darwin, Ponce-Garcia, Gustavo, Blitvich, Bradley J, Contreras-Cordero, Juan Francisco, Gonzalez-Rojas, Jose Ignacio, Mercado-Hernandez, Roberto, Beaty, Barry J, & Fernandez-Salas, Ildefonso. 2006. Gonotrophic cycle and survivorship of *Culex quinquefasciatus* (Diptera: Culicidae) using sticky ovitraps in Monterrey, northeastern Mexico. *Journal of the American Mosquito Control Association*, **22**(1), 10–14.

Eyles, DE, Bishop, LK, *et al.* . 1943a. An Experiment on the Range of Dispersion of *Anopheles quadrimaculatus. American Journal of Hygiene*, **37**(3).

Eyles, Don E, Cox, Wm W, *et al.* . 1943b. The Measurement of a Population of *Anopheles quadrimaculatus Say. Journal of the National Malaria Society*, **2**(2).

Eyles, Don E, Sabrosky, Curtis W, Russell, John C, *et al.* . 1946. *Long-range dispersal of* Anopheles quadrimaculatus. US Government Printing Office.

Fabian, Mashauri M, Toma, Takako, Tsuzuki, Ataru, Saita, Susumu, & Miyagi, Ichiro. 2005. Mark-release-recapture experiments with *Anopheles saperoi* (Diptera: Culicidae) in the Yona Forest, northern Okinawa, Japan. *Southeast Asian Journal of Tropical Medicine and Public Health*, **36**(1), 54.

Fernandez-Salas, Ildefonso, Rodriguez, Mario Henry, & Roberts, Donald R. 1994. Gonotrophic cycle and survivorship of *Anopheles pseudopunctipennis* (Diptera: Culicidae) in the Tapachula foothills of southern Mexico. *Journal of Medical Entomology*, **31**(3), 340–347.

Foll, CV, & Pant, CP. 1966. The conditions of malaria transmission in Katsina Province, Northern Nigeria, and a discussion of the effects of dichlorvos application. *Bulletin of the World Health Organisation*, **34**(3), 395.

Forattini, Oswaldo Paulo, Kakitani, Iná, Massad, Eduardo, & Marucci, Daniel. 1996. Studies on mosquitoes (Diptera: Culicidae) and anthropic environment: 11-Biting activity and blood-seeking parity of *Anopheles* (Kerteszia) in South-Eastern Brazil. *Revista de Saude Publica*, **30**, 107–114.

Forattini, Oswaldo Paulo, Kakitani, Iná, Santos, Roseli La Corte dos, Ueno, Helene Mariko, & Kobayashi, Keilla Miki. 1999. Role of *Anopheles (Kerteszia) bellator* as malaria vector in southeastern Brazil (Diptera: Culicidae). *Memórias do Instituto Oswaldo Cruz*, **94**(6), 715–718.

Gelman, Andrew, & Rubin, Donald B. 1992. Inference from iterative simulation using multiple sequences. *Statistical Science*, 457–472.

Gelman, Andrew, Carlin, John B, Stern, Hal S, & Rubin, Donald B. 2014. *Bayesian data analysis.* Vol. 2. Taylor & Francis.

Germain, Max, Hervé, Jean-Pierre, & Geoffroy, Bernard. 1974. Evaluation de la durée du cycle trophogonique d'*Aedes africanus* (Théobald), vecteur potentiel de fièvre jaune, dans une galerie forestière du sud de la République Centrafricaine. *Cahiers ORSTOM. Série Entomologie Médicale et Parasitologie*, **12**(2), 127–133.

Ghosh, Anupam, Mandal, Samir, & Chandra, Goutam. 2010. Seasonal distribution, parity, resting, host-seeking behavior and association of malarial parasites of *Anopheles stephensi Liston* in Kolkata, West Bengal. *Entomological Research*, **40**(1), 46–54.

Gillies, MT. 1953. The duration of the gonotrophic cycle in *Anopheles gambiae* and *Anopheles funestus*, with a note on the efficiency of hand catching. *East African Medical Journal*, **30**(4).

Gillies, MT, & Wilkes, TJ. 1965. A study of the age-composition of populations of *Anopheles gambiae* Giles and A. funestus Giles in North-Eastern Tanzania. *Bulletin of Entomological Research*, **56**(02), 237–262.

Gillies, MT, & Wilkes, TJ. 1972. The range of attraction of animal baits and carbon dioxide for mosquitoes. Studies in a freshwater area of West Africa. *Bulletin of Entomological Research*, **61**(3), 389–404.

Grard, Gilda, Caron, Mélanie, Mombo, Illich Manfred, Nkoghe, Dieudonné, Ondo, Statiana Mboui, Jiolle, Davy, Fontenille, Didier, Paupy, Christophe, & Leroy, Eric Maurice. 2014. Zika virus in Gabon (Central Africa)–2007: a new threat from Aedes albopictus? *PLoS neglected tropical diseases*, **8**(2), e2681.

Green, CA, & Miles, SJ. 1980. Chromosomal evidence for sibling species of the malaria vector Anopheles (Cellia) culicifacies Giles. *The Journal of tropical medicine and hygiene*, **83**(2), 75–78.

Guerra, Carlos A, Reiner, Robert C, Perkins, AT, Lindsay, Steve W, Midega, J, Brady, Oliver J, Barker, Christopher M, Reisen, William K, Harrington, Laura C, Takken, Willem, *et al.* . 2014. A global assembly of adult female mosquito mark-release-recapture data to inform the control of mosquito-borne pathogens. *Parasite & Vectors*, **7**(1), 276.

Hackett, Lewis W, Missiroli, Alberto, *et al.* . 1935. The varieties of Anopheles maculipennis and their relation to the distribution of malaria in Europe. *Rivista di Malariologia*, **14**(1).

Hancock, PA, Thomas, MB, & Godfray, HCJ. 2009. An age-structured model to evaluate the potential of novel malaria-control interventions: a case study of fungal biopesticide sprays. *Proceedings of the Royal Society of London B: Biological Sciences*, **276**(1654), 71–80.

Harrington, Laura C, Buonaccorsi, John P, Edman, John D, Costero, Adriana, Kittayapong, Pattamaporn, Clark, Gary G, & Scott, Thomas W. 2001. Analysis of survival of young and old *Aedes aegypti* (Diptera: Culicidae) from Puerto Rico and Thailand. *Journal of Medical Entomology*, **38**(4), 537–547.

Harrington, Laura C, Jones, James J, Kitthawee, Sangvorn, Sithiprasasna, Ratana, Edman, John D, Scott, Thomas W, *et al.* . 2008. Age-dependent survival of the dengue vector *Aedes aegypti* (Diptera: Culicidae) demonstrated by simultaneous release–recapture of different age cohorts. *Journal of Medical Entomology*, **45**(2), 307–313.

Hii, JK, Birley, MH, & Sang, VY. 1990. Estimation of survival rate and oviposition interval of *Anopheles balabacensis* mosquitoes from mark-recapture experiments in Sabah, Malaysia. *Medical and Veterinary Entomology*, **4**(2), 135–140.

Hitchcock Jr, James G. 1968. Age composition of a natural population of *Anopheles quadrimaculatus Say* (Diptera: Culicidae) in Maryland, USA. *Journal of Medical Entomology*, **5**(1), 125–134.

Hoc, TQ, & Wilkes, TJ. 1995. The ovariole structure of *Anopheles gambiae* (Diptera: Culicidae) and its use in determining physiological age. *Bulletin of Entomological Research*, **85**(01), 59–69.

Hoffman, Matthew D, & Gelman, Andrew. 2014. The No-U-turn sampler: adaptively setting path lengths in Hamiltonian Monte Carlo. *Journal of Machine Learning Research*, **15**(1), 1593–1623.

Ijumba, JN, Mosha, FW, & Lindsay, SW. 2002. Malaria transmission risk variations derived from different agricultural practices in an irrigated area of northern Tanzania. *Medical and Veterinary Entomology*, **16**(1), 28–38.

Jaal, Z, & MacDonald, WW. 1992. A mark-release-recapture experiment with *Anopheles lesteri paraliae* in northwest Peninsular Malaysia. *Annals of Tropical Medicine & Parasitology*, **86**(4), 419–424.

Jayanetti, SR, Wijesundera, Manel de S, & Amaresinghe, FP. 1987. A Study on the Bionomics of Indoor Resting Mosquitoes in Kandy.

Jensen, Truls, & Washino, Robert K. 1991. An assessment of the biological capacity of a Sacramento Valley population of Aedes melanimon to vector arboviruses. *The American Journal of Tropical Medicine and Hygiene*, **44**(4), 355–363.

Jensen, Truls, & Washino, Robert K. 1994. Comparison of recapture patterns of marked and released Aedes vexans and Ae. melanimon (Diptera: Culicidae) in the Sacramento Valley of California. *Journal of Medical Entomology*, **31**(4), 607–610.

Kanda, T, Joo, CY, & Choi, DW. 1975. Epidemiological studies on Malayan filariasis in an inland area in Kyungpook, Korea, 2: The periodicity of the microfilariae and the bionomics of the vector [Brugia malayi, *Anopheles sinensis*]. *Mosquito News*.

Kay, BH. 1979. Age structure of populations of *Culex annulirostris* (Diptera: Culicidae) at Kowanyama and Charleville, Queensland. *Journal of Medical Entomology*, **16**(4), 309–316.

Kenawy, Mohamed A. 1991. Development and survival of *Anopheles pharoensis* and An. multicolor from Faiyum, Egypt. *J Am Mosq Control Assoc*, **7**(4), 551–5.

Kitthawee, S, Edman, JD, & Upatham, ES. 1992. Relationship between female Anopheles dirus (Diptera: Culicidae) body size and parity in a biting population. *Journal of medical entomology*, **29**(6), 921–926.

Kohavi, Ron, *et al.* . 1995. A study of cross-validation and bootstrap for accuracy estimation and model selection. *Pages 1137–1145 of: Ijcai*, vol. 14.

Kraemer, Moritz UG, Sinka, Marianne E, Duda, Kirsten A, Mylne, Adrian QN, Shearer, Freya M, Barker, Christopher M, Moore, Chester G, Carvalho, Roberta G, Coelho, Giovanini E, Van Bortel, Wim, *et al.* . 2015. The global distribution of the arbovirus vectors *Aedes aegypti* and Ae. albopictus. *eLife*, **4**, e08347.

La Corte Dos Santos, Roseli, Forattini, Oswaldo Paulo, & Burattini, Marcelo Nascimento. 2004. *Anopheles albitarsis* sl (Diptera: Culicidae) survivorship and density in a rice irrigation area of the State of São Paulo, Brazil. *Journal of Medical Entomology*, **41**(5), 997–1000.

Lacroix, R, Delatte, Hélène, Hue, T, & Reiter, P. 2009. Dispersal and survival of male and female *Aedes albopictus* (Diptera: Culicidae) on Reunion Island. *Journal of Medical Entomology*, **46**(5), 1117–1124.

Lines, JD, Wilkes, TJ, & Lyimo, EO. 1991a. Human malaria infectiousness measured by age-specific sporozoite rates in *Anopheles gambiae* in Tanzania. *Parasitology*, **102**(2), 167–177.

Lines, JD, Curtis, CF, Wilkes, TJ, & Njunwa, KJ. 1991b. Monitoring human-biting mosquitoes (Diptera: Culicidae) in Tanzania with light-traps hung beside mosquito nets. *Bulletin of Entomological Research*, **81**(1), 77–84.

Loong, KP, Chiang, GL, Eng, KL, Chan, Seng Thim, Yap, HH, *et al.* . 1990. Survival and feeding behaviour of Malaysian strain of *Anopheles maculatus Theobald* (Diptera: Culicidae) and their role in malaria transmission. *Tropical Biomedicine*, **7**(1), 71–76.

Lorenz, Camila, Marques, Tatiani Cristina, Sallum, Maria Anice Mureb, & Suesdek, Lincoln. 2012. Morphometrical diagnosis of the malaria vectors *Anopheles cruzii*, *An. homunculus* and *An. bellator*. *Parasites & Vectors*, **5**(1), 257.

Lowe, RE, Ford, HR, Smittle, BJ, Weidhaas, DE, *et al.* . 1973. Reproductive behaviour of *Culex pipiens quinquefasciatus* released into a natural population. *Mosquito News*, **33**(2), 221–7.

Lutwama, JJ, & Mukwaya, LG. 1994. Mark-release-recapture studies on three anthropophilic populations of Aedes (Stegomyia) simpsoni comple (Diptera: Culicidae) in Uganda. *Bulletin of Entomological Research*, **84**(4), 521–527.

Macdonald, WW, Sebastian, A, & Tun, Maung Maung. 1968. A mark-release-recapture experiment with *Culex pipiens fatigans* in the village of Okpo, Burma. *Annals of Tropical Medicine & Parasitology*, **62**(2), 200–209.

Maciel-de Freitas, R, Codeco, CT, & Lourenço-de Oliveira, R. 2007. Body size-associated survival and dispersal rates of *Aedes aegypti* in Rio de Janeiro. *Medical and Veterinary Entomology*, **21**(3), 284–292.

Maciel-de Freitas, Rafael, Neto, Roman Brocki, Gonçalves, Jaylei Monteiro, Codeço, Claudia Torres, & Lourenço-de Oliveira, Ricardo. 2006. Movement of dengue vectors between the human modified environment and an urban forest in Rio de Janeiro. *Journal of Medical Entomology*, **43**(6), 1112–1120.

Maciel-De-Freitas, Rafael, Codeco, Claudia Torres, & Lourenco-De-Oliveira, Ricardo. 2007. Daily survival rates and dispersal of *Aedes aegypti* females in Rio de Janeiro, Brazil. *The American Journal of Tropical Medicine and Hygiene*, **76**(4), 659–665.

Maciel-de Freitas, Rafael, Eiras, Álvaro E, & Lourenço-de Oliveira, Ricardo. 2008. Calculating the survival rate and estimated population density of gravid *Aedes aegypti* (Diptera, Culicidae) in Rio de Janeiro, Brazil. *Cadernos de Saúde Pública*, **24**(12), 2747–2754.

Magesa, SM, Wilkes, TJ, Mnzava, AEP, Njunwa, KJ, Myamba, J, Kivuyo, MDP, Hill, N, Lines, JD, & Curtis, CF. 1991. Trial of pyrethroid impregnated bednets in an area of Tanzania holoendemic for malaria Part 2. Effects on the malaria vector population. *Acta Tropica*, **49**(2), 97–108.

Mahanta, B, Handique, R, Dutta, P, Narain, K, & Mahanta, J. 1999. Temporal variations in biting density and rhythm of *Culex quinquefasciatus* in tea agro-ecosystem of Assam, India.

Mahmood, Farida, & Reisen, William K. 1981. Duration of the gonotrophic cycles of *Anopheles culicifacies Giles* and *Anopheles stephensi Liston*, with observations on reproductive activity and survivorship during winter in Punjab province, Pakistan. *Mosquito News*.

Mala, Albert O, Irungu, Lucy W, Mitaki, Elizabeth K, Shililu, Josephat I, Mbogo, Charles M, Njagi, Joseph K, & Githure, John I. 2014. Gonotrophic cycle duration, fecundity and parity of *Anopheles gambiae* complex mosquitoes during an extended period of dry weather in a semi arid area in Baringo County, Kenya. *International Journal of Mosquito Research*, **1**(2), 28–34.

Marini, F, Caputo, B, Pombi, M, Tarsitani, G, & Della Torre, A. 2010. Study of *Aedes albopictus* dispersal in Rome, Italy, using sticky traps in mark–release–recapture experiments. *Medical and Veterinary Entomology*, **24**(4), 361–368.

Marsland, Stephen. 2015. *Machine learning: an algorithmic perspective.* CRC press.

Massey, NC, Garrod, G, Wiebe, A, Henry, AJ, Huang, Z, Moyes, CL, & Sinka, ME. 2016. A global bionomic database for the dominant vectors of human malaria. *Scientific data*, **3**, 160014.

McDonald, PT. 1977. Population characteristics of domestic *Aedes aegypti* (Diptera: Culicidae) in villages on the Kenya Coast I. Adult survivorship and population size. *Journal of Medical Entomology*, **14**(1), 42–48.

Mendis, C, Thompson, R, Begtrupl, K, Cuamba, N, & Dgedge, M. 1998. Cordon sanitaire or laissez faire: differential dispersal of young and old females of the malaria vector *Anopheles funestus* Giles (Diptera: Culicidae) in southern Mozambique. *African Entomology*, **6**(1).

Midega, Janet T, Mbogo, Charles M, Mwambi, Henr, Wilson, Michael D, Ojwang, Gordo, Mwangangi, Joseph M, Nzovu, Joseph G, Githure, John I, Yan, Guiyu, & Beier, John C. 2007. Estimating dispersal and survival of *Anopheles gambiae* and *Anopheles funestus* along the Kenyan coast by using mark–release–recapture methods. *Journal of Medical Entomology*, **44**(6), 923–929.

Milby, MM, & Reisen, WK. 1989. Estimation of vectorial capacity: vector survivorship. *Bulletin of the Society of Vector Ecology*, **14**(1), 47–54.

Mori, A, Wada, Y, *et al.* . 1977. The gonotrophic cycle of *Aedes albopictus* in the field. *Tropical Medicine*, **19**(3/4), 141–146.

Muir, Lynda E, & Kay, Brian H. 1998. *Aedes aegypti* survival and dispersal estimated by mark-release-recapture in northern Australia. *The American Journal of Tropical Medicine and Hygiene*, **58**(3), 277–282.

Müller, G, & Schlein, Y. 2005. Plant tissues: the frugal diet of mosquitoes in adverse conditions. *Medical and Veterinary Entomology*, **19**(4), 413–422.

Müller, Günter C, Beier, John C, Traore, Sekou F, Toure, Mahamoudou B, Traore, Mohamed M, Bah, Sekou, Doumbia, Seydou, & Schlein, Yosef. 2010. Successful field trial of attractive toxic sugar bait (ATSB) plant-spraying methods against malaria vectors in the *Anopheles gambiae* complex in Mali, West Africa. *Malaria Journal*, **9**(1), 210.

Nathan, Michael B. 1981. Bancroftian filariasis in coastal North Trinidad, West Indies: intensity of transmission by *Culex quinquefasciatus*. *Transactions of the Royal Society of Tropical Medicine and Hygiene*, **75**(5), 721–730.

Nayar, JK, Provost, MW, & Hansen, CW. 1980. Quantitative bionomics of *Culex nigripalpus* (Diptera: Culicidae) populations in Florida: 2. Distribution, dispersal and survival patterns. *Journal of Medical Entomology*, **17**(1), 40–50.

Nelson, RL, & Milby, MM. 1980. Dispersal and survival of field and laboratory strains of *Culex tarsalis* (Diptera: Culicidae). *Journal of Medical Entomology*, **17**(2), 146–150.

Nelson, RL, Milby, MM, Reeves, WC, & Fine, PEM. 1978. Estimates of survival, population size, and emergence of *Culex tarsalis* at an isolated site. *Annals of the Entomological Society of America*, **71**(5), 801–808.

Novoseltsev, Vasiliy N, Michalski, Anatoli I, Novoseltseva, Janna A, Yashin, Anatoliy I, Carey, James R, & Ellis, Alicia M. 2012. An age-structured extension to the vectorial capacity model. *PLoS One*, **7**(6).

Ordóñez González, José Genaro, Mercado Hernández, Roberto, Flores Suárez, Adriana Elizabeth, & Fernández Salas, Ildefonso. 2001. The use of sticky ovitraps to estimate dispersal of *Aedes aegypti* in northeastern Mexico. *Journal of the American Mosquito Control Association*, **17**(2), 93–97.

Pant, Chandra Prakash, World Health Organisation, *et al.* . 1962. Distribution of anophelines in relation to altitude in Nepal.

Pant, CP, & Yasuno, M. 1973. Field studies on the gonotrophic cycle of *Aedes aegypti* in Bangkok, Thailand. *Journal of Medical Entomology*, **10**(2), 219–223.

Penilla, RP, Rodriguez, MH, Lopez, AD, Viader-Salvadó, JM, & Sanchez, CN. 2002. Pteridine concentrations differ between insectary-reared and field-collected *Anopheles albimanus* mosquitoes of the same physiological age. *Medical and Veterinary Entomology*, **16**(3), 225–234.

Polovodova, VP. 1949. The determination of the physiological age of female *Anopheles* by the number of gonotrophic cycles completed. *Meditsinskaia Parazitologiia Parazitar Bolezni*, **18**, 352–355.

Pumpuni, CB, & Walker, ED. 1989. Population size and survivorship of adult Aedes triseriatus in a scrap tireyard in northern Indiana. *Journal of the American Mosquito Control Association*, **5**(2), 166–172.

Qualls, Whitney A, Müller, Günter C, Traore, Sekou F, Traore, Mohamed M, Arheart, Kristopher L, Doumbia, Seydou, Schlein, Yosef, Kravchenko, Vasiliy D, Xue, Rui-De, & Beier, John C. 2015. Indoor use of attractive toxic sugar bait (ATSB) to effectively control malaria vectors in Mali, West Africa. *Malaria Journal*, **14**(1), 301.

Quinones, ML, Lines, JD, Thomson, MC, Jawara, M, Morris, J, & Greenwood, BM. 1997. *Anopheles gambiae* gonotrophic cycle duration, biting and exiting behaviour unaffected by permethrin-impregnated bednets in The Gambia. *Medical and Veterinary Entomology*, **11**(1), 71–78.

Rajagopalan, PK. 1980. Population dynamics of culex pipiens fatigans, the filariasis vector, in pondicherry: influence of climate and environment. *Pages 745–52 of: Proceedings of the Indian National Sciences Academy*, vol. 46.

Ramaiah, KD, & Das, PK. 1992. Non-involvement of nulliparous females in the transmission of bancroftian filariasis. *Acta Tropica*, **52**(2-3), 149–153.

Rawlings, P, & Curtis, Chris F. 1982. Tests for the existence of genetic variability in the tendency of *Anopheles culicifacies* species B to rest in houses and to bite man. *Bulletin of the World Health Organisation*, **60**(3), 427.

Rawlings, P, & Davidson, G. 1982. The dispersal and survival of *Anopheles culicifacies Giles* (Diptera: Culicidae) in a Sri Lankan village under malathion spraying. *Bulletin of Entomological Research*, **72**(1), 139–144.

Rawlings, P, Curtis, CF, Wickramasinghe, MB, & Lines, J. 1981. The influence of age and season on dispersal and recapture of *Anopheles culicifacies* in Sri Lanka. *Ecological Entomology*, **6**(3), 307–319.

Reisen, William K, Mahmood, Farida, & Parveen, Tauheeda. 1979. *Anopheles subpictus Grassi*: observations on survivorship and population size using mark-release-recapture and dissection methods. *Researches on Population Ecology*, **21**(1), 12–29.

Reisen, William K, Mahmood, Farida, & Parveen, Tauheeda. 1980. *Anopheles culicifacies Giles*: a release-recapture experiment with cohorts of known age with implications for malaria epidemiology and genetical control in Pakistan. *Transactions of the Royal Society of Tropical Medicine and Hygiene*, **74**(3), 307–317.

Reisen, William K, Mahmood, Farida, & Azra, Khawar. 1981. *Anopheles culicifacies giles*: Adult ecological parameters measured in rural punjab province, pakistan using capture-mark-release-recapture and dissection methods, with comparative observations onAn. stephensi liston andAn. subpictus grassi. *Researches on Population Ecology*, **23**(1), 39–60.

Reisen, William K, Milby, Marilyn M, Reeves, William C, Meyer, Richard P, & Bock, Martha E. 1983. Population ecology of *Culex tarsalis* (Diptera: Culicidae) in a foothill environment of Kern County, California: temporal changes in female relative abundance, reproductive status, and survivorship. *Annals of the Entomological Society of America*, **76**(4), 800–808.

Reisen, WK, Sakai, RK, Baker, RH, Azra, K, Niaz, S, *et al.* . 1982. *Anopheles culicifacies*: observations on population ecology and reproductive behavior. *Mosquito News*, **42**(1), 93–101.

Reisen, WK, Yoshimura, G, Reeves, WC, Milby, MM, & Meyer, RP. 1984. The impact of aerial applications of ultra-low volume adulticides on *Culex tarsalis* populations (Diptera: Culicidae) in Kern County, California, USA, 1982. *Journal of Medical Entomology*, **21**(5), 573–585.

Renshaw, M, Service, MW, & Birley, MH. 1994. Host finding, feeding patterns and evidence for a memorized home range of the mosquito Aedes cantans. *Medical and Veterinary Entomology*, **8**(2), 187–193.

Rodriguez, Mario H, Bown, David N, Arredondo-Jimenez, Juan I, Villarreal, Cuauhtemoc, Loyola, Enrique G, & Frederickson, Christian E. 1992. Gonotrophic cycle and survivorship of *Anopheles albimanus* (Diptera: Culicidae) in southern Mexico. *Journal of Medical Entomology*, **29**(3), 395–399.

Rohatgi, Ankit. 2017 (January). *WebPlotDigitizer.*

Ross, Ronald. 1910. *The prevention of malaria.* Dutton.

Rúa, Guillermo L, Quiñones, Martha L, Vélez, Iván D, Zuluaga, Juan S, Rojas, William, Poveda, Germán, & Ruiz, Daniel. 2005. Laboratory estimation of the effects of increasing temperatures on the duration of gonotrophic cycle of *Anopheles albimanus* (Diptera: Culicidae). *Memorias Do Instituto Oswaldo Cruz*, **100**(5), 515–520.

Russell, Richard C. 1986. *Culex annulirostris Skuse* (Diptera: Culicidae) at Appin, NSW?bionomics and behaviour. *Austral Entomology*, **25**(2), 103–109.

Samarawickrema, WA. 1967. A study of the age-composition of natural populations of *Culex pipiens fatigans Wiedemann* in relation to the transmission of filariasis due to Wuchereria bancrofti (Cobbold) in Ceylon. *Bulletin of the World Health Organisation*, **37**(1), 117.

Samarawickrema, WA. 1968. Biting cycles and parity of the mosquito *Mansonia* (Mansonioides) uniformis (Theo.) in Ceylon. *Bulletin of Entomological research*, **58**(02), 299–314.

Samarawickrema, WA, Sone, Fola, & Cummings, RF. 1987. Seasonal abundance, diel biting activity and parity of *Aedes polynesiensis marks* and *A. samoanus* (Grünberg) (Diptera: Culicidae) in Samoa. *Bulletin of Entomological Research*, **77**(2), 191–200.

Schlein, Yosef, & Müller, Gunter C. 2008. An approach to mosquito control: Using the dominant attraction of flowering Tamarix jordanis trees against *Culex pipiens*. *Journal of Medical Entomology*, **45**(3), 384–390.

Schlein, Yosef, & Müller, Günter C. 2012. Diurnal resting behavior of adult *Culex pipiens* in an arid habitat in Israel and possible control measurements with toxic sugar baits. *Acta Tropica*, **124**(1), 48–53.

Schlein, Yosef, & Müller, Günter C. 2015. Decrease of larval and subsequent adult *Anopheles sergentii* populations following feeding of adult mosquitoes from Bacillus sphaericus-containing attractive sugar baits. *Parasites & Vectors*, **8**(1), 244.

Scholl, PJ, Porter, CHARLES H, & Defoliart, GENE R. 1979. *Aedes triseriatus*: persistence of nulliparous females under field conditions. *Mosquito News*, **39**, 368–371.

Sempala, SDK. 1981. The ecology of Aedes (Stegomyia) africanus (Theobald) in a tropical forest in Uganda: mark-release-recapture studies on a female adult population. *International Journal of Tropical Insect Science*, **1**(3), 211–224.

Service, MW, Barnish, G, & Toure, YT. 1995. Vectorial capacity and entomological inoculation rates of Anopheles gambiae in a high rainfall forested area of southern Sierra Leone. *Trop Med Parasitol*, **46**, 164–171.

Shalaby, AM, World Health Organisation, *et al.* . 1962. Studies on the age composition of *Anopheles Culicifacies Giles* at different phases of the development of resistance to DDT in Panchmahals district of Gujarat State, India.

Sheppard, PM, Macdonald, WW, Tonn, RJ, & Grab, B. 1969. The dynamics of an adult population of *Aedes aegypti* in relation to dengue haemorrhagic fever in Bangkok. *The Journal of Animal Ecology*, 661–702.

Shriram, AN, & Krishnamoorthy, K. 2011. Population dynamics, age composition and survival of Downsiomyia nivea in relation to transmission of diurnally subperiodic filariasis. *Journal of Asia-Pacific Entomology*, **14**(1), 34–40.

Silver, John B. 2007. *Mosquito ecology: field sampling methods.* Springer Science & Business Media.

Sinka, Marianne E, Bangs, Michael J, Manguin, Sylvie, Coetzee, Maureen, Mbogo, Charles M, Hemingway, Janet, Patil, Anand P, Temperley, Will H, Gething, Peter W, Kabaria, Caroline W, Okara, Robi M, Boeckel, Thomas Van, Godfray, H Charles J, Harbach, Ralph E, & Hay, Simon I. 2010. The dominant *Anopheles* vectors of human malaria in Africa, Europe and the Middle East: occurrence data, distribution maps and bionomic précis. *Parasites & Vectors*, **3**(1), 117.

Sinka, Marianne E, Bangs, Michael J, Manguin, Sylvie, Rubio-Palis, Yasmin, Chareonviriyaphap, Theeraphap, Coetzee, Maureen, Mbogo, Charles M, Hemingway, Janet, Patil, Anand P, Temperley, William H, *et al.* . 2012. A global map of dominant malaria vectors. *Parasites & Vectors*, **5**(1), 1.

Smith, David L, & McKenzie, F Ellis. 2004. Statics and dynamics of malaria infection in *Anopheles* mosquitoes. *Malaria Journal*, **3**(1), 13.

Smith, Gordon E, Watson, Robert B, & Cromwell, Robert L. 1941. Observations on the flight range of Anopheles quadrimaculatus, Say. *American Journal of Epidemiology*, **34**(2), 102–113.

Smith, Stephen M, & Kurtz, RM. 1994. The age structure of a population of *Aedes provocans* (Diptera: Culicidae) in southwestern Ontario. *Great Lakes Entomologist*, **27**(2), 113.

Snow, WF, & Boreham, PFL. 1978. The host-feeding patterns of some culicine mosquitoes (Diptera: Culicidae) in The Gambia. *Bulletin of Entomological Research*, **68**(4), 695–706.

Stan Development Team. 2014. *Stan: A C++ Library for Probability and Sampling, Version 2.5.0.*

Styer, Linda M, Carey, James R, Wang, Jane-Ling, & Scott, Thomas W. 2007. Mosquitoes do senesce: departure from the paradigm of constant mortality. *The American Journal of Tropical Medicine and Hygiene*, **76**(1), 111–117.

Surendran, SN, Ramasamy, MS, De Silva, BGDNK, & Ramasamy, R. 2006. *Anopheles culicifacies* sibling species B and E in Sri Lanka differ in longevity and in their susceptibility to malaria parasite infection and common insecticides. *Medical and Veterinary Entomology*, **20**(1), 153–156.

Suzuki, Takeshi. 1978. Preliminary studies on blood meal interval of *Aedes polynesiensis* in the field. *Medical Entomology and Zoology*, **29**(2), 169–174.

Takagi, Masahiro, Tsuda, Yoshio, Suzuki, Akemi, & Wada, Yoshito. 1995. Movement of individually marked *Aedes albopictus* females in Nagasaki, Japan. *Tropical Medicine*, **37**(2), 79–85.

Takken, W, Charlwood, JD, Billingsley, PF, & Gort, G. 1998. Dispersal and survival of *Anopheles funestus* and *A. gambiae* sl (Diptera: Culicidae) during the rainy season in southeast Tanzania. *Bulletin of Entomological Research*, **88**(5), 561–566.

Touré, Yeya T, Dolo, Guimogo, Petrarca, Vincenzo, Dao, Adama, Carnahan, John, & Taylor, Charles E. 1998. Mark–release–recapture experiments with *Anopheles gambiae* sl in Banambani Village, Mali, to determine population size and structure. *Medical and Veterinary Entomology*, **12**(1), 74–83.

Trpis, Milan, & Hausermann, Walter. 1975. Demonstration of differential domesticity of *Aedes aegypti* (L.)(Diptera, Culicidae) in Africa by mark-release-recapture. *Bulletin of Entomological Research*, **65**(2), 199–208.

Trpis, Milan, Häusermann, Walter, & Craig Jr, George B. 1995. Estimates of population size, dispersal, and longevity of domestic *Aedes aegypti* aegypti (Diptera: Culicidae) by mark–release–recapture in the village of Shauri Moyo in eastern Kenya. *Journal of Medical Entomology*, **32**(1), 27–33.

Tsuda, Yoshio, Takagi, M, Wang, S, Wang, Z, & Tang, L. 2001. Movement of *Aedes aegypti* (Diptera: Culicidae) released in a small isolated village on Hainan Island, China. *Journal of Medical Entomology*, **38**(1), 93–98.

Tuchinda, P, Kitaoka, Masami, Ogata, Takayuki, & Kurihara, T. 1969. On the diurnal rhythmus of biting behavior of Aëdes aegypti in relation to the age and to the hemorrhagic fever in Bangkok, 1964. *Japanese Journal of Tropical Medicine*, **10**(1), 1–6.

Uttah, Emmanuel C, Iboh, Cletus I, Ajang, Raymond, Osim, SE, & Etta, Hannah. 2013. Physiological age composition of female anopheline mosquitoes in an area endemic for malaria and filariasis. *International Journal of Scientific and Research Publications*, **3**(7), 1–4.

Valerio, Laura, Facchinelli, Luca, Ramsey, Janine M, & Scott, Thomas W. 2012. Dispersal of male *Aedes aegypti* in a coastal village in southern Mexico. *The American Journal of Tropical Medicine and Hygiene*, **86**(4), 665–676.

Vehtari, Aki, Gelman, Andrew, & Gabry, Jonah. 2015. Efficient implementation of leave-one-out cross-validation and WAIC for evaluating fitted Bayesian models. *arXiv preprint arXiv:1507.04544*.

Vythilingam, Indra, Oda, Kazumasa, Mahadevan, S, Abdullah, Ghani, Thim, Chan Seng, Hong, Choo Choon, Vijayamalar, B, Sinniah, Mangalam, & Igarashi, Akira. 1997. Abundance, parity, and Japanese encephalitis virus infection of mosquitoes (Diptera: Culicidae) in Sepang District, Malaysia. *Journal of Medical Entomology*, **34**(3), 257–262.

Wada, Yoshito, Kawai, Senji, Oda, Tsutomu, Miyagi, Ichiro, Suenaga, Osamu, Nishigaki, Jojiro, Omori, Nanzaburo, Takahashi, Katsumi, Matsuo, Reizo, Itoh, Tatsuya, *et al.* . 1969. Dispersal experiment of *Culex tritaeniorhynchus* in Nagasaki area (Preliminary report). *Tropical Medicine*, **11**(1), 37–44.

Watson, Tonya M, Saul, Allan, & Kay, Brian H. 2000. Aedes notoscriptus (Diptera: Culicidae) survival and dispersal estimated by mark-release-recapture in Brisbane, Queensland, Australia. *Journal of Medical Entomology*, **37**(3), 380–384.

Wilkes, TJ, Matola, YG, & Charlwood, JD. 1996. *Anopheles rivulorum*, a vector of human malaria in Africa. *Medical and Veterinary Entomology*, **10**(1), 108–110.

Wong, Jacklyn, Astete, Helvio, Morrison, Amy C, & Scott, Thomas W. 2014. Sampling considerations for designing *Aedes aegypti* (Diptera: Culicidae) oviposition studies in Iquitos, Peru: substrate preference, diurnal periodicity, and gonotrophic cycle length. *Journal of Medical Entomology*, **48**(1), 45–52.

World Health Organisation, *et al.* . 1960. Preliminary appraisal on the use of age-grouping methods in anopheline mosquitos.

Yasuno, M, Rajagopalan, PK, Laüreque, GC, *et al.* . 1975. Migration patterns of *Culex fatigans* around Delhi, India. *Tropical Medicine*, **17**(2), 91–96.

Zetek, James. 1915. Behavior of *Anopheles albimanus Wiede* and *tarsimaculata Goeldi. Annals of the Entomological Society of America*, **8**(3), 221–271.
