## Supplementary figures for "A Meta-analysis of Longevity Estimates of Mosquito Vectors of Disease"

### A Meta-analysis of Longevity Estimates of Mosquito Vectors of Disease: Supplementary Figures

May 29, 2022

#### S1 Lifespan estimates

| Analysis | Grouping | Log-likelihood | z | p |
|---|---|---|---|---|
| MRR | genus | -7315.2 | - | - |
| | overall | -7511.0 | 11.3 | 0.0 |
| | species | -7748.9 | 5.5 | 0.0 |
| Detinova | species | -14,672.4 | - | - |
| | species-complex | -15,504.9 | 3.6 | 0.0 |
| | continent | -16,731.1 | 0.4 | 0.3 |
| Polovodova | species | -3818.3 | - | - |
| | genus | -3848.3 | 0.4 | 0.3 |
| | overall | -3895.5 | 0.4 | 0.4 |

Table S1: **Model comparison by K-fold cross-validation.** Here, we present the estimates of model fit on hold-out datasets (see SOM) as measured by the sum of the pointwise log-likelihood values averaged across all posterior draws (in the "Log-likelihood" column) across each of the three analyses and each grouping of studies considered. Within each row-block ("MRR", "Detinova", "Polovodova"), the models are arranged according to the point estimate of the fit: best at top; worst at bottom. The $z$ scores show the scaled difference in the averaged pointwise log-likelihood (Vehtari *et al.* , 2015) versus the model immediately above it in the ranking, and the "p" column shows the corresponding $p$ values representing the cumulative density function from a standard normal.

| Genus | Species | 5% | 25% | 50% | 75% | 95% | Mean | Std. dev. |
|---|---|---|---|---|---|---|---|---|
| *Aedes* | *simpsoni s.l.* | 6.3 | 11.3 | 18.3 | 31.8 | 75.0 | 26.9 | 29.5 |
| *Aedes* | *communis* | 10.0 | 13.2 | 15.4 | 18.2 | 24.3 | 16.2 | 5.2 |
| *Anopheles* | *koliensis* | 6.7 | 10.5 | 15.2 | 24.0 | 56.5 | 21.9 | 25.2 |
| *Anopheles* | *punctulatus* | 7.2 | 10.7 | 15.1 | 22.8 | 48.9 | 20.3 | 28.6 |
| *Aedes* | *cantans* | 9.7 | 12.0 | 13.9 | 16.1 | 20.6 | 14.4 | 4.7 |
| *Aedes* | *albopictus* | 8.0 | 10.0 | 11.6 | 13.7 | 17.7 | 12.1 | 3.4 |
| *Anopheles* | *albimanus* | 5.9 | 8.6 | 11.6 | 16.9 | 35.7 | 15.2 | 18.7 |
| *Anopheles* | *darlingi* | 4.9 | 7.3 | 10.2 | 16.0 | 37.5 | 14.5 | 14.8 |
| *Anopheles* | *maculatus* | 5.7 | 7.9 | 10.2 | 13.7 | 24.3 | 12.2 | 8.7 |
| *Anopheles* | *culicifacies s.l.* | 4.6 | 7.3 | 9.1 | 11.4 | 18.2 | 10.1 | 6.9 |
| *Anopheles* | *sergenti* | 3.6 | 5.9 | 8.7 | 13.8 | 32.6 | 12.4 | 13.4 |
| *Aedes* | *triseriatus* | 5.6 | 6.6 | 7.5 | 8.5 | 10.7 | 7.7 | 1.8 |
| *Anopheles* | *farauti s.l.* | 4.2 | 5.4 | 6.4 | 7.7 | 10.5 | 6.8 | 2.1 |
| *Aedes* | *africanus* | 3.4 | 5.1 | 6.2 | 7.8 | 11.9 | 7.0 | 9.3 |
| *Aedes* | *aegypti* | 2.8 | 4.5 | 6.2 | 8.5 | 13.9 | 7.0 | 3.8 |
| *Culex* | *quinquefasciatus* | 4.3 | 5.2 | 5.9 | 7.0 | 9.4 | 6.3 | 1.8 |
| *Culex* | *nigripalpus* | 3.5 | 4.6 | 5.8 | 7.5 | 11.3 | 6.4 | 3.1 |
| *Aedes* | *notoscriptus* | 2.9 | 4.0 | 5.4 | 8.2 | 20.2 | 7.9 | 9.5 |
| *Culex* | *pipiens* | 3.4 | 4.4 | 5.2 | 6.4 | 9.0 | 5.6 | 2.0 |
| *Anopheles* | *saperoi* | 3.5 | 4.2 | 4.9 | 5.8 | 8.2 | 5.2 | 2.0 |
| *Anopheles* | *gambiae s.l.* | 3.0 | 3.8 | 4.4 | 5.1 | 6.4 | 4.5 | 1.1 |
| *Aedes* | *melanimon* | 3.3 | 3.9 | 4.3 | 4.8 | 5.7 | 4.4 | 1.0 |
| *Anopheles* | *funestus s.l.* | 2.9 | 3.6 | 4.2 | 4.8 | 6.0 | 4.3 | 1.0 |
| *Anopheles* | *lesteri* | 2.4 | 3.2 | 3.9 | 5.0 | 7.5 | 4.4 | 2.1 |
| *Anopheles* | *quadrimaculatus* | 3.3 | 3.6 | 3.9 | 4.2 | 4.8 | 4.0 | 0.5 |
| *Anopheles* | *pseudopunctipennis s.l.* | 2.3 | 2.9 | 3.5 | 4.3 | 6.7 | 3.8 | 1.9 |
| *Aedes* | *vexans* | 1.9 | 2.3 | 2.6 | 3.0 | 4.1 | 2.8 | 1.0 |
| *Culex* | *tarsalis* | 1.9 | 2.1 | 2.3 | 2.4 | 2.7 | 2.3 | 0.3 |
| *Anopheles* | *vestitipennis* | 1.3 | 1.8 | 2.2 | 2.9 | 4.4 | 2.5 | 1.2 |
| *Anopheles* | *albitarsis s.l.* | 1.3 | 1.6 | 1.9 | 2.2 | 2.9 | 2.0 | 1.0 |
| *Culex* | *tritaeniorhynchus* | 0.7 | 0.9 | 1.1 | 1.2 | 1.6 | 1.1 | 1.0 |
| *Anopheles* | *subpictus s.l.* | 0.4 | 0.6 | 0.8 | 1.1 | 2.4 | 1.1 | 2.6 |
| *Aedes* | | 2.8 | 4.8 | 6.9 | 10.0 | 17.2 | 8.1 | 4.9 |
| *Anopheles* | | 1.5 | 3.0 | 5.0 | 8.4 | 17.8 | 6.8 | 6.4 |
| *Culex* | | 1.1 | 1.8 | 2.5 | 3.5 | 5.6 | 2.9 | 1.5 |
| **Overall** | | 1.4 | 2.8 | 4.6 | 7.5 | 15.4 | 6.0 | 5.2 |

Table S2: **MRR: mean unfed female mosquito lifespan estimates.** The 5%, 25%, 50%, 75% and 95% columns indicate the respective quantiles of the posterior distribution for mean lifespan, and the 'Mean' and 'Std. dev.' columns indicate the posterior mean and standard deviation of mean lifespan. Units of lifespan are days.

| Genus | Species | 5% | 25% | 50% | 75% | 95% | Mean | Std. dev. |
|---|---|---|---|---|---|---|---|---|
| *Anopheles* | *gambiae (Form M)* | 0.9 | 1.8 | 2.6 | 3.6 | 6.7 | 3.2 | 4.6 |
| *Anopheles* | *gambiae* | 0.6 | 1.1 | 1.7 | 2.7 | 5.6 | 2.2 | 2.2 |
| *Anopheles* | *arabiensis* | 0.4 | 0.9 | 1.5 | 2.6 | 6.3 | 2.3 | 3.9 |
| *Anopheles* | *gambiae s.l.* | 0.7 | 1.7 | 3.1 | 6.3 | 21.2 | 7.3 | 56.1 |
| *Anopheles* | *moucheti* | 1.0 | 1.9 | 2.8 | 4.2 | 8.9 | 3.8 | 6.3 |
| *Anopheles* | *funestus* | 0.8 | 1.6 | 2.7 | 4.4 | 10.6 | 4.0 | 10.6 |
| *Anopheles* | *funestus s.l.* | 0.6 | 1.2 | 2.1 | 3.7 | 9.8 | 3.3 | 4.8 |
| *Anopheles* | *nili s.l.* | 1.0 | 1.4 | 1.7 | 2.1 | 2.8 | 1.8 | 0.6 |
| *Anopheles* | *subpictus s.l.* | 0.9 | 1.5 | 2.1 | 2.8 | 4.8 | 2.4 | 1.8 |
| *Anopheles* | *dirus (formerly Sp. A)* | 0.5 | 1.0 | 1.5 | 2.2 | 4.2 | 1.8 | 1.5 |
| *Anopheles* | *dirus s.l.* | 0.6 | 1.2 | 2.0 | 3.1 | 7.4 | 2.9 | 6.8 |
| *Anopheles* | *aconitus* | 0.5 | 1.1 | 1.8 | 2.8 | 6.3 | 2.5 | 4.7 |
| *Anopheles* | *sundaicus s.l.* | 0.4 | 0.9 | 1.6 | 2.9 | 7.5 | 2.7 | 7.4 |
| *Anopheles* | *minimus s.l.* | 0.5 | 1.1 | 1.6 | 2.5 | 5.8 | 2.3 | 5.6 |
| *Anopheles* | *fluviatilis s.l.* | 0.4 | 0.9 | 1.4 | 2.3 | 5.1 | 2.0 | 3.0 |
| *Anopheles* | *maculatus s.l.* | 0.6 | 1.0 | 1.2 | 1.6 | 2.5 | 1.4 | 0.8 |
| *Anopheles* | *farauti s.l.* | 0.3 | 0.7 | 1.2 | 2.0 | 4.9 | 1.8 | 2.3 |
| *Anopheles* | *annularis s.l.* | 0.3 | 0.7 | 1.2 | 2.0 | 4.7 | 1.7 | 1.9 |
| *Anopheles* | *sinensis* | 0.4 | 0.8 | 1.1 | 1.7 | 3.0 | 1.4 | 1.0 |
| *Anopheles* | *culicifacies s.l.* | 0.5 | 0.8 | 1.1 | 1.5 | 2.3 | 1.2 | 0.6 |
| *Anopheles* | *anthropophagus (lesteri)* | 0.4 | 0.9 | 1.4 | 2.3 | 5.1 | 2.0 | 3.0 |
| *Anopheles* | *lesteri s.l.* | 0.3 | 0.6 | 1.1 | 1.9 | 4.7 | 1.7 | 2.4 |
| *Anopheles* | *barbirostris s.l.* | 0.4 | 0.7 | 1.0 | 1.4 | 2.8 | 1.2 | 1.3 |
| *Anopheles* | *albitarsis (formerly Sp C)* | 0.8 | 2.0 | 3.9 | 8.4 | 32.6 | 13.3 | >100 |
| *Anopheles* | *albitarsis (formerly Sp A)* | 0.2 | 0.4 | 0.6 | 1.0 | 2.3 | 0.8 | 0.9 |
| *Anopheles* | *albitarsis (Sp. B)* | 0.1 | 0.3 | 0.6 | 0.9 | 1.9 | 0.7 | 0.8 |
| *Anopheles* | *albitarsis s.l.* | 0.4 | 0.9 | 1.5 | 3.0 | 8.3 | 2.8 | 5.8 |
| *Anopheles* | *nuneztovari s.l.* | 0.3 | 0.6 | 1.1 | 1.9 | 4.4 | 1.6 | 2.3 |
| *Anopheles* | *darlingi* | 0.2 | 0.7 | 1.2 | 2.4 | 6.7 | 2.2 | 4.2 |
| *Anopheles* | *albimanus* | 0.2 | 0.5 | 0.9 | 1.6 | 3.6 | 1.3 | 1.6 |
| *Anopheles* | *aquasalis* | 0.3 | 0.6 | 1.1 | 1.8 | 4.4 | 1.6 | 4.0 |
| *Anopheles* | *aquasalis s.l.* | 0.2 | 0.4 | 0.7 | 1.0 | 2.4 | 0.9 | 1.3 |

Table S3: **Detinova dissection: mean female mosquito lifespan estimates in terms of gonotrophic cycles.** The 5%, 25%, 50%, 75% and 95% columns indicate the respective quantiles of the posterior distribution for mean lifespan, and the 'Mean' and 'Std. dev.' columns indicate the posterior mean and standard deviation of mean lifespan.

| Genus | Species | 5% | 25% | 50% | 75% | 95% | Mean | Std. dev. |
|---|---|---|---|---|---|---|---|---|
| *Anopheles* | *sergentii* | 1.0 | 1.9 | 2.5 | 3.4 | 6.2 | 3.0 | 2.5 |
| *Anopheles* | *gambiae s.l.* | 0.7 | 1.2 | 1.9 | 2.9 | 5.4 | 2.4 | 1.9 |
| *Culex* | *thalassius* | 1.1 | 1.6 | 1.8 | 2.1 | 2.9 | 1.9 | 1.0 |
| *Anopheles* | *farauti s.l.* | 1.5 | 1.6 | 1.7 | 1.7 | 1.9 | 1.7 | 0.1 |
| *Anopheles* | *minimus* | 0.7 | 1.2 | 1.7 | 2.2 | 4.1 | 2.0 | 2.2 |
| *Anopheles* | *maculipennis* | 0.8 | 1.1 | 1.4 | 1.8 | 2.9 | 1.6 | 0.7 |
| *Culex* | *pipiens* | 1.1 | 1.3 | 1.4 | 1.5 | 1.7 | 1.4 | 0.2 |
| *Mansonia* | *uniformis* | 0.9 | 1.1 | 1.1 | 1.2 | 1.4 | 1.1 | 0.2 |
| *Anopheles* | *rivulorum* | 0.8 | 1.0 | 1.1 | 1.1 | 1.4 | 1.1 | 0.2 |
| *Anopheles* | *melas* | 0.8 | 1.0 | 1.0 | 1.1 | 1.3 | 1.0 | 0.2 |
| *Anopheles* | *culicifacies* | 0.9 | 1.0 | 1.0 | 1.1 | 1.2 | 1.0 | 0.1 |
| *Anopheles* | *subpictus* | 0.6 | 0.9 | 1.0 | 1.2 | 2.1 | 1.2 | 2.0 |
| *Aedes* | *aegypti* | 0.5 | 0.8 | 1.0 | 1.2 | 2.0 | 1.2 | 3.0 |
| *Anopheles* | *quadrimaculatus* | 0.7 | 0.9 | 1.0 | 1.1 | 1.3 | 1.0 | 0.2 |
| *Anopheles* | *stephensi* | 0.6 | 0.8 | 1.0 | 1.1 | 1.6 | 1.0 | 0.4 |
| *Anopheles* | *darlingi* | 0.7 | 0.8 | 0.9 | 1.0 | 1.3 | 1.0 | 0.2 |
| *Culex* | *quinquefasciatus* | 0.4 | 0.7 | 0.9 | 1.3 | 2.2 | 1.1 | 0.6 |
| *Aedes* | *polynesiensis* | 0.8 | 0.9 | 0.9 | 0.9 | 1.0 | 0.9 | 0.1 |
| *Culex* | *tritaeniorhynchus* | 0.3 | 0.6 | 0.9 | 1.3 | 4.2 | 2.5 | 30.8 |
| *Culex* | *annulirostris* | 0.5 | 0.8 | 0.9 | 1.0 | 1.4 | 0.9 | 0.5 |
| *Aedes* | *samoanus* | 0.6 | 0.7 | 0.8 | 0.9 | 1.0 | 0.8 | 0.2 |
| *Aedes* | *sollicitans* | 0.4 | 0.5 | 0.6 | 0.8 | 1.3 | 0.7 | 0.5 |
| *Aedes* | *vexans* | 0.4 | 0.5 | 0.6 | 0.7 | 0.9 | 0.6 | 0.7 |
| *Anopheles* | *cruzii* | 0.3 | 0.5 | 0.5 | 0.7 | 1.1 | 0.6 | 0.4 |
| *Anopheles* | *bellator* | 0.3 | 0.4 | 0.5 | 0.6 | 1.0 | 0.6 | 1.8 |
| *Anopheles* | | 0.6 | 1.0 | 1.4 | 1.9 | 3.2 | 1.6 | 0.9 |
| *Mansonia* | | 0.9 | 1.1 | 1.1 | 1.2 | 1.4 | 1.1 | 0.2 |
| *Culex* | | 0.5 | 0.8 | 1.0 | 1.4 | 2.3 | 1.2 | 0.6 |
| *Aedes* | | 0.6 | 0.7 | 0.8 | 0.9 | 1.1 | 0.8 | 0.2 |
| **Overall** | | 0.5 | 0.8 | 1.2 | 1.7 | 2.7 | 1.3 | 0.7 |

Table S4: **Polovodova dissection: mean female mosquito lifespan in terms of gonotrophic cycles.** The 5%, 25%, 50%, 75% and 95% columns indicate the respective quantiles of the posterior distribution for mean lifespan, and the 'Mean' and 'Std. dev.' columns indicate the posterior mean and standard deviation of mean lifespan.

| Genus | Species | 5% | 25% | 50% | 75% | 95% | Mean | Std. dev. |
|---|---|---|---|---|---|---|---|---|
| *Anopheles* | *gambiae (Form M)* | 3.3 | 6.2 | 8.7 | 11.9 | 22.3 | 10.7 | 14.9 |
| *Anopheles* | *gambiae* | 2.0 | 3.8 | 5.8 | 9.0 | 18.4 | 7.6 | 7.1 |
| *Anopheles* | *arabiensis* | 1.4 | 3.2 | 5.3 | 8.8 | 20.8 | 7.7 | 11.9 |
| *Anopheles* | *gambiae s.l.* | 2.5 | 5.7 | 10.3 | 20.6 | 68.4 | 23.9 | >100 |
| *Anopheles* | *moucheti* | 3.5 | 6.4 | 9.2 | 13.9 | 29.0 | 12.5 | 19.7 |
| *Anopheles* | *funestus* | 2.9 | 5.7 | 9.0 | 14.5 | 34.4 | 13.4 | 31.0 |
| *Anopheles* | *funestus s.l.* | 2.0 | 4.3 | 7.1 | 12.5 | 32.2 | 11.1 | 15.6 |
| *Anopheles* | *nili s.l.* | 3.6 | 4.8 | 5.8 | 7.1 | 9.6 | 6.1 | 2.0 |
| *Anopheles* | *subpictus s.l.* | 3.4 | 5.3 | 7.1 | 9.5 | 16.2 | 8.2 | 6.2 |
| *Anopheles* | *dirus (formerly Sp. A)* | 1.9 | 3.6 | 5.1 | 7.4 | 13.8 | 6.2 | 5.0 |
| *Anopheles* | *dirus s.l.* | 2.0 | 4.4 | 6.8 | 10.5 | 24.1 | 9.8 | 23.0 |
| *Anopheles* | *aconitus* | 1.9 | 4.0 | 6.2 | 9.6 | 20.7 | 8.5 | 15.4 |
| *Anopheles* | *sundaicus s.l.* | 1.5 | 3.4 | 5.6 | 9.7 | 24.7 | 9.1 | 23.6 |
| *Anopheles* | *minimus s.l.* | 1.9 | 3.8 | 5.5 | 8.3 | 19.2 | 7.9 | 17.8 |
| *Anopheles* | *fluviatilis s.l.* | 1.6 | 3.2 | 5.0 | 7.8 | 16.9 | 6.8 | 9.6 |
| *Anopheles* | *maculatus s.l.* | 2.2 | 3.5 | 4.5 | 5.6 | 8.6 | 4.9 | 2.6 |
| *Anopheles* | *farauti s.l.* | 1.2 | 2.6 | 4.3 | 6.9 | 16.0 | 6.1 | 7.5 |
| *Anopheles* | *annularis s.l.* | 1.1 | 2.5 | 4.3 | 6.9 | 15.6 | 5.8 | 6.2 |
| *Anopheles* | *sinensis* | 1.6 | 2.8 | 4.1 | 5.9 | 10.3 | 4.8 | 3.2 |
| *Anopheles* | *culicifacies s.l.* | 1.9 | 3.0 | 4.0 | 5.2 | 8.0 | 4.3 | 2.0 |
| *Anopheles* | *anthropophagus (lesteri)* | 1.6 | 3.2 | 5.0 | 7.7 | 16.9 | 6.8 | 9.9 |
| *Anopheles* | *lesteri s.l.* | 0.9 | 2.2 | 3.9 | 6.5 | 15.6 | 5.7 | 7.8 |
| *Anopheles* | *barbirostris s.l.* | 1.4 | 2.5 | 3.5 | 4.9 | 9.6 | 4.3 | 4.1 |
| *Anopheles* | *albitarsis (formerly Sp C)* | 3.0 | 6.8 | 13.0 | 27.3 | >100 | 44.2 | >100 |
| *Anopheles* | *albitarsis (formerly Sp A)* | 0.6 | 1.3 | 2.2 | 3.7 | 7.7 | 3.0 | 3.1 |
| *Anopheles* | *albitarsis (Sp. B)* | 0.5 | 1.2 | 2.0 | 3.3 | 6.6 | 2.6 | 2.6 |
| *Anopheles* | *albitarsis s.l.* | 1.2 | 3.1 | 5.4 | 9.9 | 26.9 | 9.3 | 18.6 |
| *Anopheles* | *nuneztovari s.l.* | 1.0 | 2.3 | 3.9 | 6.4 | 14.7 | 5.6 | 7.6 |
| *Anopheles* | *darlingi* | 0.9 | 2.4 | 4.5 | 8.1 | 22.0 | 7.4 | 13.6 |
| *Anopheles* | *albimanus* | 0.9 | 2.0 | 3.3 | 5.4 | 12.0 | 4.5 | 5.3 |
| *Anopheles* | *aquasalis* | 1.0 | 2.3 | 3.9 | 6.3 | 14.4 | 5.6 | 12.9 |
| *Anopheles* | *aquasalis s.l.* | 0.8 | 1.6 | 2.4 | 3.8 | 8.2 | 3.3 | 4.1 |

Table S5: **Detinova dissection: mean female mosquito lifespan estimates in terms of chronological time.** The 5%, 25%, 50%, 75% and 95% columns indicate the respective quantiles of the posterior distribution for mean lifespan, and the 'Mean' and 'Std. dev.' columns indicate the posterior mean and standard deviation of mean lifespan. Units of lifespan are days.

| **Genus** | **Species** | **5%** | **25%** | **50%** | **75%** | **95%** | **Mean** | **Std. dev.** |
|---|---|---|---|---|---|---|---|---|
| *Culex* | *thalassius* | 5.7 | 8.2 | 9.4 | 11.0 | 15.6 | 10.1 | 5.2 |
| *Anopheles* | *sergentii* | 3.7 | 6.4 | 8.5 | 11.5 | 20.6 | 10.1 | 8.1 |
| *Culex* | *pipiens* | 5.7 | 6.6 | 7.2 | 7.8 | 8.9 | 7.2 | 1.0 |
| *Anopheles* | *gambiae s.l.* | 2.4 | 4.3 | 6.4 | 9.8 | 18.3 | 8.0 | 6.3 |
| *Anopheles* | *farauti s.l.* | 4.8 | 5.5 | 5.9 | 6.3 | 6.9 | 5.9 | 0.6 |
| *Anopheles* | *minimus* | 2.7 | 4.4 | 5.8 | 7.7 | 13.7 | 6.9 | 8.1 |
| *Anopheles* | *maculipennis* | 2.8 | 4.0 | 5.1 | 6.4 | 9.6 | 5.5 | 2.3 |
| *Culex* | *quinquefasciatus* | 2.0 | 3.4 | 4.8 | 6.9 | 11.7 | 5.6 | 3.3 |
| *Aedes* | *aegypti* | 2.3 | 3.7 | 4.7 | 5.6 | 9.3 | 5.6 | 12.3 |
| *Culex* | *annulirostris* | 2.7 | 3.9 | 4.6 | 5.4 | 7.5 | 4.9 | 2.4 |
| *Culex* | *tritaeniorhynchus* | 1.6 | 3.2 | 4.6 | 7.2 | 22.8 | 13.0 | 174.0 |
| *Mansonia* | *uniformis* | 3.4 | 4.0 | 4.4 | 4.9 | 5.5 | 4.5 | 0.7 |
| *Aedes* | *polynesiensis* | 3.2 | 3.7 | 4.1 | 4.5 | 5.1 | 4.1 | 0.6 |
| *Anopheles* | *rivulorum* | 2.8 | 3.4 | 3.8 | 4.2 | 5.0 | 3.9 | 0.7 |
| *Anopheles* | *melas* | 2.8 | 3.3 | 3.7 | 4.1 | 4.8 | 3.8 | 0.7 |
| *Anopheles* | *culicifacies s.l.* | 2.8 | 3.3 | 3.7 | 4.1 | 4.7 | 3.7 | 0.6 |
| *Anopheles* | *subpictus s.l.* | 2.2 | 3.1 | 3.7 | 4.5 | 7.0 | 4.3 | 6.6 |
| *Aedes* | *samoanus* | 2.7 | 3.2 | 3.6 | 4.0 | 4.9 | 3.7 | 1.0 |
| *Anopheles* | *quadrimaculatus* | 2.4 | 3.1 | 3.5 | 4.0 | 4.8 | 3.6 | 0.8 |
| *Anopheles* | *stephensi* | 2.1 | 2.9 | 3.5 | 4.2 | 5.7 | 3.7 | 1.5 |
| *Anopheles* | *darlingi* | 2.3 | 2.9 | 3.4 | 3.8 | 4.7 | 3.4 | 0.9 |
| *Aedes* | *sollicitans* | 1.6 | 2.3 | 2.9 | 3.6 | 6.1 | 3.3 | 2.5 |
| *Aedes* | *vexans* | 1.7 | 2.2 | 2.7 | 3.1 | 4.2 | 3.0 | 3.3 |
| *Anopheles* | *cruzii* | 1.2 | 1.6 | 2.0 | 2.5 | 4.0 | 2.2 | 1.3 |
| *Anopheles* | *bellator* | 1.0 | 1.4 | 1.7 | 2.1 | 3.6 | 2.1 | 5.4 |
| *Culex* | | 2.5 | 4.0 | 5.4 | 7.6 | 12.3 | 6.2 | 3.2 |
| *Anopheles* | | 2.1 | 3.6 | 4.9 | 6.8 | 10.8 | 5.5 | 2.9 |
| *Mansonia* | | 3.3 | 4.0 | 4.5 | 4.9 | 5.6 | 4.5 | 0.8 |
| *Aedes* | | 2.5 | 3.3 | 3.8 | 4.3 | 5.2 | 3.8 | 0.9 |
| *Overall* | | 2.0 | 3.3 | 4.7 | 6.4 | 10.2 | 5.2 | 2.6 |

Table S6: **Polovodova dissection: mean female mosquito lifespan estimates in terms of chronological time.** The 5%, 25%, 50%, 75% and 95% columns indicate the respective quantiles of the posterior distribution for mean lifespan, and the 'Mean' and 'Std. dev.' columns indicate the posterior mean and standard deviation of mean lifespan. Units of lifespan are days.

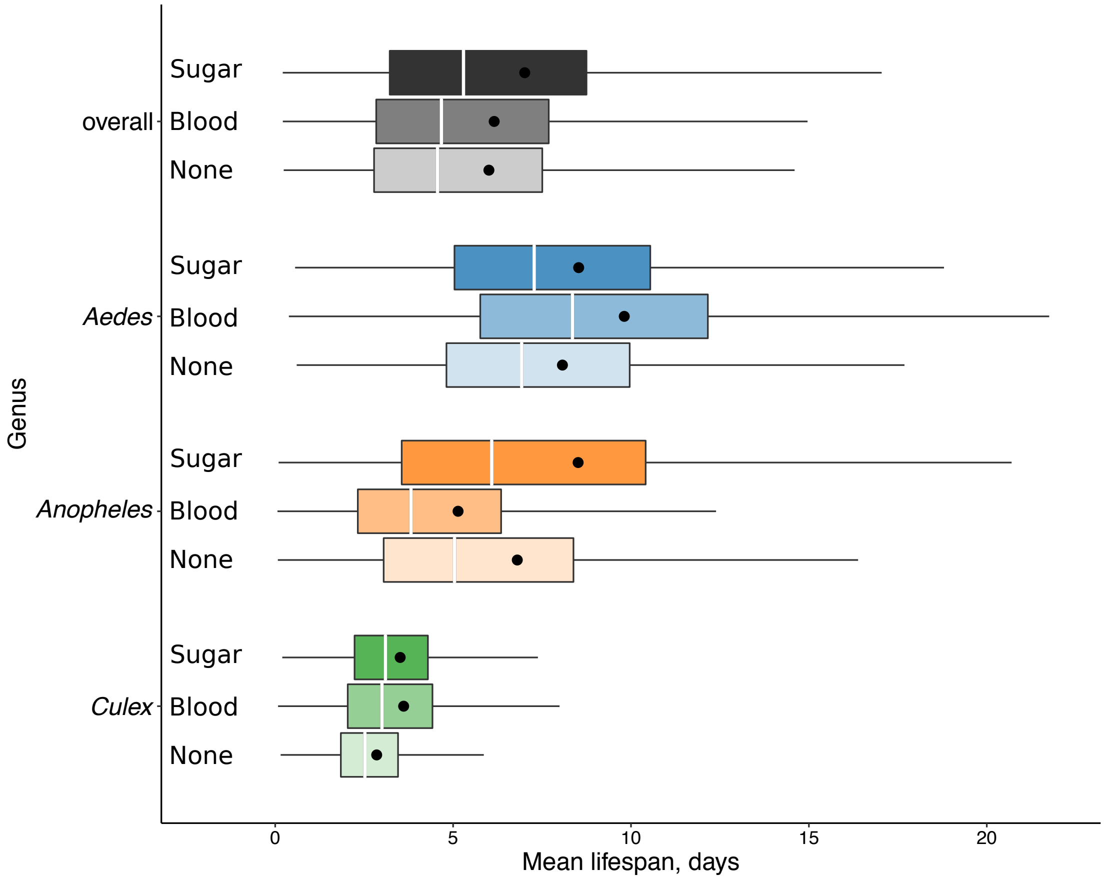


Figure S1: **MRR: female mosquito mean lifespan by pre-release feeding status.** Labels to the left of the vertical axis indicate genus or overall grouping; the labels to the right of the axis indicate pre-release feeding status. The middle line in each box shows the median estimates and the solid point indicates the mean. The left and right box edges show the 25%, and 75% posterior quantiles respectively. The whiskers show the range of the data, excluding points lying more than 1.5 times the interquartile range away from each edge of the box. All estimates were obtained using the hierarchical exponential survival model.

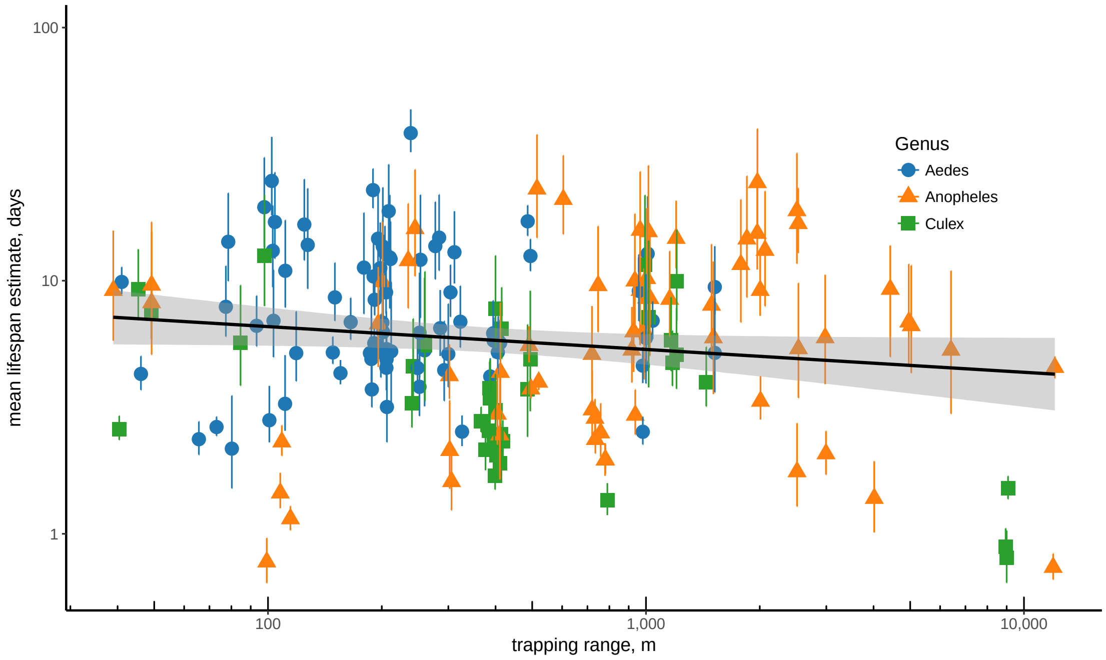


Figure S2: **MRR: mean mosquito lifespan versus trapping range.** The markers show the median posterior lifespan estimates for each time series with the lower and upper bounds indicating the 25% and 75% quantiles, respectively. The black line shows a linear regression line estimated using the median posterior lifetimes, with the grey shading indicating 95% confidence intervals. All estimates were obtained using the non-hierarchical exponential survival model.

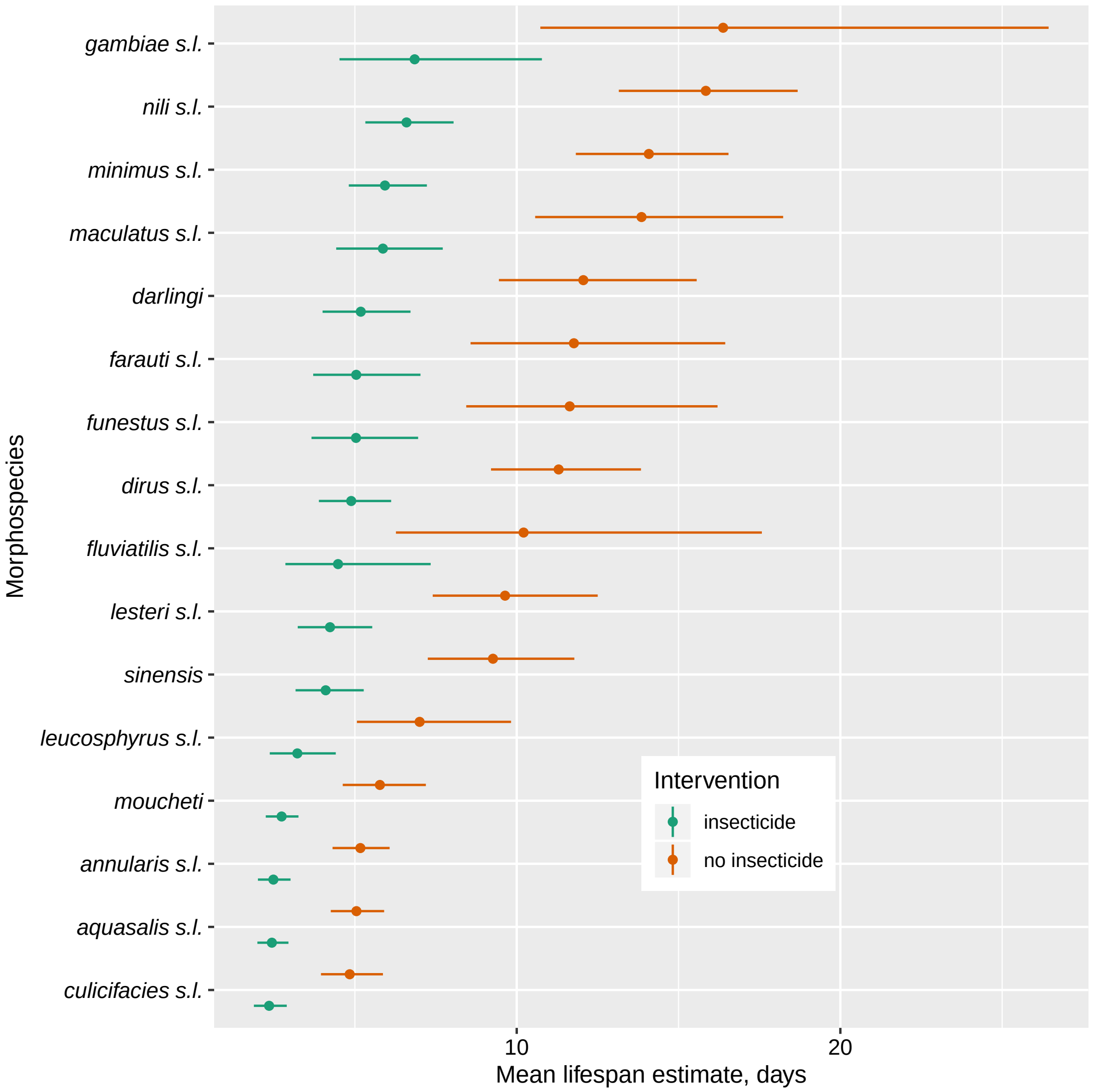


Figure S3: **Detinova dissection: estimated impact of insecticide on lifespan.** The middle line in each box shows the median estimates. The left and right box whiskers show the 10%, and 90% posterior quantiles respectively.

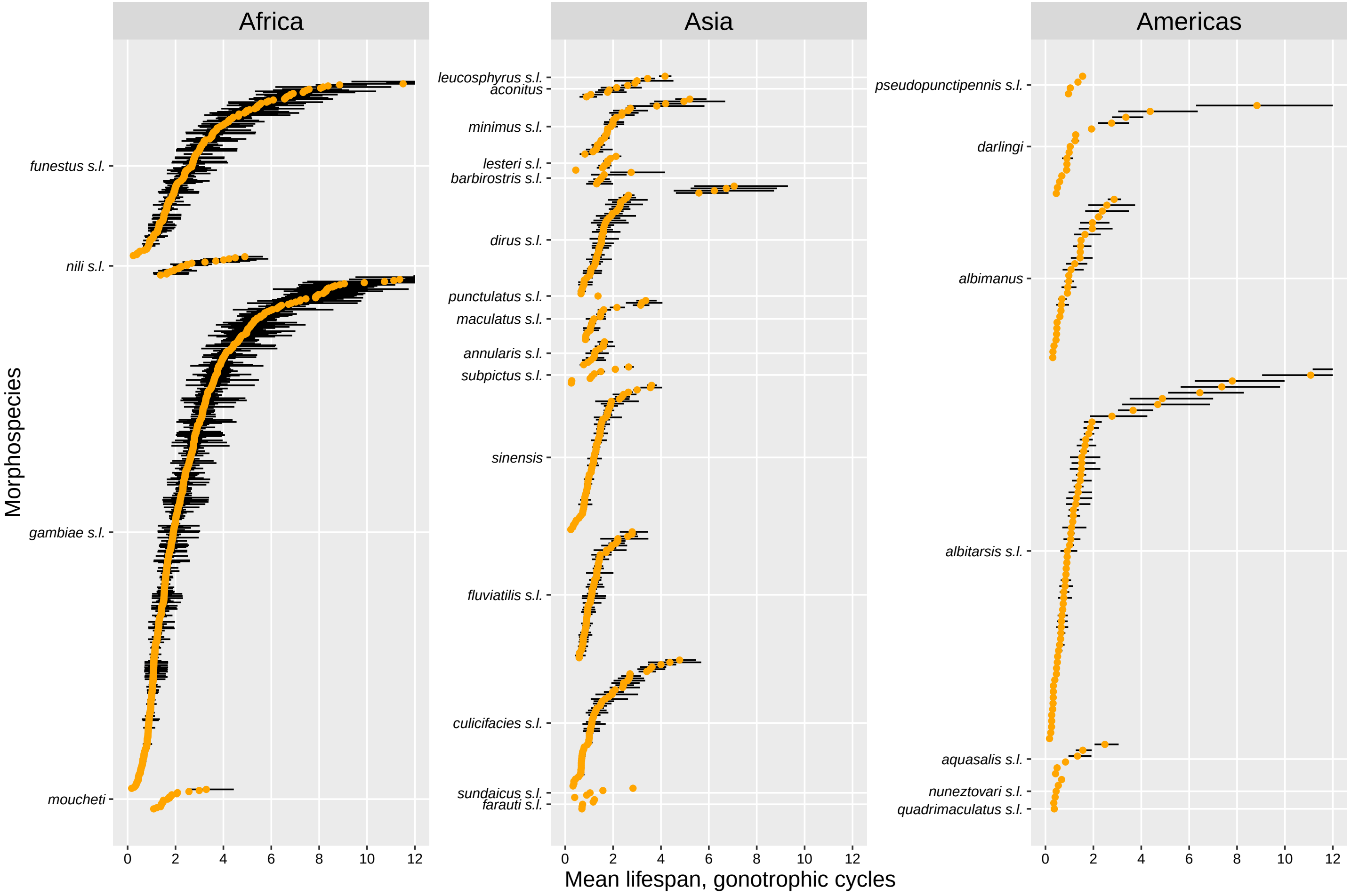


Figure S4: **Detinova dissection: individual time-series estimates of adult mosquito mean lifespan.** The middle line in each box shows the median estimates. The left and right box whiskers show the 25%, and 75% posterior quantiles respectively. Note, that a handful of observations are not shown in the plot since their estimates exceeded the reasonable range (here deemed to be up to 12 cycles).

### S2 Evidence for senescence

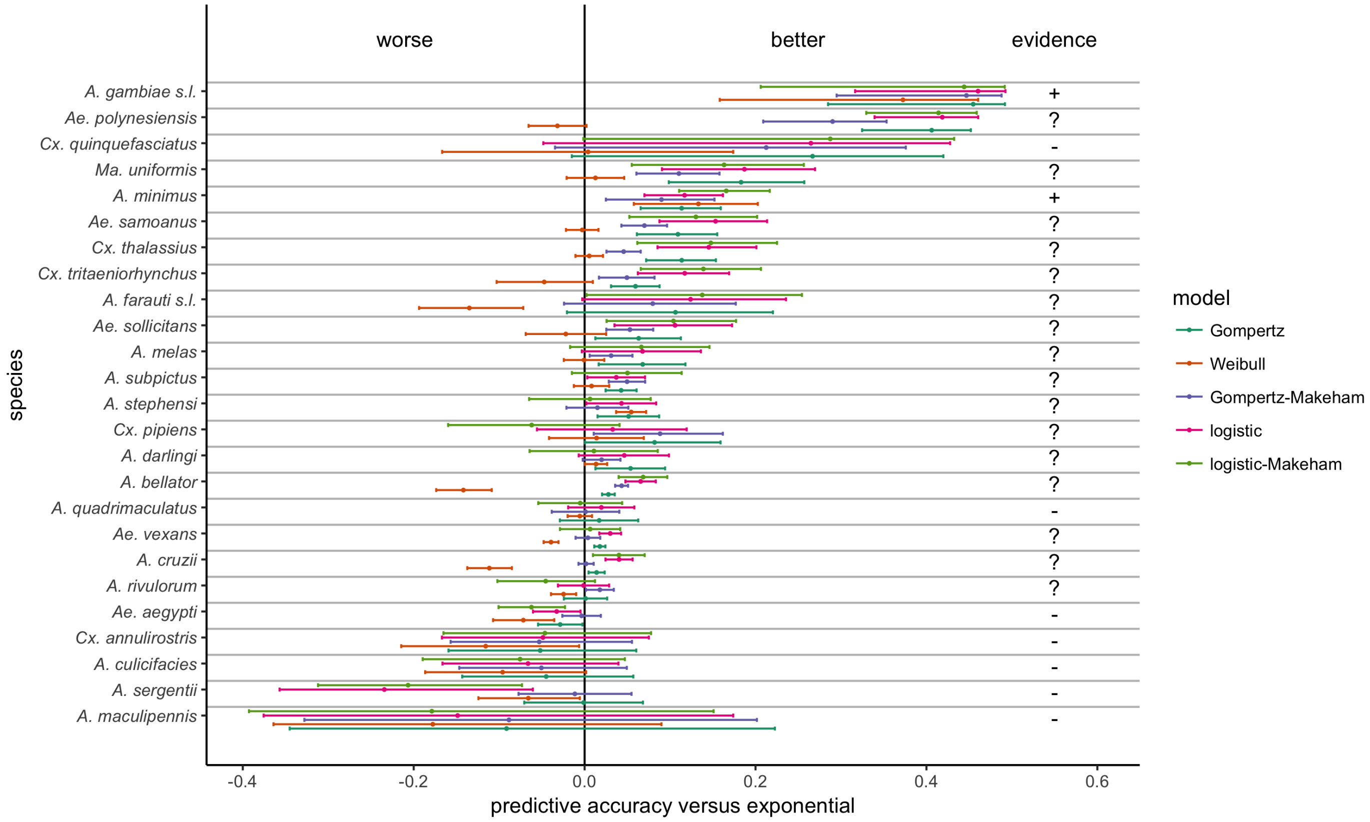


Figure S5: **Polovodova dissection: evidence for senescence.** Here, we consider the predictive accuracy of age-dependent models of mortality versus the exponential model by species. The predictive accuracy was determined using K-Fold cross-validation as described in SOM; the measure of accuracy presented here by the central dots is the difference in estimated expected log pointwise predictive density compared to the exponential model. The lower and upper whiskers represent the lower and upper bounds of the 95% confidence interval in predictive accuracy. The species have been ordered so that those with the highest average predictive accuracy across all age-dependent models compared to the exponential model appear at the top. If all age-dependent models outperform the exponential, we deem this as evidence for age-dependent mortality ('+'); if a subsample of models perform better, we deem the evidence ambiguous ('?'); and if no age-dependent model exceeds the performance of the exponential, we conclude no evidence in favour of senescence ('-').

### S3 Power analysis of MRR studies

Consider an ideal MRR experiment, where mosquitoes do not migrate out of the study area and marking is harmless. With these data, how accurately can we estimate mosquito lifespan? To address this question, we generated artificial MRR data and attempted to estimate the (known) parameters by maximum likelihood (a 'Monte Carlo' analysis). We simulated releases of $N$ mosquitoes that are monitored for $m$ days in each case (with collections occurring on each day) and determined how the errors in estimating lifespan depended on these two parameters. In order to focus on estimating lifespan, we assumed the mosquitoes experience constant mortality, and recaptures follow the negative binomial sampling model where the (known) recapture probability parameters $\psi = 0.03$ and $\kappa = 3.18$ were the averages of the posterior draws across all species in the species-level model. We assume that all parameters: the true lifespan, $\psi$ and $\kappa$ were unknown and were estimated via maximum likelihood. Here, we investigated two lifespans: 5 days and 10 days indicative of estimates obtained in our analyses. For each simulation, we determined the Root Mean Square Error (RMSE) in predicting mean lifespan.

Unsurprisingly, the error in predicting lifespan declines as the study duration increases (Fig. S6A). However once a study length is much longer than the lifespan of the majority of mosquitoes, the predictive power cannot be improved by extending the study duration. The effect of increasing release size (while holding study length constant) similarly exhibits diminishing returns (Fig. S6B). For the parameters we use, there are significant gains in accuracy from releasing 1000 rather than 100 mosquitoes, but modest gains from releasing $10,000$ rather than 1000 mosquitoes.

We also conducted a power analysis for the detection of senescence in MRR experiments (Fig. S7). Here we calculated the power to detect senescence assuming that lifespan follows a Gompertz survival function (see Table SM3). We simulated MRR time series for single releases, for case study populations with three different levels of senescence assuming a Gompertz survival function (Fig. S7A.). We assumed that recaptures followed a negative binomial sampling model with $\psi = 0.03$ and $\kappa = 3.18$. Using each simulated time series of recaptures, we then performed two different maximum likelihood estimations: one, where we (falsely) assumed that lifespan followed an exponential distribution (i.e. there was no senescence); and another where we assumed it followed a Gompertz survival function. In both cases, we estimated the parameters of the corresponding survival function along with $\psi$ and $\kappa$. We then performed a likelihood ratio test (using a 5% test size) comparing the fit of the Gompertz model to that of the simpler exponential model: in doing so, we determined whether there was evidence of senescence. We performed 500 experimental replicates at each parameter set.

These indicate that the power to detect senescence is strongly dependent on study length (Fig. S7B.) but is insensitive to release size (Fig. S7C.). Clements & Paterson (1981) conducted a meta-analysis of MRR and dissection-based experiments and found evidence of senescence that is, at least, qualitatively similar in magnitude to that of the 'mild' case considered above. For this case, detecting senescence with a power of 80% requires a study length of at least 18 days. Since the median study length for experiments included in our analysis was 10 days, (Table SM1) this could partly explain our failure to detect senescence at the species level.

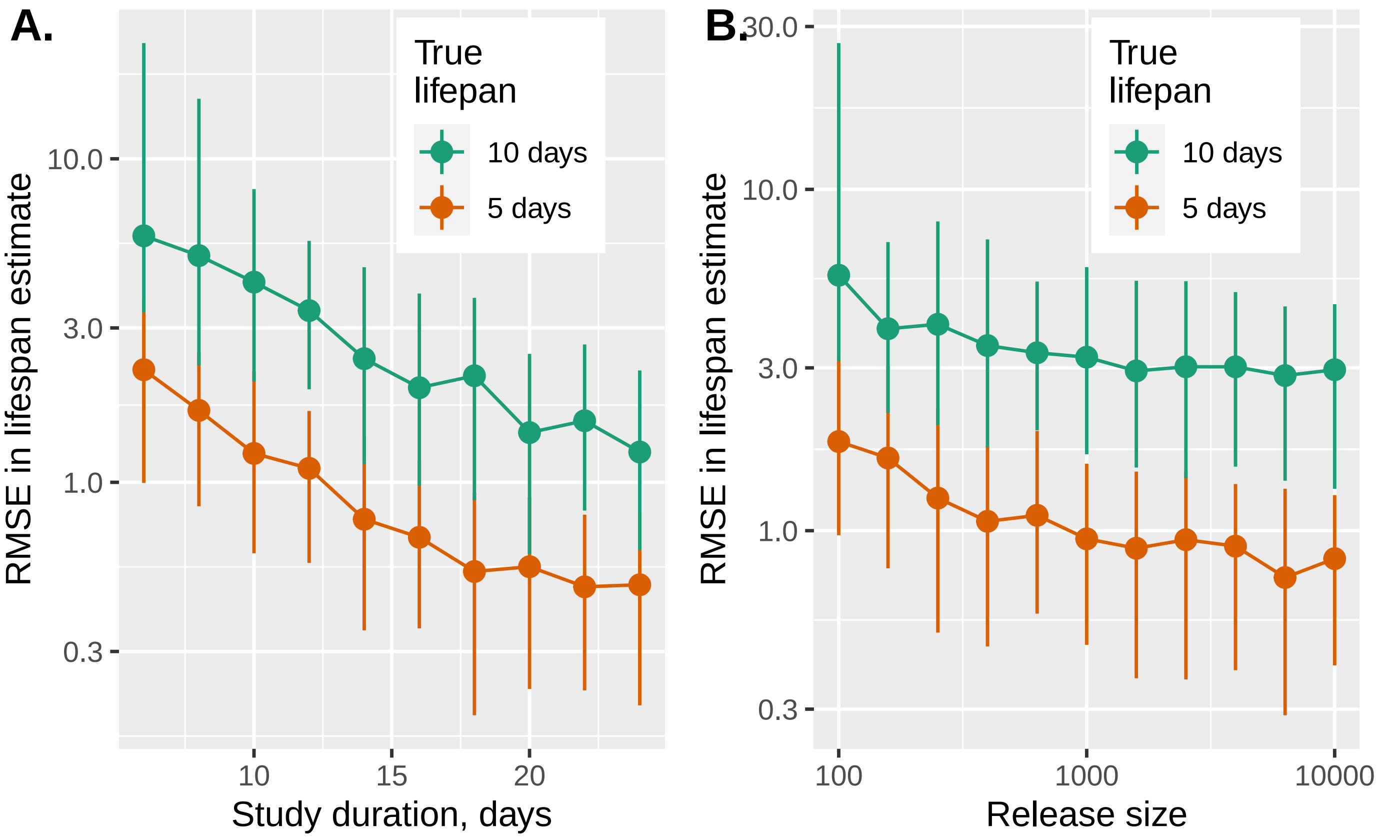


Figure S6: **MRR: Monte Carlo analysis of errors in predicting mean lifespan.** This shows the Root Mean Square Error (RMSE) in predicting mean lifespan as a function of study length (A) and number of released mosquitoes (B) Monte Carlo analysis. For each parameter set, we generated 200 simulated data series using the negative binomial sampling model with an exponential survival function as described in the text and estimated the mean lifespan using maximum likelihood. The error bars show the 25%-75% quantiles, and the points show the medians.

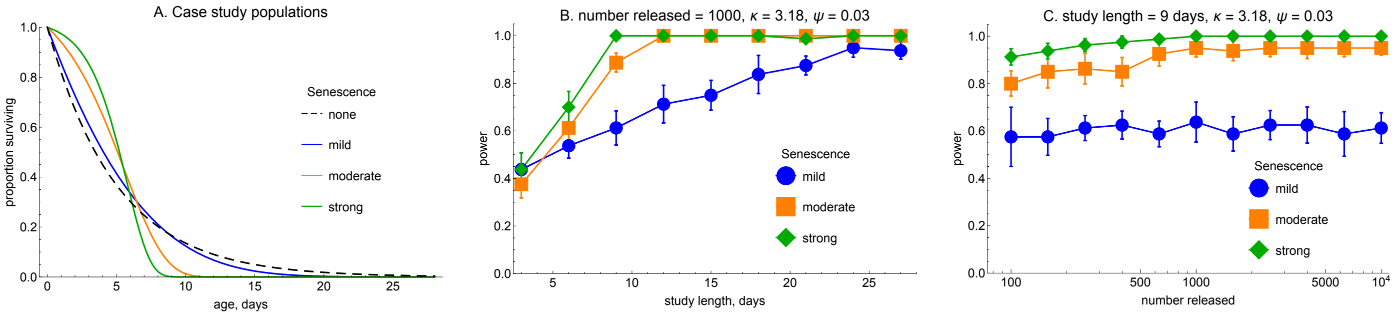


Figure S7: **MRR: statistical power analysis for senescence detection.** The plot shows the statistical power to detect senescence for A. three case study populations as a function of B. study length and C. number released. For each parameter set we generated 500 simulated data series using the negative binomial sampling model with a gompertz hazard function ($\lambda(t) = \alpha e^{\beta t}$), and estimated the $\beta$ parameter using maximum likelihood and tested the null hypothesis $\beta = 0$ (constant mortality risk) against the alternative $\beta > 0$ using an likelihood ratio test with a 5% test size. The error bars show the standard deviation in the power for each parameter set using groups of 40 simulated time series as bootstrap samples.

### References

Clements, AN, & Paterson, GD. 1981. The analysis of mortality and survival rates in wild populations of mosquitoes. *Journal of Applied Ecology*, 373–399.

Vehtari, Aki, Gelman, Andrew, & Gabry, Jonah. 2015. Efficient implementation of leave-one-out cross-validation and WAIC for evaluating fitted Bayesian models. *arXiv preprint arXiv:1507.04544*.
