## Supplementary material for "A Meta-analysis of Longevity Estimates of Mosquito Vectors of Disease": Posterior predictive checks for MRR series

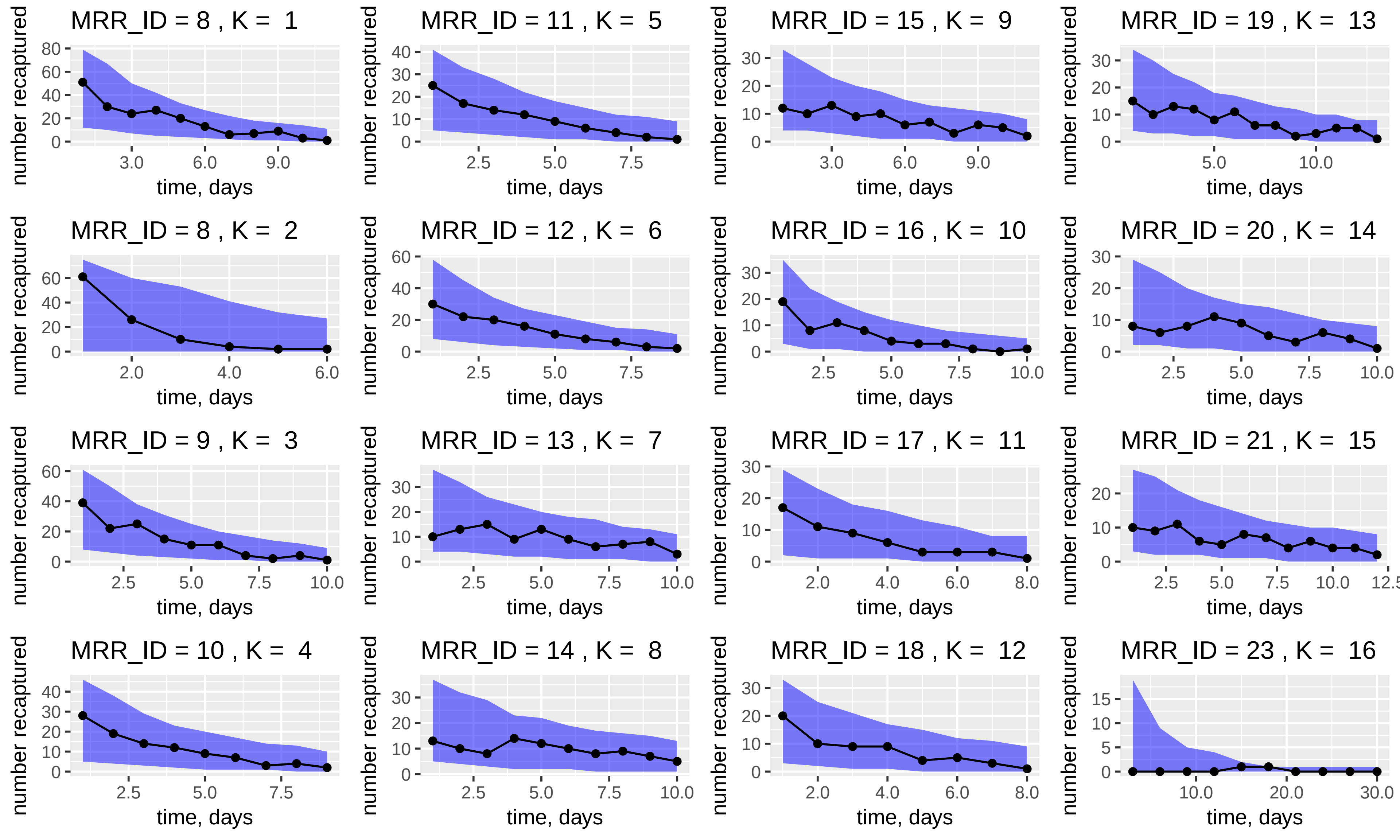
MRR_ID = 8 , K = 1
MRR_ID = 11 , K = 5
MRR_ID = 15 , K = 9
MRR_ID = 19 , K = 13
MRR_ID = 8 , K = 2
MRR_ID = 12 , K = 6
MRR_ID = 16 , K = 10
MRR_ID = 20 , K = 14
MRR_ID = 9 , K = 3
MRR_ID = 13 , K = 7
MRR_ID = 17 , K = 11
MRR_ID = 21 , K = 15
MRR_ID = 10 , K = 4
MRR_ID = 14 , K = 8
MRR_ID = 18 , K = 12
MRR_ID = 23 , K = 16
number recaptured
time, days

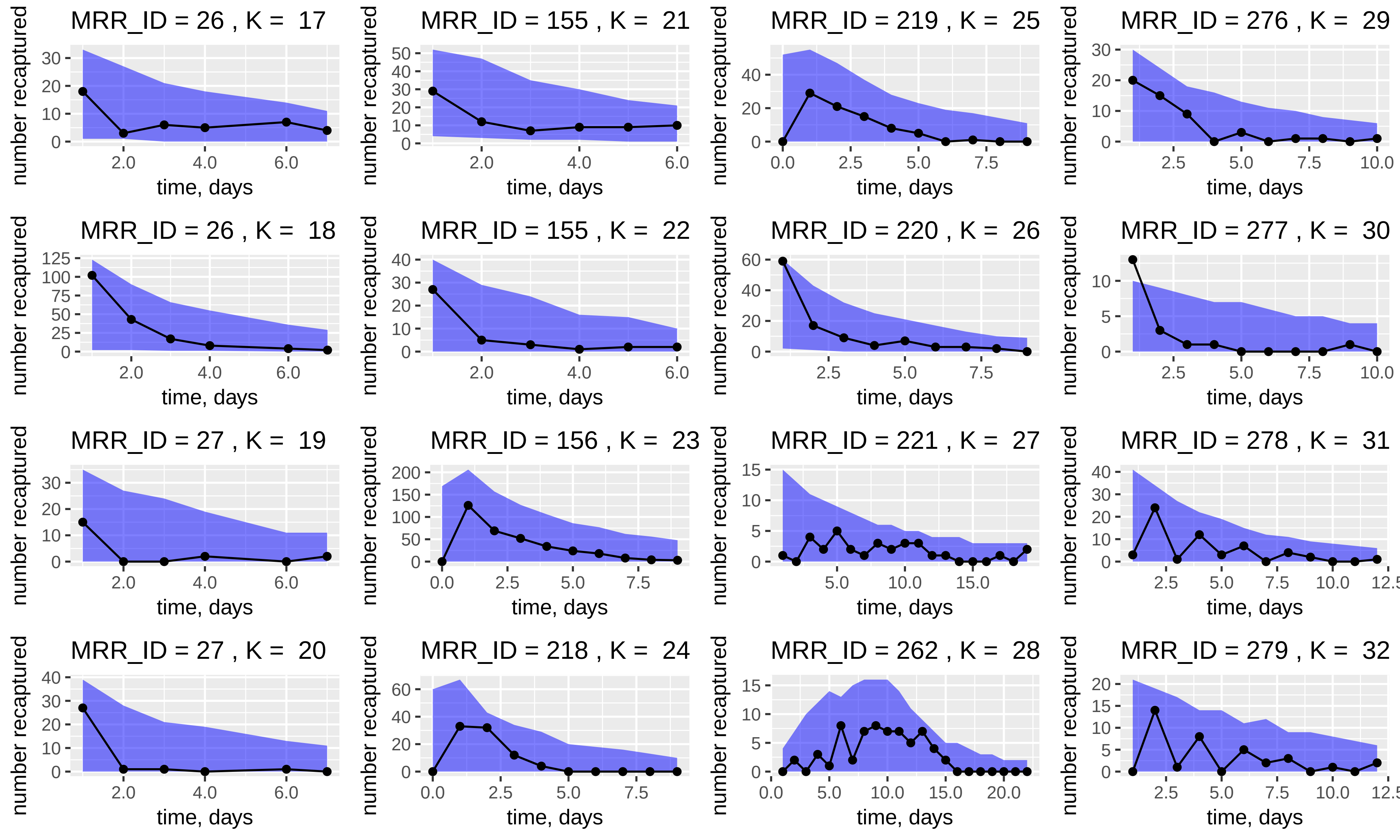
MRR_ID = 26 , K = 17
MRR_ID = 155 , K = 21
MRR_ID = 219 , K = 25
MRR_ID = 276 , K = 29
MRR_ID = 26 , K = 18
MRR_ID = 155 , K = 22
MRR_ID = 220 , K = 26
MRR_ID = 277 , K = 30
MRR_ID = 27 , K = 19
MRR_ID = 156 , K = 23
MRR_ID = 221 , K = 27
MRR_ID = 278 , K = 31
MRR_ID = 27 , K = 20
MRR_ID = 218 , K = 24
MRR_ID = 262 , K = 28
MRR_ID = 279 , K = 32
number recaptured
time, days

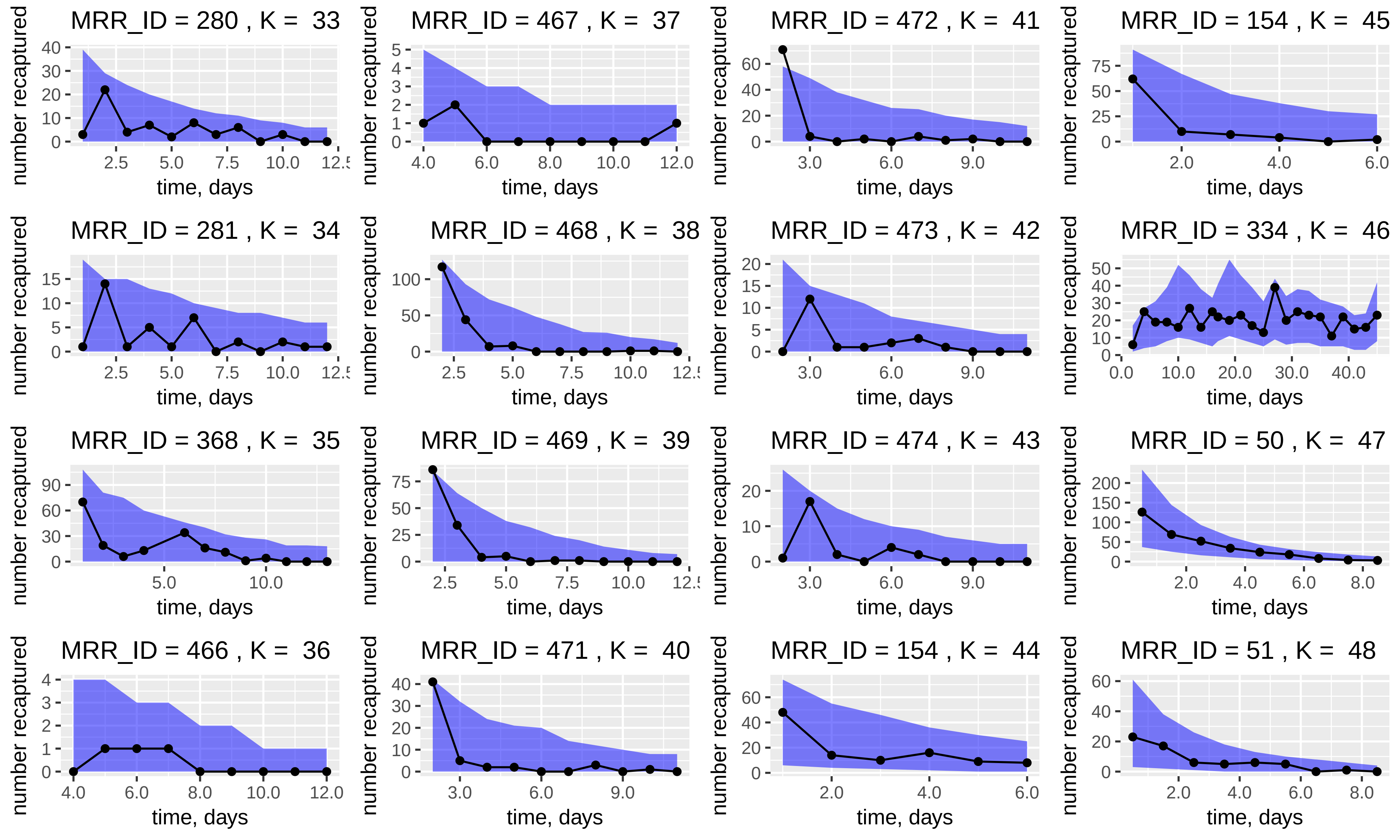

MRR_ID = 280 , K = 33
MRR_ID = 467 , K = 37
MRR_ID = 472 , K = 41
MRR_ID = 154 , K = 45
MRR_ID = 281 , K = 34
MRR_ID = 468 , K = 38
MRR_ID = 473 , K = 42
MRR_ID = 334 , K = 46
MRR_ID = 368 , K = 35
MRR_ID = 469 , K = 39
MRR_ID = 474 , K = 43
MRR_ID = 50 , K = 47
MRR_ID = 466 , K = 36
MRR_ID = 471 , K = 40
MRR_ID = 154 , K = 44
MRR_ID = 51 , K = 48
number recaptured
time, days

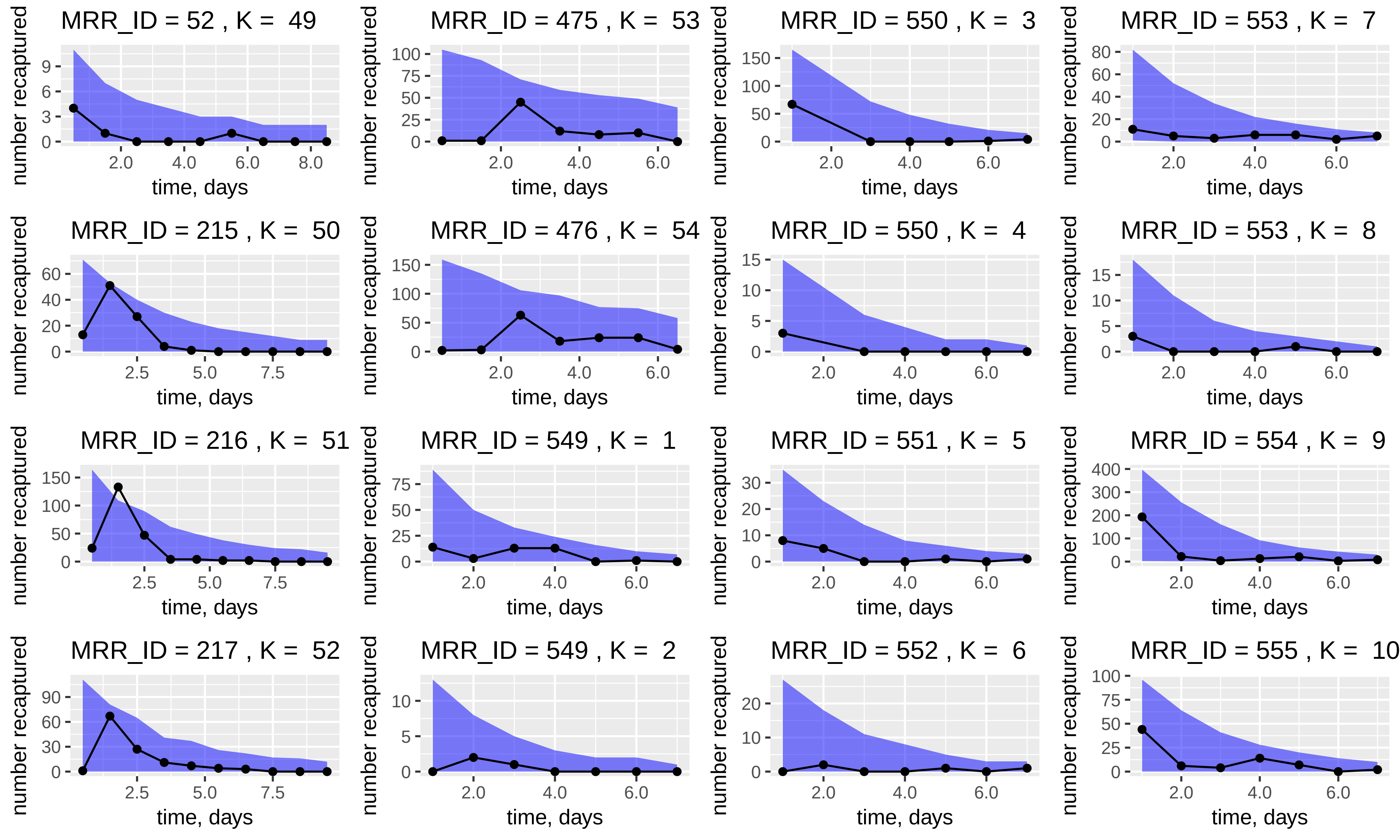

MRR_ID = 52 , K = 49
MRR_ID = 475 , K = 53
MRR_ID = 550 , K = 3
MRR_ID = 553 , K = 7
MRR_ID = 215 , K = 50
MRR_ID = 476 , K = 54
MRR_ID = 550 , K = 4
MRR_ID = 553 , K = 8
MRR_ID = 216 , K = 51
MRR_ID = 549 , K = 1
MRR_ID = 551 , K = 5
MRR_ID = 554 , K = 9
MRR_ID = 217 , K = 52
MRR_ID = 549 , K = 2
MRR_ID = 552 , K = 6
MRR_ID = 555 , K = 10
number recaptured
time, days

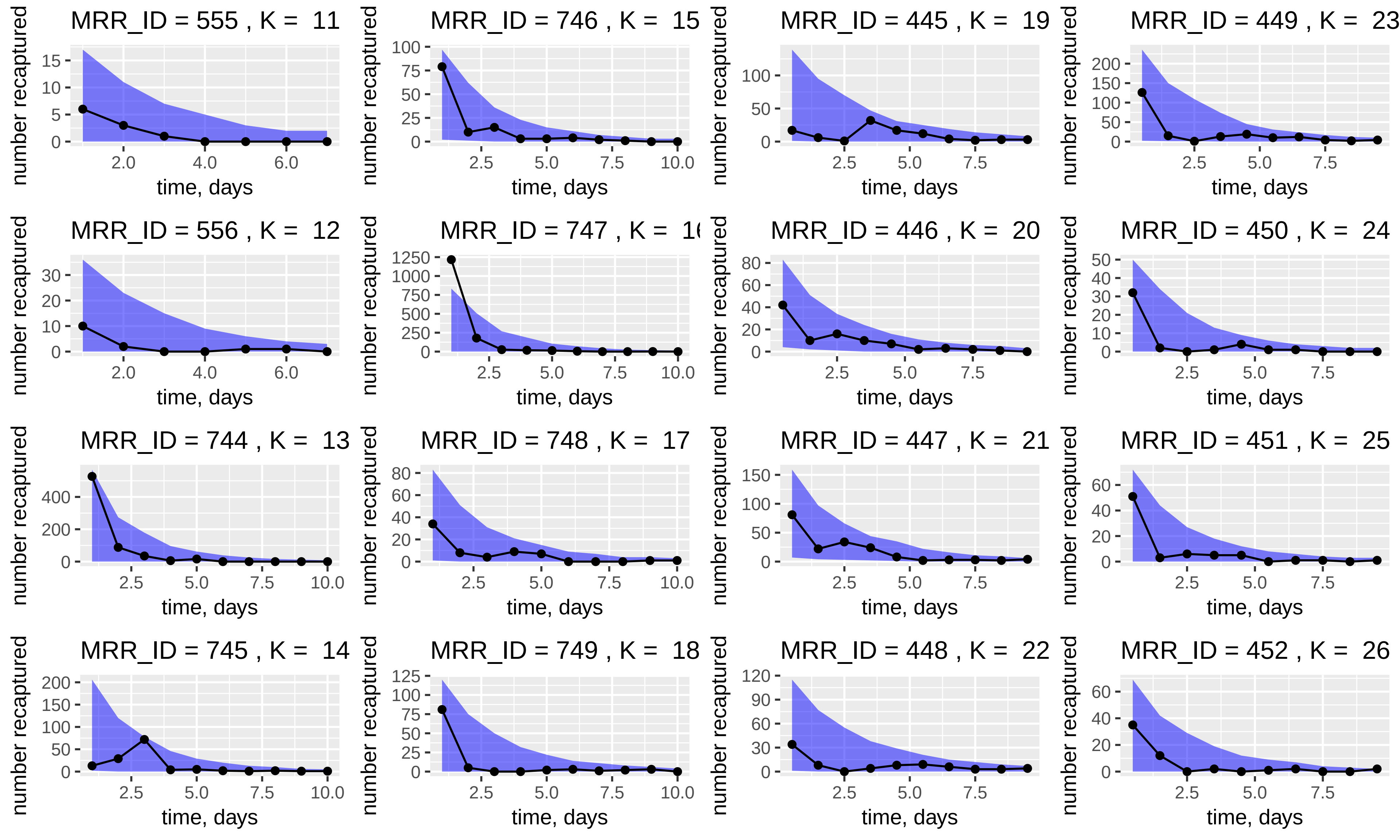
MRR_ID = 555 , K = 11
MRR_ID = 746 , K = 15
MRR_ID = 445 , K = 19
MRR_ID = 449 , K = 23
MRR_ID = 556 , K = 12
MRR_ID = 747 , K = 1
MRR_ID = 446 , K = 20
MRR_ID = 450 , K = 24
MRR_ID = 744 , K = 13
MRR_ID = 748 , K = 17
MRR_ID = 447 , K = 21
MRR_ID = 451 , K = 25
MRR_ID = 745 , K = 14
MRR_ID = 749 , K = 18
MRR_ID = 448 , K = 22
MRR_ID = 452 , K = 26
number recaptured
time, days

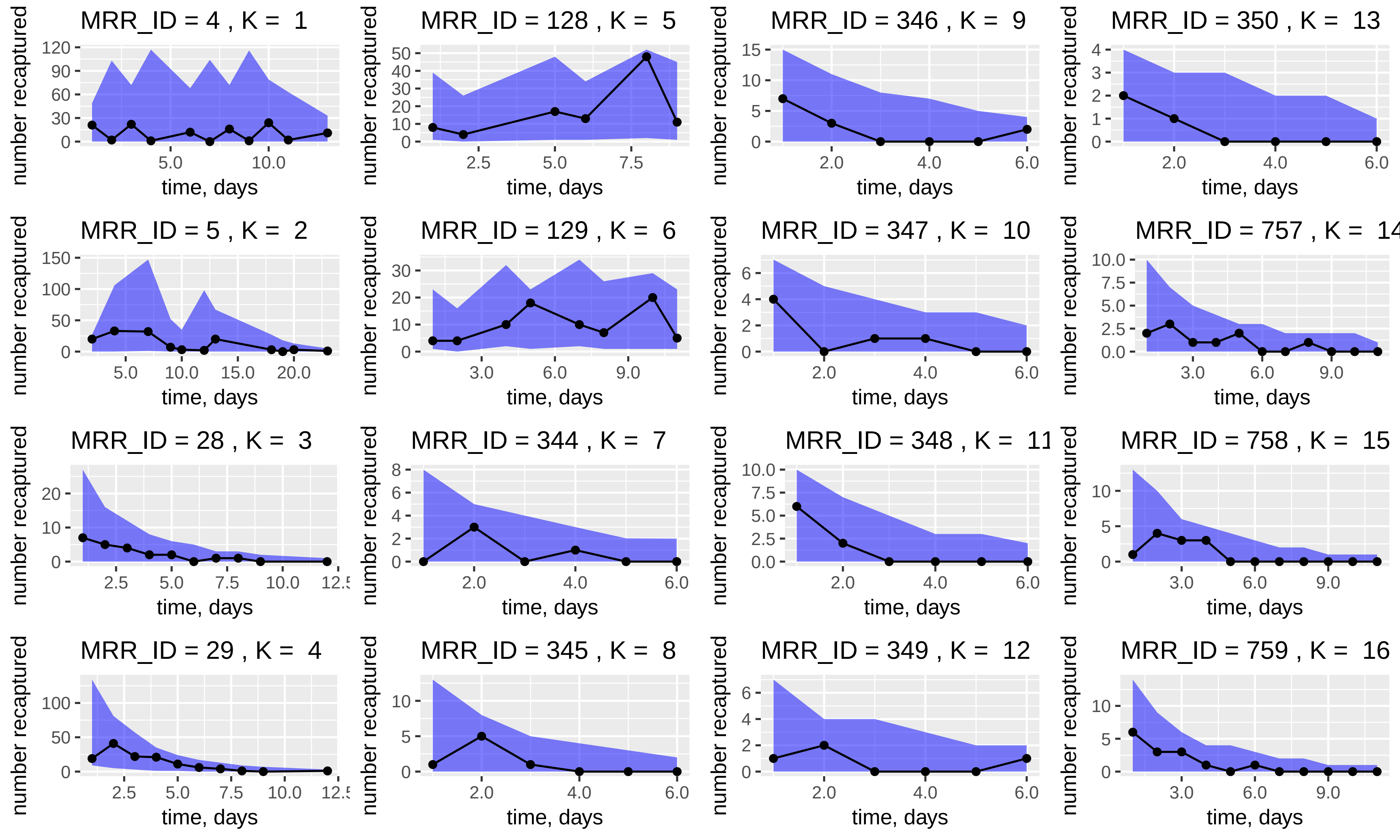

MRR_ID = 4 , K = 1
MRR_ID = 128 , K = 5
MRR_ID = 346 , K = 9
MRR_ID = 350 , K = 13
MRR_ID = 5 , K = 2
MRR_ID = 129 , K = 6
MRR_ID = 347 , K = 10
MRR_ID = 757 , K =
MRR_ID = 28 , K = 3
MRR_ID = 344 , K = 7
MRR_ID = 348 , K = 11
MRR_ID = 758 , K = 15
MRR_ID = 29 , K = 4
MRR_ID = 345 , K = 8
MRR_ID = 349 , K = 12
MRR_ID = 759 , K = 16
number recaptured
time, days

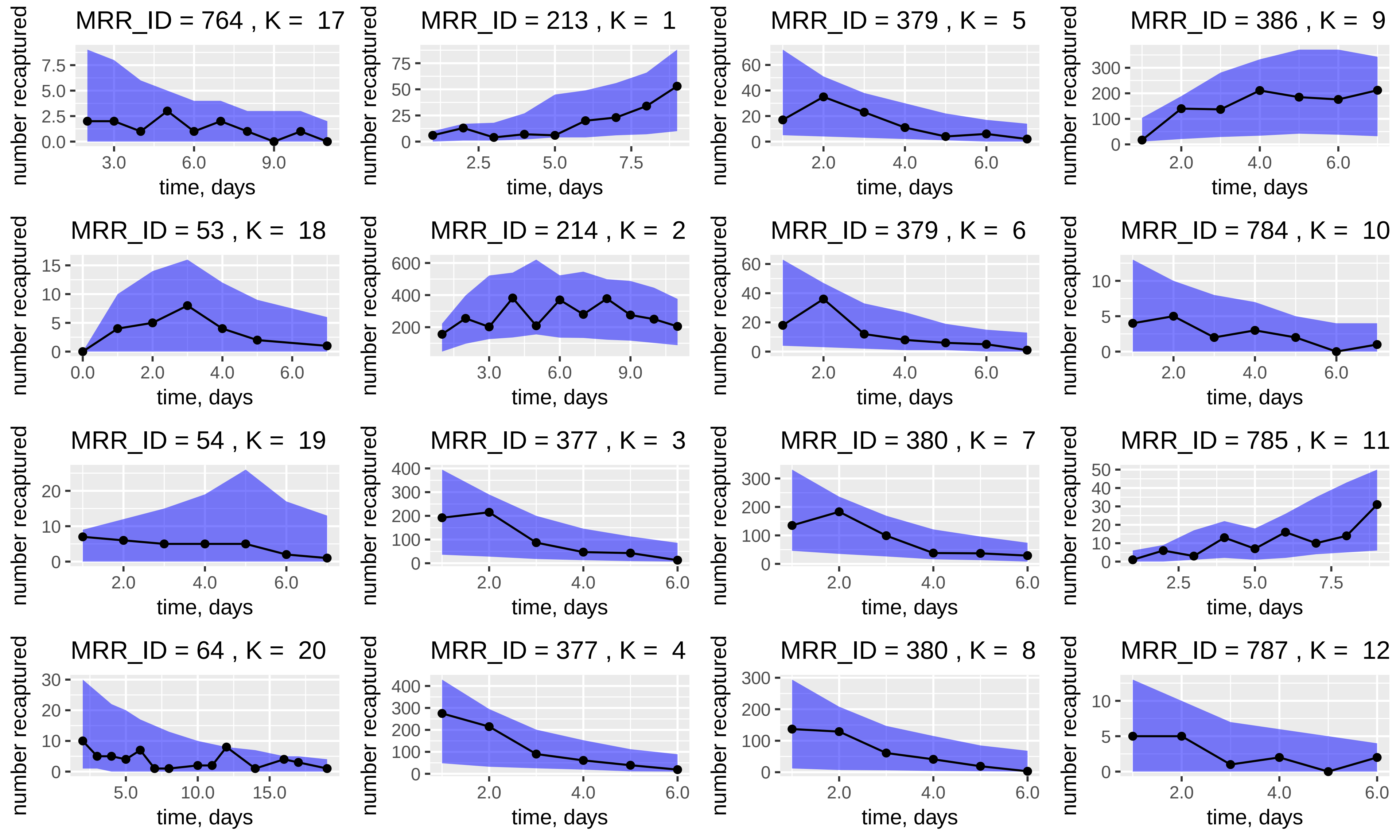
MRR_ID = 764 , K = 17
MRR_ID = 213 , K = 1
MRR_ID = 379 , K = 5
MRR_ID = 386 , K = 9
MRR_ID = 53 , K = 18
MRR_ID = 214 , K = 2
MRR_ID = 379 , K = 6
MRR_ID = 784 , K = 10
MRR_ID = 54 , K = 19
MRR_ID = 377 , K = 3
MRR_ID = 380 , K = 7
MRR_ID = 785 , K = 11
MRR_ID = 64 , K = 20
MRR_ID = 377 , K = 4
MRR_ID = 380 , K = 8
MRR_ID = 787 , K = 12
number recaptured
time, days

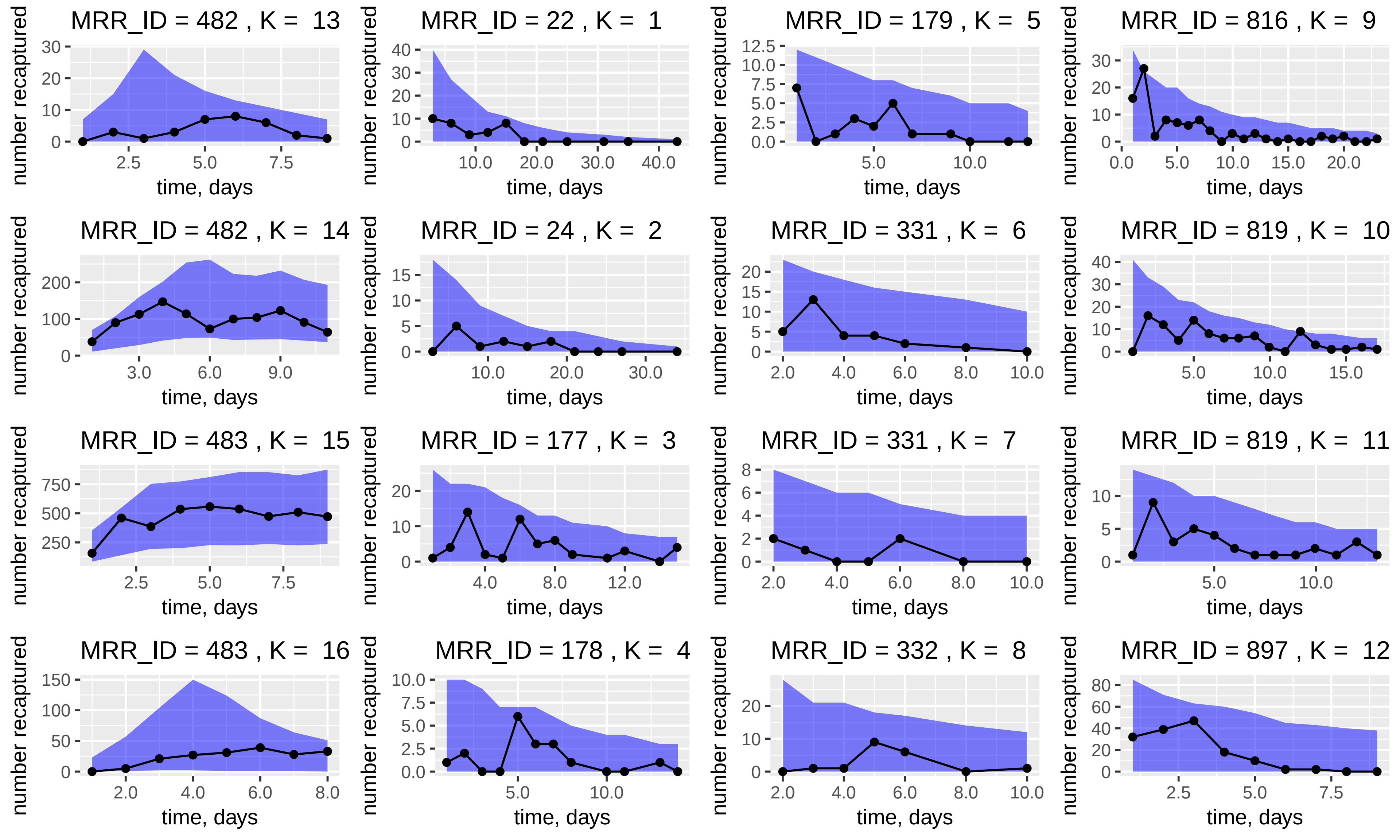
MRR_ID = 482 , K = 13
MRR_ID = 22 , K = 1
MRR_ID = 179 , K = 5
MRR_ID = 816 , K = 9
MRR_ID = 482 , K = 14
MRR_ID = 24 , K = 2
MRR_ID = 331 , K = 6
MRR_ID = 819 , K = 10
MRR_ID = 483 , K = 15
MRR_ID = 177 , K = 3
MRR_ID = 331 , K = 7
MRR_ID = 819 , K = 11
MRR_ID = 483 , K = 16
MRR_ID = 178 , K = 4
MRR_ID = 332 , K = 8
MRR_ID = 897 , K = 12
number recaptured
time, days

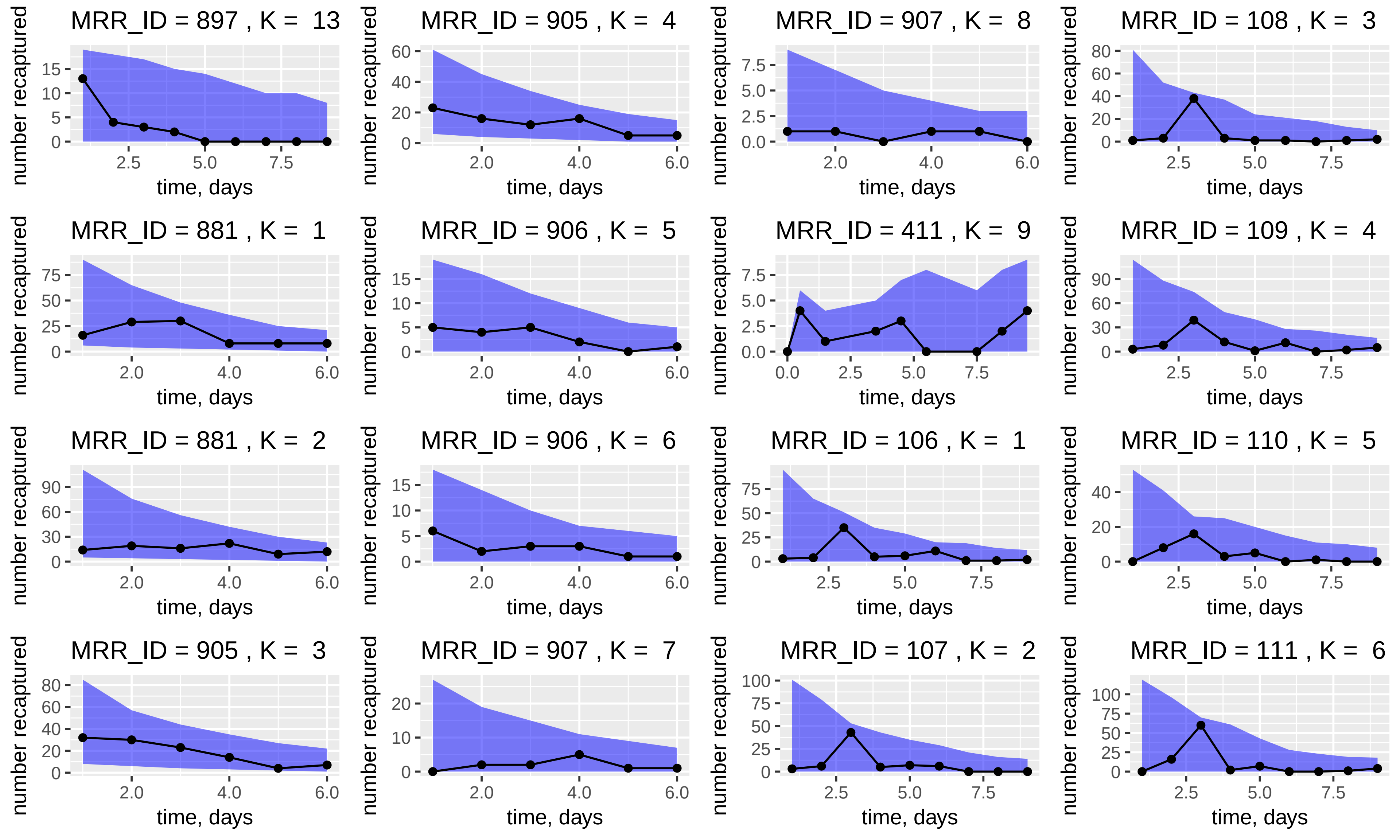
MRR_ID = 897 , K = 13
MRR_ID = 905 , K = 4
MRR_ID = 907 , K = 8
MRR_ID = 108 , K = 3
MRR_ID = 881 , K = 1
MRR_ID = 906 , K = 5
MRR_ID = 411 , K = 9
MRR_ID = 109 , K = 4
MRR_ID = 881 , K = 2
MRR_ID = 906 , K = 6
MRR_ID = 106 , K = 1
MRR_ID = 110 , K = 5
MRR_ID = 905 , K = 3
MRR_ID = 907 , K = 7
MRR_ID = 107 , K = 2
MRR_ID = 111 , K = 6
number recaptured
time, days

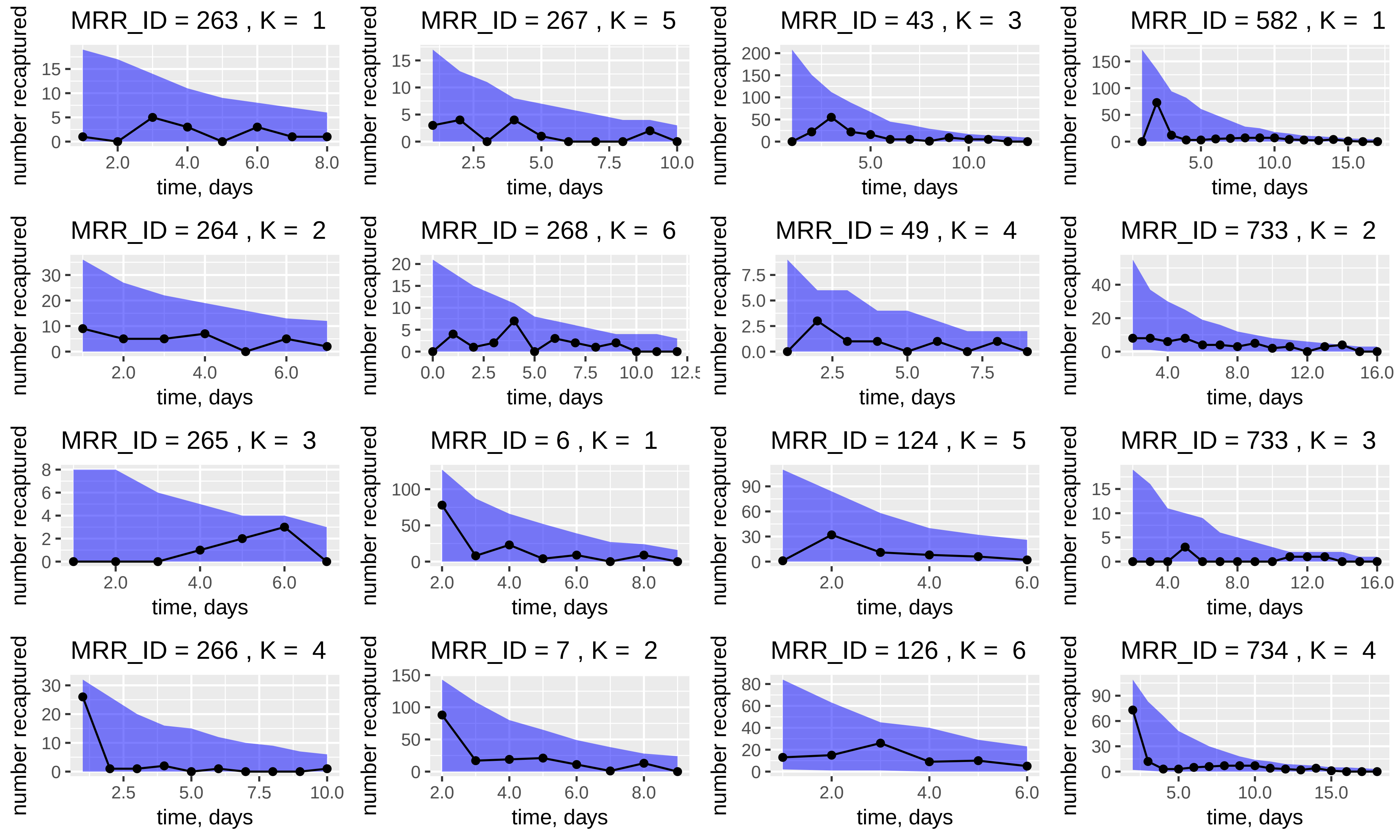

MRR_ID = 263 , K = 1
MRR_ID = 267 , K = 5
MRR_ID = 43 , K = 3
MRR_ID = 582 , K = 1
MRR_ID = 264 , K = 2
MRR_ID = 268 , K = 6
MRR_ID = 49 , K = 4
MRR_ID = 733 , K = 2
MRR_ID = 265 , K = 3
MRR_ID = 6 , K = 1
MRR_ID = 124 , K = 5
MRR_ID = 733 , K = 3
MRR_ID = 266 , K = 4
MRR_ID = 7 , K = 2
MRR_ID = 126 , K = 6
MRR_ID = 734 , K = 4
number recaptured
time, days

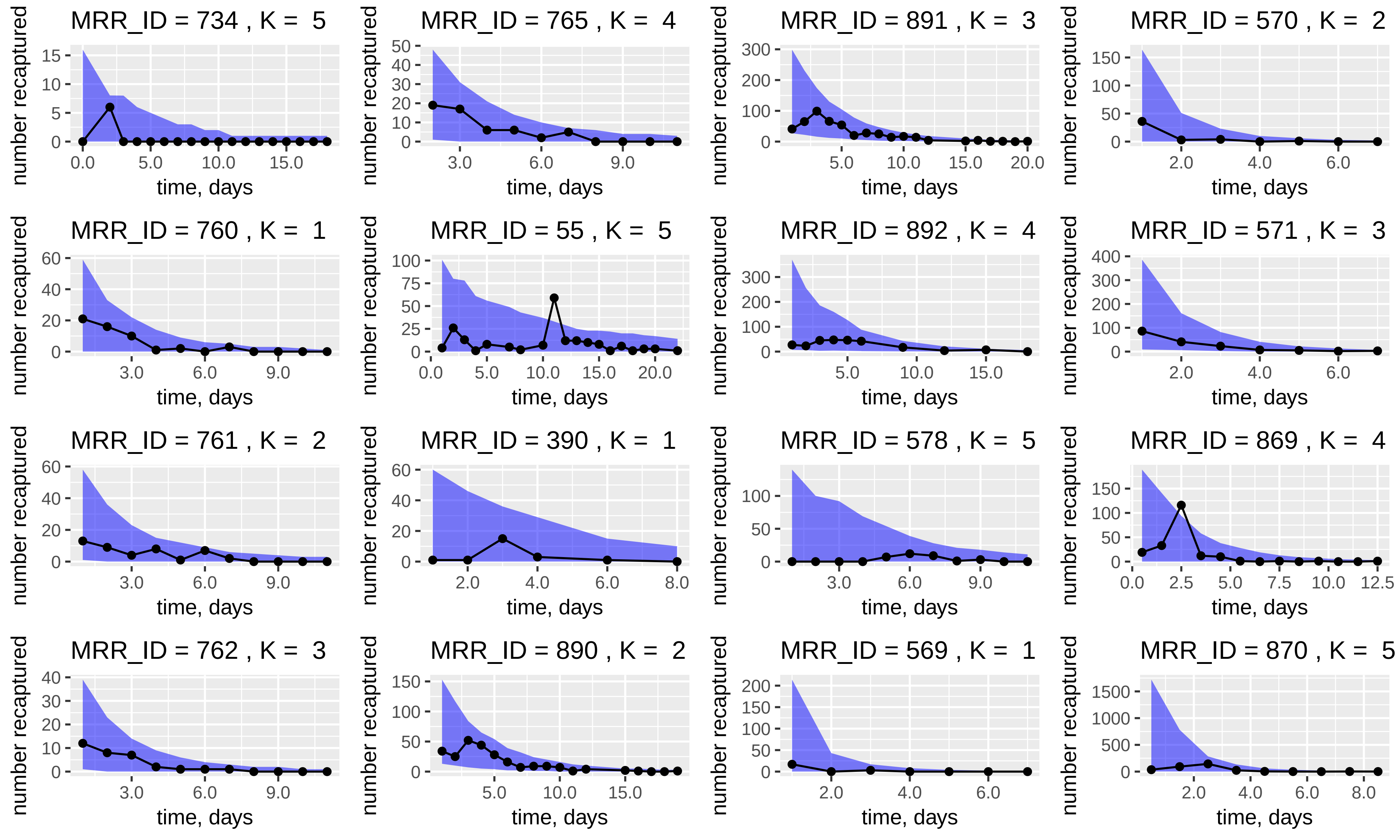
MRR_ID = 734 , K = 5
MRR_ID = 765 , K = 4
MRR_ID = 891 , K = 3
MRR_ID = 570 , K = 2
MRR_ID = 760 , K = 1
MRR_ID = 55 , K = 5
MRR_ID = 892 , K = 4
MRR_ID = 571 , K = 3
MRR_ID = 761 , K = 2
MRR_ID = 390 , K = 1
MRR_ID = 578 , K = 5
MRR_ID = 869 , K = 4
MRR_ID = 762 , K = 3
MRR_ID = 890 , K = 2
MRR_ID = 569 , K = 1
MRR_ID = 870 , K = 5
number recaptured
time, days

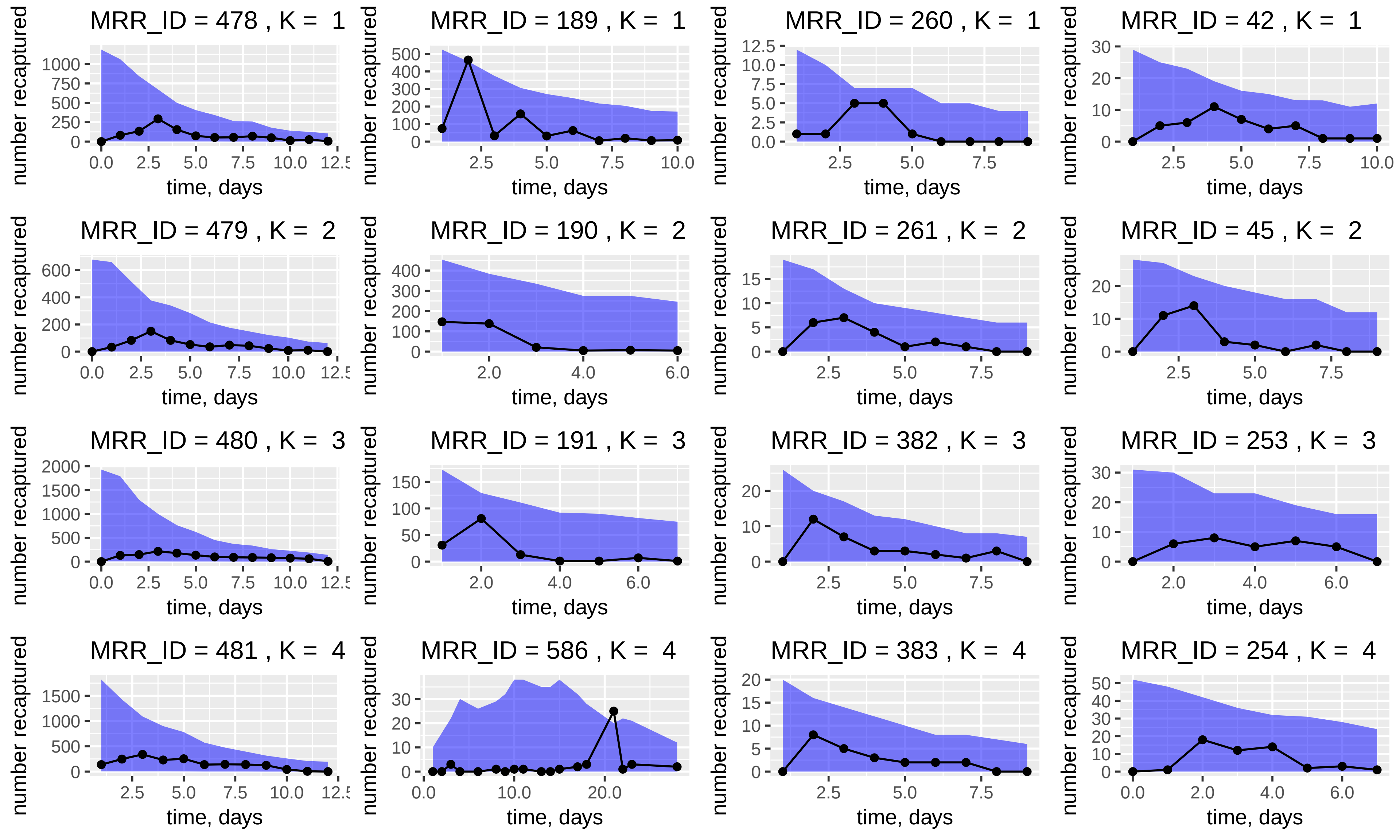
MRR_ID = 478 , K = 1
MRR_ID = 189 , K = 1
MRR_ID = 260 , K = 1
MRR_ID = 42 , K = 1
MRR_ID = 479 , K = 2
MRR_ID = 190 , K = 2
MRR_ID = 261 , K = 2
MRR_ID = 45 , K = 2
MRR_ID = 480 , K = 3
MRR_ID = 191 , K = 3
MRR_ID = 382 , K = 3
MRR_ID = 253 , K = 3
MRR_ID = 481 , K = 4
MRR_ID = 586 , K = 4
MRR_ID = 383 , K = 4
MRR_ID = 254 , K = 4
number recaptured
time, days

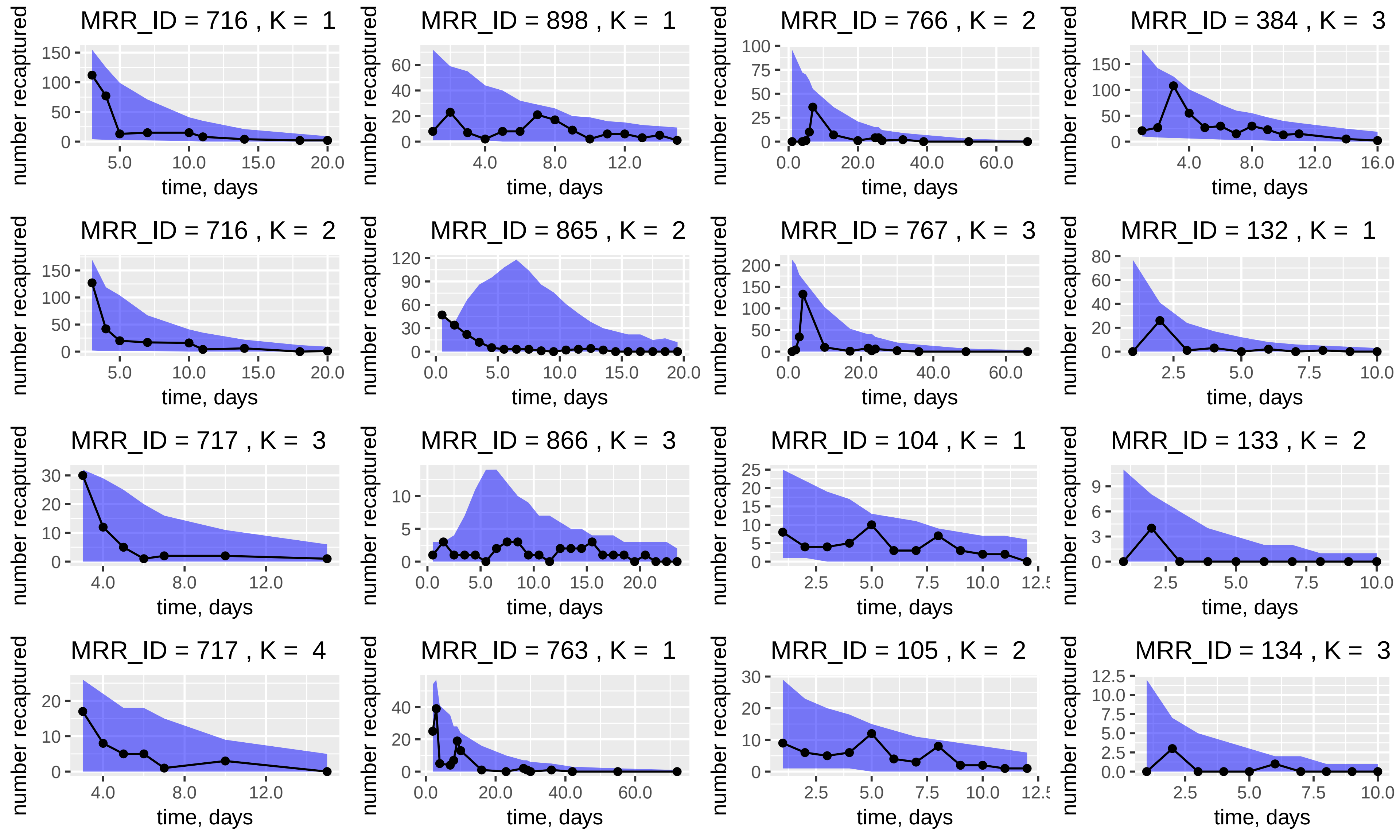
MRR_ID = 716 , K = 1
MRR_ID = 898 , K = 1
MRR_ID = 766 , K = 2
MRR_ID = 384 , K = 3
MRR_ID = 716 , K = 2
MRR_ID = 865 , K = 2
MRR_ID = 767 , K = 3
MRR_ID = 132 , K = 1
MRR_ID = 717 , K = 3
MRR_ID = 866 , K = 3
MRR_ID = 104 , K = 1
MRR_ID = 133 , K = 2
MRR_ID = 717 , K = 4
MRR_ID = 763 , K = 1
MRR_ID = 105 , K = 2
MRR_ID = 134 , K = 3
number recaptured
time, days

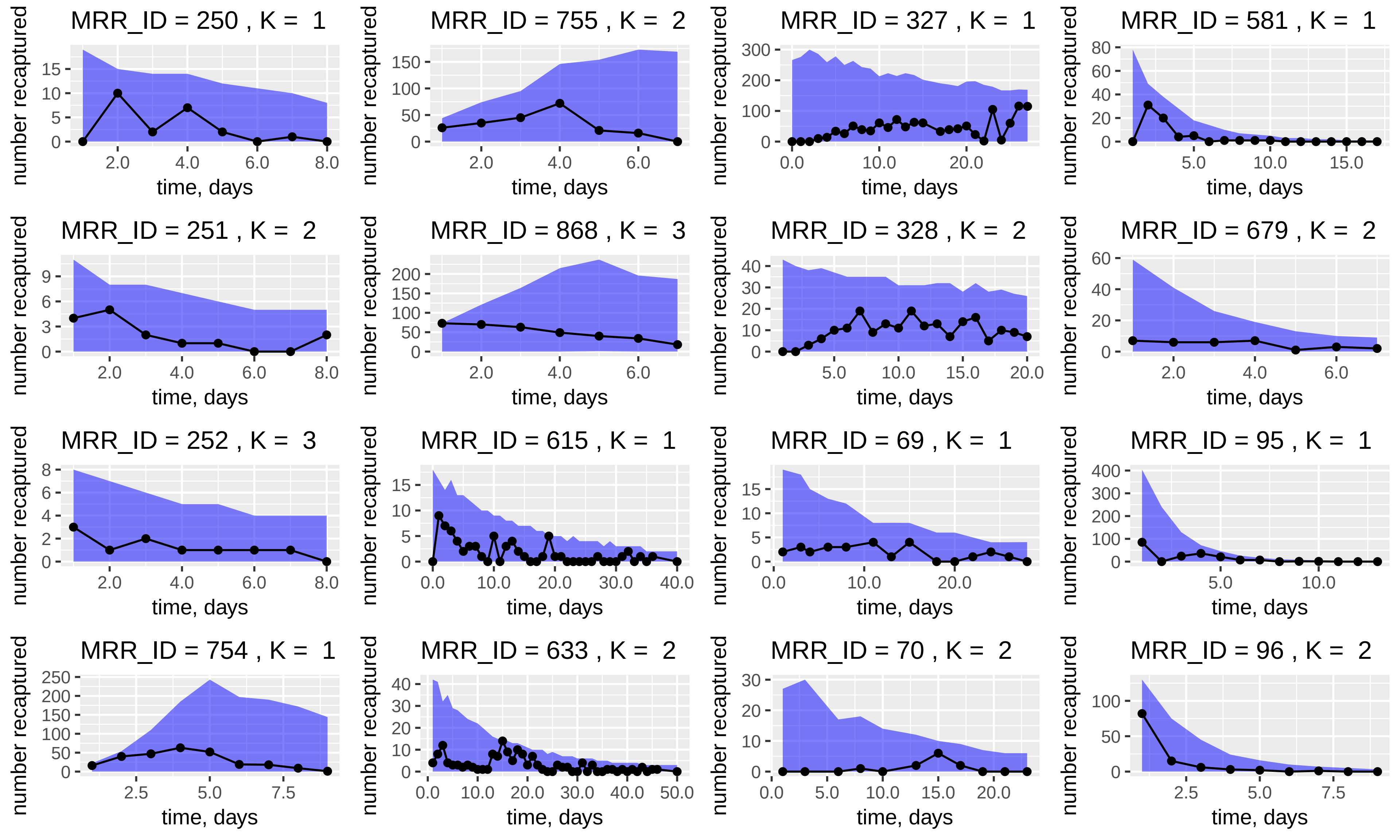
MRR_ID = 250 , K = 1
MRR_ID = 755 , K = 2
MRR_ID = 327 , K = 1
MRR_ID = 581 , K = 1
MRR_ID = 251 , K = 2
MRR_ID = 868 , K = 3
MRR_ID = 328 , K = 2
MRR_ID = 679 , K = 2
MRR_ID = 252 , K = 3
MRR_ID = 615 , K = 1
MRR_ID = 69 , K = 1
MRR_ID = 95 , K = 1
MRR_ID = 754 , K = 1
MRR_ID = 633 , K = 2
MRR_ID = 70 , K = 2
MRR_ID = 96 , K = 2
number recaptured
time, days

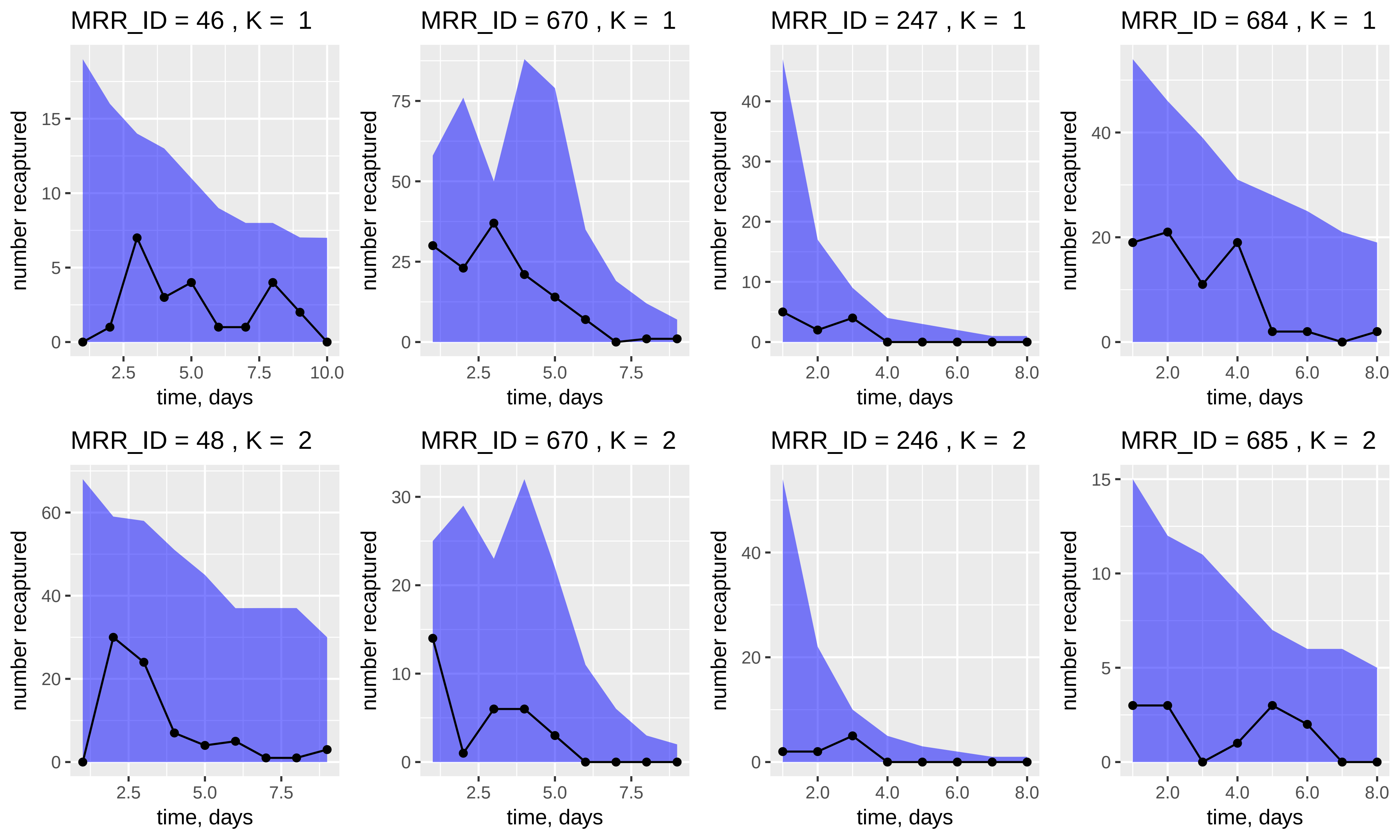
MRR_ID = 46 , K = 1
MRR_ID = 670 , K = 1
MRR_ID = 247 , K = 1
MRR_ID = 684 , K = 1
MRR_ID = 48 , K = 2
MRR_ID = 670 , K = 2
MRR_ID = 246 , K = 2
MRR_ID = 685 , K = 2
number recaptured
time, days
